# Chemical and structural investigation of the paroxetine-human serotonin transporter complex

**DOI:** 10.1101/2020.02.26.966895

**Authors:** Jonathan A. Coleman, Vikas Navratna, Daniele Antermite, Dongxue Yang, James A. Bull, Eric Gouaux

## Abstract

Antidepressants target the serotonin transporter (SERT) by inhibiting serotonin reuptake. Structural and biochemical studies aiming to understand the binding of small-molecules to conformationally dynamic transporters like SERT often require thermostabilizing mutations and antibodies to stabilize a specific conformation. Such modifications to SERT have led to questions about the relationships of these structures to the *bona fide* conformation and inhibitor binding poses of the wild-type transporter. To address these concerns, we characterized wild-type SERT with truncated N- and C-termini and thermostabilized variants of SERT bound with paroxetine using x-ray crystallography, single particle cryo-EM and biochemical techniques. Moreover, using a C–H functionalization approach to synthesize enantiopure analogues, we replaced the halide of the fluorophenyl group in paroxetine with either bromine or iodine. We then exploited the anomalous scattering of Br and I to define the pose of the respective paroxetine analog. These structures provide mutually consistent insights into how paroxetine and its analogs bind to the central substrate-binding site of SERT, stabilize the outward-open conformation, and inhibit serotonin transport.

## INTRODUCTION

Serotonin or 5-hydroxytryptamine (5-HT) is a chemical messenger which acts on cells throughout the human body, beginning in early development and throughout adulthood^1^. 5-HT acts as both a neurotransmitter and a hormone that regulates blood vessel constriction and intestinal motility^1^. In the central nervous system, 5-HT is released from presynaptic neurons where it diffuses across the synaptic space and binds to 5-HT receptors, promoting downstream signaling and activating postsynaptic neurons^2,3^. Thus, 5-HT is a master regulator of circuits, physiology and behavioral functions including the sleep/wake cycle, sexual interest, locomotion, thermoregulation, hunger, mood, and pain^1^. 5-HT is cleared from synapses and taken into presynaptic neurons by the serotonin transporter (SERT), thus terminating serotonergic signaling^2–4^. SERT resides in the plasma membrane of neurons and belongs to a family of neurotransmitter sodium symporters (NSSs) which also includes the dopamine (DAT) and norepinephrine transporters (NET)^2–4^. NSSs are twelve transmembrane spanning secondary active transporters which utilize sodium and chloride gradients to energize the transport of neurotransmitter across the membrane^4–6^ (Fig. 1a).

**Figure 1.**
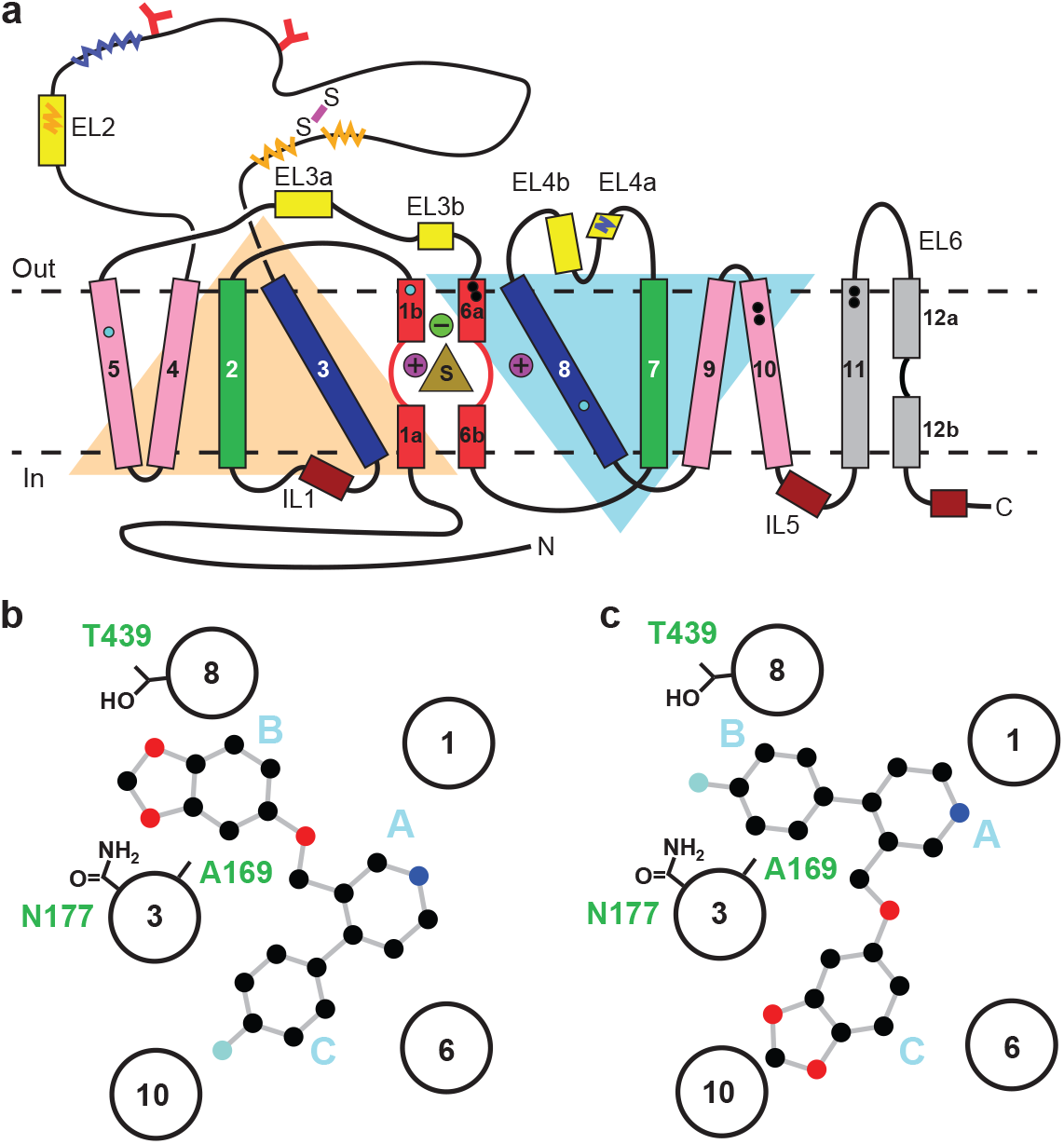
Topology of SERT. **a**, The substrate is bound at the central site (sand, triangle), near two sodium ions (purple, spheres +) and a chloride ion (green, sphere -). The light orange and light blue triangles depict pseudo two-fold symmetric helical repeats comprised of TM1-5 and 6-10, respectively. The disulfide bond (purple line) and *N*-linked glycosylation (red ‘Y’ shapes) in extracellular loop 2, along with sites of thermostable mutations (Tyr110Ala, TM1a; Ile291Ala, TM5; Thr439Ser, TM8) are also shown (cyan-filled circles). Structural elements involved in binding allosteric ligands are depicted as black-filled circles. Epitopes for the 8B6 and 15B8 Fab binding sites are in squiggly dark-blue and orange lines, respectively. **b**, Schematic of the ABC pose of paroxetine bound to the central binding site, derived from the previously determined x-ray structures^8,9^. The transmembrane helices are shown with circles and mutated residues in subsite B are in sticks. **c**, The ACB pose of paroxetine bound to the central binding site of SERT predicted by molecular dynamics simulations and mutagenesis^18,20^.

The function of NSSs is modulated by a spectrum of small-molecule drugs, thus in turn controlling the availability of neurotransmitter at synapses. Selective serotonin reuptake inhibitors (SSRIs) are a class of drugs which inhibit SERT and are used to treat major depression and anxiety disorders^7^. Using x-ray crystallography and cryo-EM, we have determined structures of thermostabilized variants of human SERT complexed with SSRIs, which together explain many of the common features and differences associated with SERT-SSRI interactions^8,9^. SSRIs are competitive inhibitors that bind with high-affinity and specificity to a central substrate-binding site in SERT, preventing 5-HT binding and arresting SERT in an outward-open conformation^2,3,9^.

The central site in NSSs is composed of three subsites: A, B, and C^10^ (Fig. 1b). In all NSS-ligand structures, the amine group of ligands resides in subsite A and interacts with a conserved Asp residue (Asp98 in SERT). The heterocyclic electronegative group of the ligand is positioned in subsite B^5^. For example, the alkoxyphenoxy groups of reboxetine and nisoxetine^11^ in *Drosophila* DAT (dDAT) structures, the halophenyl groups of cocaine analogs in dDAT and *S*-citalopram in SERT, and the catechol derivatives in DCP-dDAT and sertraline-SERT all occupy subsite B^8,9,12^. In addition to the central binding site, the activity of SERT and NSSs can also be modulated by small-molecules which bind to an allosteric site located in an extracellular vestibule, typically resulting in non-competitive inhibition of transport^9,13–15^.

Paroxetine is an SSRI which exhibits the highest known binding affinity for the central site of SERT (70.2 ± 0.6 pM) compared to any other currently prescribed antidepressants^16^. Despite its high affinity, paroxetine is frequently associated with serious side effects including infertility, birth defects, cognitive impairment, sexual dysfunction, weight gain, suicidality, and cardiovascular issues^17^. As a result, the mechanism of paroxetine binding to SERT has been studied extensively in order to design drugs with higher-specificity and less adverse side-effects. Despite these efforts, however, the binding pose of paroxetine remains a subject of debate^8,9,18–20^.

Paroxetine is composed of a secondary amine which resides in a piperidine ring, which in turn is connected to benzodioxol and fluorophenyl groups (Fig. 1b). X-ray structures of the SERT-paroxetine complex revealed that the piperidine ring binds to subsite A while the benzodioxol and fluorophenyl groups occupy subsite B and C in the central site, respectively^8,9^ (ABC pose). However, recent mutagenesis, molecular dynamics, and binding studies with paroxetine analogues suggest that paroxetine may occupy two distinct poses in which the benzodioxol and fluorophenyl groups reside in subsite B or C, depending on the rotameric position of Phe341 and the presence of the thermostabilizing mutation Thr439Ser^18,20^ (ACB pose, Fig. 1c). Paroxetine is also thought to interact with the allosteric site of SERT, albeit with low-affinity^15^. We have, however, been unable to visualize paroxetine binding at the allosteric site using structural methods. Our x-ray maps, by contrast, resolve a density feature at the allosteric site which instead resembles a molecule of detergent^9^.

To resolve the ambiguity of paroxetine binding poses at the central binding site, we turned to paroxetine derivatives whereby the 4-fluoro group is substituted with either a bromine or an iodine group. Using transport and binding assays, anomalous x-ray diffraction, and cryo-EM, we have examined the binding poses of these paroxetine analogs and their interactions at the central site. Our data shows that paroxetine binds in the ABC pose. Thus, these structures provide key insights into the recognition of high-affinity inhibitors by SERT and the rational design of new small-molecule therapeutics.

## RESULTS

To provide a robust molecular basis for the interaction of paroxetine (**1**) with SERT, we devised synthetic routes for two derivatives of paroxetine where the 4-fluoro moiety is substituted with either bromo (Br-paroxetine, **2**) or iodo (I-paroxetine, **3**) groups (Fig. 2a,b). We envisaged the use of a C–H functionalization strategy to access enantiopure hydroxymethyl intermediates **I**, from readily available *N*-Boc (*R*)-nipecotic acid **4** (Fig. 2b, Supplementary file 1). Transition metal-catalyzed C–H functionalization can promote the reaction of unactivated C(sp^3^)–H bonds with the aid of a directing group^21–26^. Here, C–H functionalization enabled installation of the appropriate aryl group on the pre-existing piperidine ring^27^, providing an attractive and short route to vary this functionality with inherent control of enantiomeric excess. In contrast, common methods for (–)-paroxetine synthesis can require the aromatic substituent to be introduced before stereoselective steps or ring construction, reducing flexibility of the process^20,28–34^.

**Figure 2.**
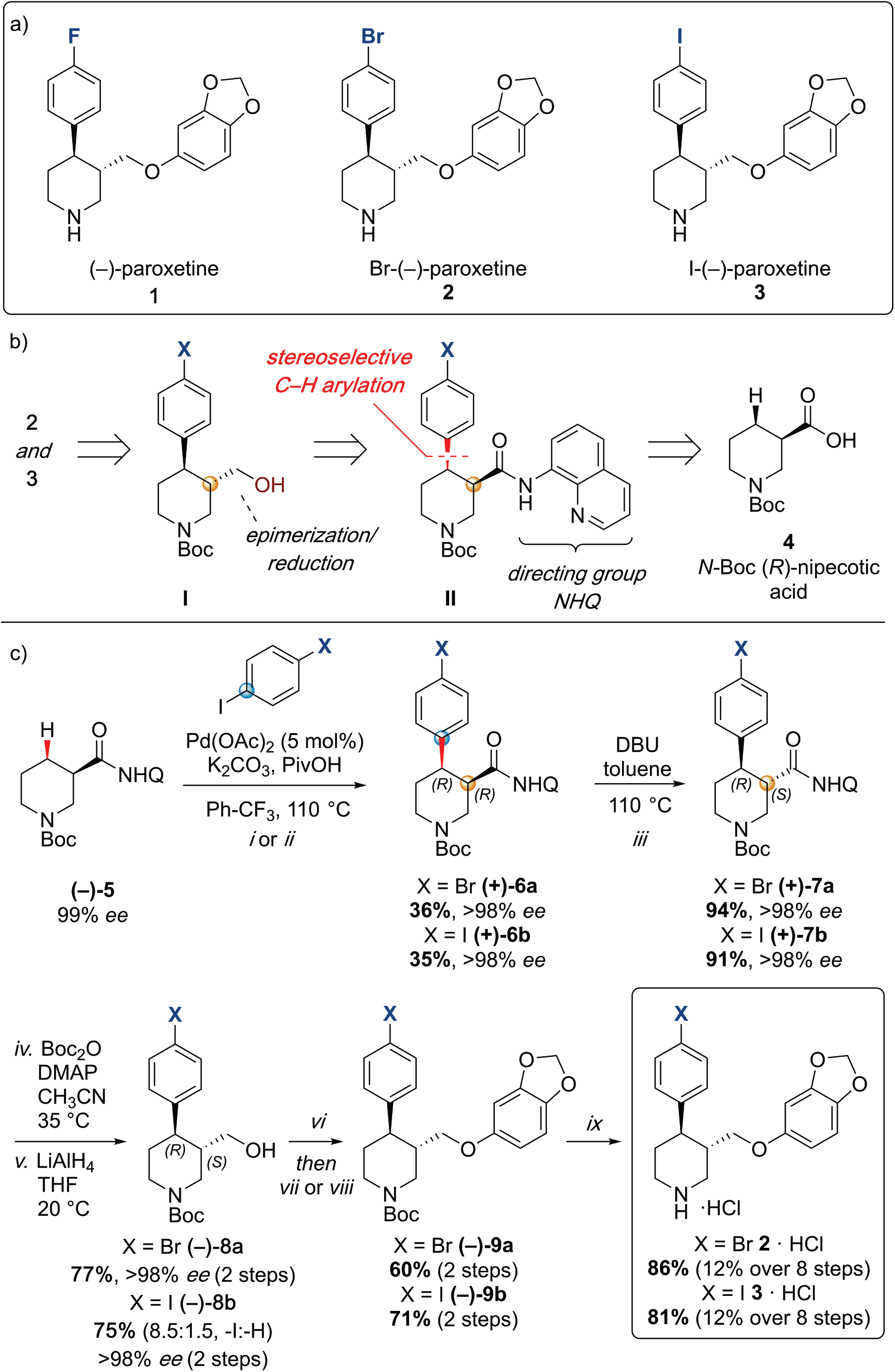
Synthesis of paroxetine analogues. **a**, Structures of (–)-paroxetine **(1)** and the targeted Br-**(2)** and I-analogues **(3)**. **b**, Retrosynthetic analysis of Br- and I-(–)-paroxetine. **c**, Synthesis of Br- and I-(–)-paroxetine **2** and **3**. Q = 8-quinolinyl-. Reaction conditions: *i*) X = Br: **(–)^−5^** (4.0 mmol), 4-bromo iodobenzene (3 equiv), Pd(OAc)_2_ (5 mol %), K_2_CO_3_ (1 equiv), PivOH (1 equiv), Ph-CF_3_ (2 mL, 2 M), 110 °C, 18 h; *ii*) X = I: **(–)-5** (4.0 mmol), 1,4-diiodobenzene (4 equiv), Pd(OAc)_2_ (5 mol %), K_2_CO_3_ (1 equiv), PivOH (1 equiv), Ph-CF_3_ (2 mL, 2 M), 110 °C, 18 h; *iii*) DBU (3 equiv), toluene (1 M), 110 °C, 24 h; *iv*) Boc_2_O (4 equiv), DMAP (20 mol %), CH_3_CN (0.5 M), 35 °C, 22 h; *v*) LiAlH_4_ (2 equiv), THF, 20 °C, 0.5 h; *vi*) MsCl (1.3 equiv), Et_3_N (1.4 equiv), CH_2_Cl_2_, 0 to 25 °C, 2 h; *vii*) X = Br: sesamol (1.6 equiv), NaH (1.7 equiv), THF, 0 to 70 °C, 18 h; *viii*) X = I: sesamol (2.0 equiv), NaH (2.2 equiv), DMF, 0 to 90 °C, 20 h; *ix*) 4 N HCl in dioxane (10 equiv), 0 to 25 °C, 18 h.

Our synthesis commenced with the C–H arylation of piperidine **(–)-5** bearing Daugulis’ aminoquinoline amide directing group^35^ at C(3). Adapting our reported method^27^, Pd-catalyzed C– H functionalization was achieved in moderate yields using 4-bromoiodobenzene or 1,4-diiodobenzene in excess to prevent bis-functionalization, with palladium acetate, K_2_CO_3_ and pivalic acid (Fig. 2c). The *cis-*arylated derivatives **(+)-6a** and **(+)-6b** were obtained with >98% *ee* and complete C(4) selectivity. Minor enantiopure *trans*-functionalized products, formed via a *trans*-palladacycle^27^, were also isolated (Supplementary File 1). Subsequent treatment with 1,8-diazabicyclo(5.4.0)undec-7-ene (DBU) gave complete C(3)-epimerization affording **(+)-7a** and **(+)-7b** with the desired *trans-*stereochemistry in 94% and 91% yields. The aminoquinoline group was removed through telescoped amide activation and reduction with LiAlH_4_ at 20 °C to give enantiopure hydroxymethyl intermediates **(–)-8a** and **(–)-8b** in 77% and 75% yield. Mesylation and nucleophilic substitution with sesamol formed ether derivatives **(–)-9a** and **(–)-9b**, which were deprotected to give Br- and I-paroxetine **2** and **3**. An overall yield of 12% over 8 steps from commercial material was obtained in both cases. At each stage the identity of the products and purity was established by acquiring ^1^H and ^13^C nuclear magnetic resonance spectra, IR spectra, and by high resolution mass spectrometry. Enantiopurity was assessed by high-performance liquid chromatography (HPLC) with reference to racemic or scalemic samples (Supplementary File 1).

We also employed several SERT variants and the 8B6 Fab in the biochemical and structural studies described here. The wild-type SERT construct used in transport experiments contains the full-length SERT sequence fused to a C-terminal GFP tag. The ts2-active variant contains two thermostabilizing mutations (Ile291Ala, Thr439Ser) which allows for purification of the apo transporter for binding studies and has kinetics of 5-HT transport (K_m_: 4.5 ± 0.6 μM, V_max_: 21 ± 5 pmol min^−1^) that are in a similar range as wild-type (K_m_: 1.9 ± 0.3 μM, V_max_: 23 ± 1 pmol min^−1^)^9,36^. The ts2-inactive variant (Tyr110Ala, Ile291Ala)^8^, by contrast, is unable to transport 5-HT but can be crystallized due to the stabilizing Tyr110Ala mutation^36^ and binds SSRIs with high-affinity. The ΔN72, ΔC13 SERT variant used for cryo-EM is otherwise wild-type SERT which has been truncated at the N- and C-termini and yet retains transport and ligand binding activities^37^. Finally, the recombinant 8B6 Fab^9,38^ was used to produce SERT-Fab complexes which were studied by x-ray crystallography and cryo-EM.

We began by accessing the functional effects of paroxetine, Br-paroxetine, and I-paroxetine on SERT activity by measuring their inhibition of 5-HT transport and *S*-citalopram competition binding. We assayed the ability of the Br- and I-paroxetine derivatives to inhibit 5-HT transport in HEK293 cells expressing wild-type SERT, observing that upon substituting the 4-fluoro group with 4-bromo or 4-iodo groups, the potency of inhibition of 5-HT transport in wild-type SERT decreased significantly from 4 ± 1 for paroxetine to 40 ± 20 for Br-paroxetine and 180 ± 70 nM for I-paroxetine (Fig. 3a, Table 1). Next, we measured the binding of paroxetine, Br-paroxetine, and I-paroxetine to ts2-active and ts2-inactive SERT using *S*-citalopram competition binding assays, finding that the SERT variants employed in this study exhibited high-affinity for paroxetine and its derivatives (Table 2). A decrease in the binding affinity upon substituting the 4-fluoro group of paroxetine with 4-bromo or 4-iodo groups was observed in the competition binding assays. However, the difference in the binding affinities between paroxetine variants measured by the competition binding assay was not as pronounced as the difference in the inhibition potencies observed in the 5-HT transport assays (Table 2). For example, the ts2-inactive (Tyr110Ala, Ile291Ala) variant employed in the previous^8^ and present x-ray studies exhibited a K_i_ of 0.17 ± 0.02 nM for paroxetine, 0.94 ± 0.01 nM for Br-paroxetine, and a further decrease in affinity to I-paroxetine (2.3 ± 0.1 nM, Fig. 3b, Table 2).

**Table 1.** Inhibition of 5-HT transport by paroxetine and its derivatives.

| Ligand | IC<sub>50</sub> (nM) |
| --- | --- |
| Paroxetine | 4 ± 1 |
| Br-paroxetine | 40 ± 20 |
| I-paroxetine | 180 ± 70 |

**Table 2.**
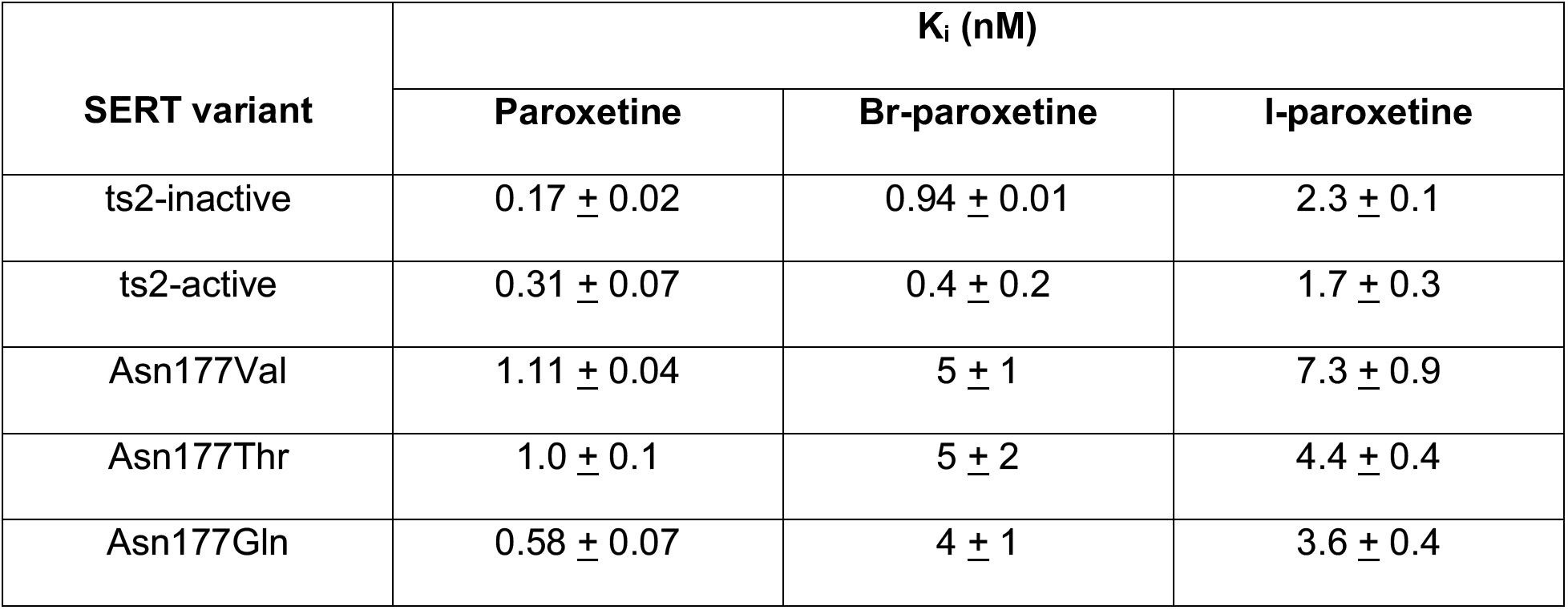
Binding of paroxetine and its derivatives to SERT variants used in this study.

**Table 3.** Cryo-EM data collection, refinement and validation statistics^a^.

|  | #1<br>(EMDB-21368)<br>(PDB 6VRH)<br>(EMPIAR-XXXX) | #2<br>(EMDB-21369)<br>(PDB 6VRK) | #3<br>(EMDB-21370)<br>(PDB 6VRL) |
| --- | --- | --- | --- |
| **Data collection and processing** |  |  |  |
| Magnification | 77,160 | 77,160 | 77,160 |
| Voltage (kV) | 300 | 300 | 300 |
| Electron exposure (e-/Å<sup>2</sup>) | 54-60 | 54 | 54 |
| Defocus range (μm) | -0.8 to -2.2 | -0.8 to -2.2 | -0.8 to -2.2 |
| Pixel size (Å) | 0.648 | 0.648 | 0.648 |
| Symmetry imposed | C1 | C1 | C1 |
| Initial particle images (no.) | 4,147,084 | 4,545,318 | 4,470,768 |
| Final particle images (no.) | 420,373 | 503,993 | 414,091 |
| Map resolution (Å) | 3.3 | 4.1 | 3.8 |
| FSC threshold | 0.143 | 0.143 | 0.143 |
| Map resolution range (Å)<sup>b</sup> | 4.25-3.25 | 5.75-3.75 | 5.50-3.50 |
| **Refinement** |  |  |  |
| Initial model used (PDB code) | 6AWN | 6VRH | 6VRH |
| Initial model CC | 0.64 | 0.70 | 0.71 |
| Model resolution (Å)<sup>c</sup> | 3.7 | 4.3 | 4.1 |
| FSC threshold | 0.5 | 0.5 | 0.5 |
| Model resolution range (Å) | 25.9-3.3 | 33.0-4.1 | 29.6-4.2 |
| Map sharpening *B* factor (Å<sup>2</sup>) | -85 | -174 | -161 |
| **Model composition** |  |  |  |
| Non-hydrogen atoms | 6143 | 6142 | 6142 |
| Protein residues | 764 | 764 | 764 |
| Ligands (atoms) | 254 | 254 | 254 |
| ***B* factors (Å<sup>2</sup>)** |  |  |  |
| Protein | 138 | 138 | 122 |
| Ligand | 129 | 113 | 112 |
| **R.m.s. deviations** |  |  |  |
| Bond lengths (Å) | 0.002 | 0.002 | 0.002 |
| Bond angles (°) | 0.48 | 0.59 | 0.54 |
| **Validation** |  |  |  |
| Refined model CC | 0.73 | 0.74 | 0.75 |
| MolProbity score | 1.86 | 1.96 | 1.88 |
| Clashscore | 9.67 | 10.26 | 10.59 |
| Poor rotamers (%) | 0 | 0 | 0.00 |
| **Ramachandran plot** |  |  |  |
| Favored (%) | 94.84 | 93.54 | 95.12 |
| Allowed (%) | 5.16 | 6.46 | 4.88 |
| Disallowed (%) | 0 | 0 | 0 |

<sup>a</sup>Data set #1 is the paroxetine reconstruction, #2 is Br-paroxetine, #3 l-paroxetine.<sup>b</sup>Local resolution range.<sup>c</sup>Resolution at which FSC between map and model is 0.5.

**Table 4.** X-ray data collection statistics.

|  | Br-paroxetine | I-paroxetine |
| --- | --- | --- |
| **Data collection** |  |  |
| Space group | C222<sub>1</sub> | C222<sub>1</sub> |
| Cell dimensions<br>*a*, *b*, *c* (Å) | 128.0, 161.9, 139.7 | 127.7, 161.9, 140.8 |
| $\alpha$, $\beta$, $\gamma$ (°) | 90, 90, 90 | 90, 90, 90 |
| Resolution (Å) | 20.45-4.69 (4.82-4.69)* | 25.98-6.12 (6.30-6.12)* |
| *R*<sub>merge</sub> | 13.60 (339.3) | 7.9 (292.9) |
| *I* / σ*I* | 5.51 (0.49) | 5.01 (0.32) |
| CC<sub>1/2</sub> | 99.9 (16.5) | 99.8 (20.0) |
| Completeness (%) | 99.2 (100.0) | 92.6 (89.7) |
| Redundancy | 6.8 (6.2) | 1.8 (1.7) |

\*Values in parentheses are for highest-resolution shell.

**Figure 3.**
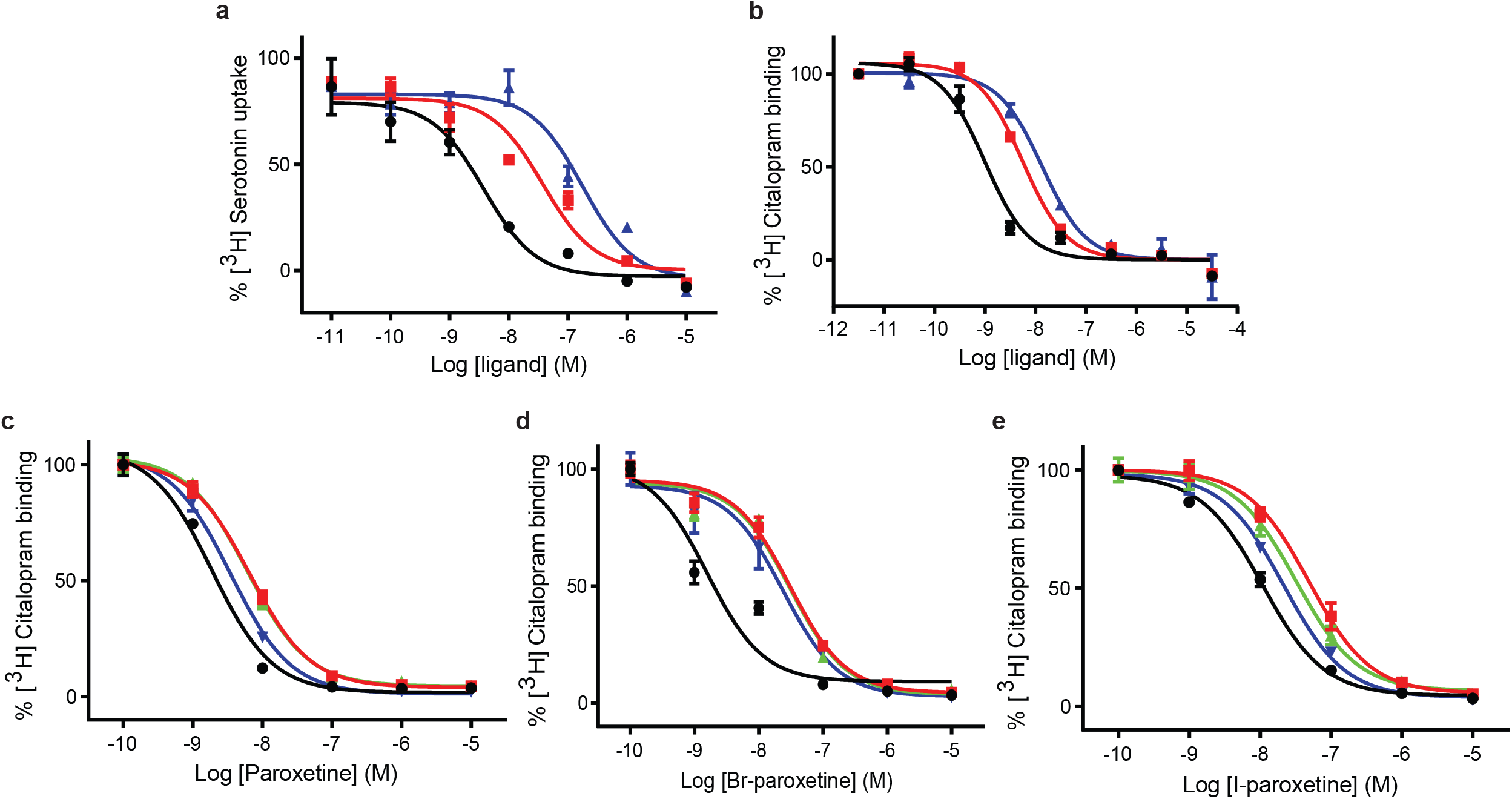
Inhibition of [^3^H]5-HT transport and [^3^H]citalopram binding by paroxetine and the Br- and I-derivatives. **a**, 5-HT-transport of wild-type SERT and its inhibition by paroxetine, Br-, and I-paroxetine. **b**, Competition binding of paroxetine and its derivatives to ts2-inactive SERT. In panels a and b, paroxetine, Br-paroxetine, and I-paroxetine curves are shown as black, red, and blue lines, respectively. **c**, Competition binding of paroxetine to ts2-active (black), Asn177Val (red), Asn177Thr (green), and Asn177Gln (blue). **d**, Competition binding of Br-paroxetine. **e**, Competition binding of I-paroxetine. The values associated with these experiments are reported in Table 1 and 2.

In the x-ray structures of SERT, the benzodioxol group of paroxetine in the ABC pose is found in subsite B^8,9^. A recent study suggested that binding affinity and potency to inhibit the transport of Br-paroxetine was only moderately affected upon mutating a non-conserved residue Ala169 to Asp in subsite B of SERT^20^ (Fig. 1b). We recently also identified a conserved residue, Asn177 in the subsite B, which upon mutation exhibited differential effects on the inhibitory potency of ibogaine and noribogaine^37^. To further probe the role of Asn177 in subsite B, we studied the binding of paroxetine and its derivatives to selected Asn177 mutants designed in the ts2-active background (Fig. 1b). We observed that the affinity of paroxetine to ts2-active SERT decreased by 2-fold when Asn177 is substituted with small non-polar or polar residues such as valine and threonine, while only a negligible change in K_i_ was observed for glutamine (Asn177Gln) (Fig. 3c). In the case of Br-paroxetine, the Asn177 variants (K_i_ between 4-5 nM) display up to a 10-fold decrease in K_i_ when compared with ts2-active SERT (0.4 ± 0.2 nM) (Fig. 3d, Table 2). A similar behavior was also observed for I-paroxetine, with ts2-active exhibiting a K_i_ of 1.7 ± 0.3 nM and the mutants a K_i_ of 4-7 nM, with the valine and glutamine substitutions exhibiting the highest and lowest Ki values respectively (Fig. 3e, Table 2).

To define the binding poses of paroxetine and its analogues to SERT, we solved the structures of the ΔN72, ΔC13 and the ts2-inactive SERT variants complexed with Br- and I-paroxetine using single particle cryo-EM and x-ray crystallography (Fig. 4, Supplementary Figs. 2,3). We began by collecting cryo-EM data sets for ΔN72, ΔC13 SERT-8B6 Fab complexes with each ligand. The TM densities in all three reconstructions were well-defined and contiguous allowing for clear positioning of the main chain in an outward-open conformation (Supplementary Figs. 4,5). Large aromatic side-chains were well-resolved for all three complexes, also suggesting that the aromatic moieties of paroxetine and its analogues could be identified and positioned in our cryo-EM maps. In addition, the particle distribution and orientations of SERT-Fab complexes in presence of Br- and I-paroxetine were similar to paroxetine, allowing for uniform comparison between the maps.

**Figure 4.**
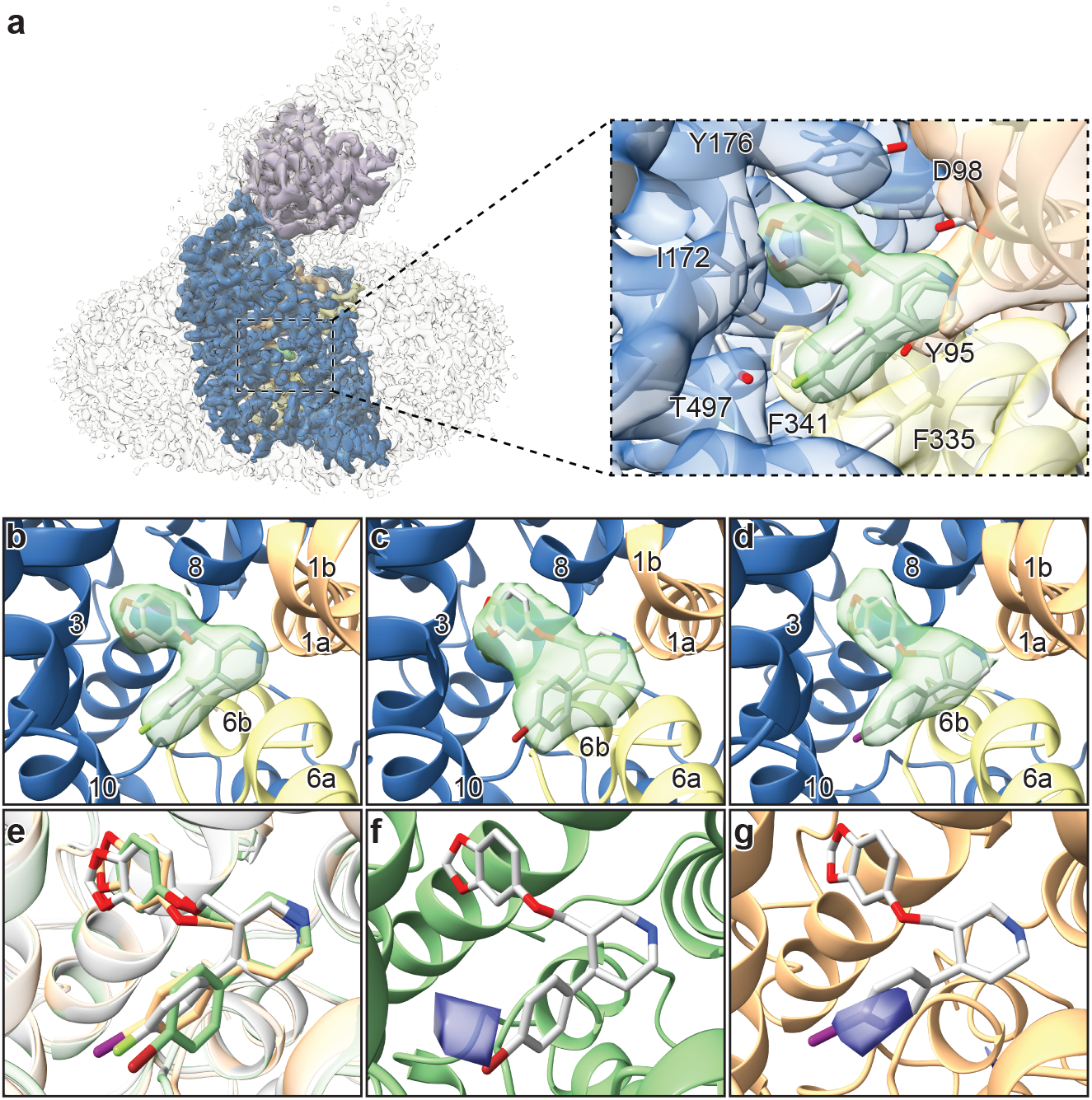
Structures of SERT-paroxetine complexes. **a**, Cryo-EM reconstruction of SERT bound to paroxetine where the shape of the SERT-8B6 Fab complex and detergent micelle is shown in transparent light grey. The density of SERT is shown in dark blue with TM1 and TM6 colored in orange and yellow, respectively, and the density for paroxetine in green. The variable domain of the 8B6 Fab is colored in purple. Inset shows the density features at the central site of paroxetine. **b**, Density feature at the central site of paroxetine. **c**, Density feature at the central site of Br-paroxetine. **d**, Density feature at the central site of I-paroxetine. **e**, Comparison of the binding poses of paroxetine (grey), Br-paroxetine (green), and I-paroxetine (orange). **f**, Anomalous difference electron density (blue) derived from Br-paroxetine, contoured at 4σ. **g**, Anomalous difference electron density (blue) derived from I-paroxetine, contoured at 4σ.

**Figure 5.**
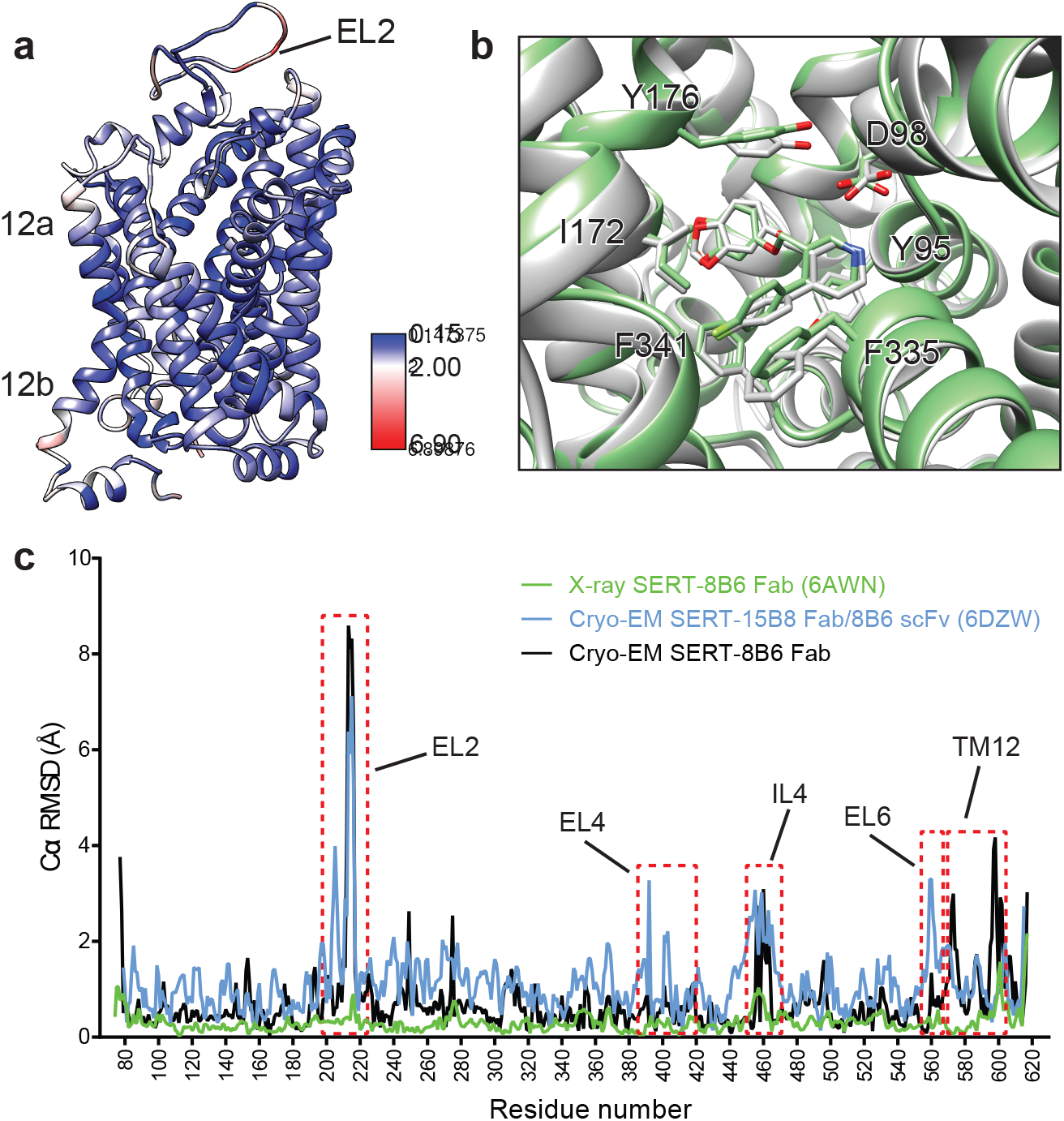
Comparison of the x-ray and cryo-EM structures of the SERT-paroxetine complex. **a**, Superposition of the x-ray ts3-SERT-8B6 paroxetine structure (PDB: 5I6X) with the SERT-8B6 paroxetine complex determined by cryo-EM. The root-mean-square-deviations (RMSD) for Cα positions were plotted onto the cryo-EM SERT-8B6 paroxetine structure. **b**, Comparison of the central binding site of the x-ray (grey) and cryo-EM (green) paroxetine structures. **c**, The structure of the ts2-inactive SERT-8B6 scFv/15B8 Fab paroxetine (cryo-EM, 6DZW), ts2-inactive SERT-8B6 Fab paroxetine (x-ray, 6AWN), and the SERT-8B6 paroxetine (cryo-EM, this work) complexes were superposed onto the ts3 SERT-8B6 paroxetine complex (x-ray, 5I6X) as a reference. The RMSD for Cα positions were calculated for each structure in comparison with the reference. Regions with RMSD > 3.0 Å are shown boxed in red.

The ~3.3 Å resolution map of the ΔN72, ΔC13 SERT-8B6 paroxetine complex allowed us to model the inhibitor unambiguously at the central site (Fig. 4a). The resolution of the Br- and I-paroxetine complexes was comparatively lower at ~4.1 Å and ~3.8 Å, respectively (Supplementary Fig. 5). Nevertheless, the ligands could also be modeled into the density at the central site with a correlation coefficient (CC) of 0.75 and 0.77, respectively (Figs. 4b-e). All three ligands best fit their respective density features in the ABC pose, as previously demonstrated in the x-ray structure of ts2-inactive and ts3 SERT^8,9^ (PDB ID: 6AWN, 5I6X). To probe the possibility of an ACB pose of paroxetine, we also flexibly modeled paroxetine in the ACB pose at the central site followed by real space refinement, finding the ACB pose could be modeled with a CC of 0.70 compared with 0.84 for the ABC pose (Supplementary Figs. 6a,b). We also compared the reconstructed complexes by calculating difference maps, attempting to identify features associated with the scattering of bromine and iodine. However, the resulting difference maps did not contain any interpretable difference densities and thus did not further assist in ligand modeling.

We then explored the binding pose of paroxetine by growing crystals and collecting x-ray data of the ts2-inactive SERT-8B6 Fab complex with Br- and I-paroxetine. Anomalous difference maps calculated from the previously determined ts2-inactive paroxetine structure (PDB ID: 6AWN) after refinement, showed clear densities for Br- and I-atoms of the paroxetine derivatives in subsite C (Figs. 4f,g). No detectable anomalous peaks were observed in either subsite B or in the allosteric site and there were no other peaks in any other location above 3σ, suggesting that under these conditions, Br-paroxetine and I-paroxetine do not bind substantially in the ACB orientation or to the allosteric site.

We next compared the cryo-EM structure of the SERT-paroxetine complex to the x-ray structure of the ts3 SERT paroxetine complex. Overall comparison of the transporter revealed only minor variation between structures solved by each method, with a Cα root-mean-square-deviation (RMSD) of 0.68 Å. The most significant differences between the cryo-EM and the x-ray structures were found at the extracellular and intracellular sites of TM12 and also in EL2, while the core of the transporter (TM1-10) was largely unchanged (Fig. 5a). These changes can largely be explained on the basis of a crystal packing interface formed by TM12 and a highly flexible EL2 that is bound to the 8B6 Fab. We also compared central site residues involved in paroxetine binding, finding that the best fit to the cryo-EM density revealed only minor differences in the side-chains of Asp98, Tyr176, and Phe335 when compared to the x-ray structure (all atom RMSD: 0.91 Å) (Fig. 5b). Finally, we compared the cryo-EM structures of the SERT 15B8 Fab/8B6 scFv paroxetine complex (PDB: 6DZW) to the SERT 8B6 Fab paroxetine complex to understand if these antibodies induce changes in transporter structure. Here we found that the most significant differences occurred in the extracellular domain and involved localized regions of EL2 and EL4 that interact with the antibody (Fig. 5c). The transporter core was largely unchanged, with the only other significant differences being found in EL6, TM12, and IL4.

## DISCUSSION

The binding of paroxetine to SERT has been extensively debated^8,9,18–20^. The first x-ray structure of the ts3-SERT variant demonstrated that the binding pose is such that the piperidine, benzodioxol, and fluorophenyl groups occupy subsites A, B, and C respectively, in the ABC pose^9^ (Fig. 1b). Competition binding experiments using a variant of SERT containing a central binding site that has been genetically engineered to possess photo-cross-linking amino acids favor the ABC pose of paroxetine. These cross-linking experiments demonstrated that paroxetine binds in a fashion that is similar to the crystal structure^8,9^, where the fluorophenyl group is in proximity to Val501^39^. However, computational docking experiments using wild type SERT showed that the position of benzodioxol and fluorophenyl groups of paroxetine is ‘flipped’, with paroxetine occupying an ACB pose^18–20^ (Fig. 1c). In this study, the authors hypothesized that the difference could be because of the crystallization conditions and thermostabilizing mutations. One of the thermostabilizing mutations in ts3-SERT, Thr439Ser, is near the central binding site and Thr439 participates in a hydrogen bonding network in subsite B that, in turn, includes the dioxol group of paroxetine.

To probe the role of the Thr439Ser mutation in modulating the binding pose of paroxetine, we solved the x-ray structure of ts2-inactive (Tyr110Ala, Ile291Ala) SERT, wherein the residue at position 439 was the wild-type threonine. Paroxetine occupies the ABC pose in the x-ray structure of ts2-inactive SERT^8^. MD simulations of ts2-inactive SERT suggested that the Thr439Ser mutation weakens the Na2 site. Furthermore, MD simulations and binding and uptake kinetics experiments using wild-type SERT in presence of paroxetine and a variant of paroxetine where in the 4-fluoro group is substituted with 4-bromo group suggested that the paroxetine binding pose in SERT could be ambiguous because of the pseudo symmetry of the paroxetine molecule. It was noted that paroxetine could occupy both ABC and ACB poses with almost equivalent preference. Upon substituting the 4-fluoro with a bulkier 4-bromo group, the ABC pose was favored^18,20^.

Structural studies of SERT in complex with paroxetine and its analogues were thus required to resolve the uncertainty in paroxetine binding pose at the central site. Previously, we had demonstrated that cryo-EM can be used to define the position of ligands at the central site of SERT^37^. Here, we employed a similar methodology using the ΔN72, ΔC13 SERT variant complexed with 8B6 Fab to study binding of paroxetine at the central site. The density feature of paroxetine in the cryo-EM map at ~3.3 Å clearly resolved the larger benzodioxol and smaller fluorophenyl groups in subsite B and C respectively (Fig. 4b). Though this reconstruction suggests that paroxetine binds in the ABC pose, we also considered the possibility that the inhibitor density feature may represent an average of the ABC and ACB poses. We expected that if Br- and I-paroxetine were suitable surrogates for paroxetine, their binding pose would be unaffected by their reduced electronegativity and the size of the halogenated groups and therefore that they would also be associated with a comparable density feature at this site, as demonstrated by our cryo-EM maps. To further explore if there was a fraction of Br- or I-paroxetine in the ACB pose, we examined the position of the Br- or I-atoms at the central site by x-ray crystallography. If Br- and I-paroxetine were to bind in both the ABC or ACB poses, we expected to observe two anomalous peaks in our x-ray maps in subsites B and C; for both ligands, however, only a single detectable peak was observed in subsite C (Fig. 4f,g). Thus, our direct biophysical observations reveal that the dominant binding mode of paroxetine is the ABC pose and that, if the ACB pose is present at all, it represents a minor fraction of binding poses.

Paroxetine is stabilized in the ABC pose at the central binding site by aromatic, ionic, non-ionic, hydrogen bonding, and cation-π interactions. The ACB pose would alter or disrupt most of these interactions, resulting in clashes in subsites A and C as earlier reported^8^. In subsite A, the amine of the piperidine ring of paroxetine binds with Asp98 (3.5 Å) and also makes a cation-π interaction with Tyr95 (Fig. 4a). The benzodioxol group of paroxetine, a catechol-like entity, occupies a position in subsite B which is similar to the binding of catechol derivative groups of sertraline and 3,4-dichlorophenethylamine in SERT^8^ and dDAT^12^ structures, respectively. In subsite B, the ring of Tyr176 makes an aromatic interaction with the benzodioxol while the hydrogen-bonding network in subsite B formed by Asn177, Thr439, backbone carbonyl oxygens, and amides are likely responsible for stabilization of the dioxol. The side-chain of Ile172 inserts between the benzodioxol and fluorophenyl, while the rings of Phe341 and Phe335 stack on either side of the fluorophenyl, ‘sandwiching’ it within subsite C. The fluorine, bromine, or iodine of paroxetine and its analogues reside adjacent to the side-chain of Thr497 (4.0 Å), which may act to stabilize these groups through hydrogen bonding (Fig. 4a). The larger atomic radius of the halogens, the longer length of the carbon-halogen bond, and the difference in electronegativity of bromine (radius: 1.85 Å, bond-length: 1.92 Å, electronegativity: 2.96) and iodine (radius: 1.98 Å, bond-length: 2.14 Å, electronegativity: 2.66) relative to fluorine (radius: 1.47 Å, bond-length: 1.35 Å, electronegativity: 3.98) explain why the fluorine is better accommodated in subsite C and why Br-paroxetine and I-paroxetine bind with weaker affinity.

We also explored the effect of conservative and non-conservative mutations in subsite B of SERT at Asn177 (Fig. 3). Asn177 participates in a hydrogen-bond network with the hydroxyl group of noribogaine and with the dioxol of paroxetine. However, this network of interactions is also important for binding halogenated inhibitors in subsite B, as in the case for *S*-citalopram, fluvoxamine, and sertraline. All of the mutants that we tested at Asn177 resulted in a loss of binding affinity to paroxetine and its analogues. Furthermore, the Ala169Asp mutation in subsite B^20^ (Fig. 1b,c) also reduced paroxetine inhibition and binding, likely also disrupting these interactions. Although the effects were less severe when compared to paroxetine, Br-paroxetine binding and inhibition was also reduced for Ala169Asp^20^. Thus, these mutations highlight the importance of subsite B interactions in paroxetine binding but they cannot be used to demonstrate the inhibitor pose because, in the ABC or ACB poses, either the dioxol or fluorine of paroxetine could act as a hydrogen-bond acceptor in subsite B.

Using a combination of chemical biology, cryo-EM, and x-ray crystallography we demonstrate that the SSRI paroxetine occupies the ABC pose at the central site, where it is involved in numerous interactions. Hence, our studies of the mechanism of paroxetine binding to SERT provide a robust framework for the design of new highly specific small-molecule SERT inhibitors.

## MATERIALS AND METHODS

### SERT expression and purification

The human SERT constructs used in this study were the wild-type, the N- and C-terminally truncated wild-type (ΔN72, ΔC13), ts2-inactive (Tyr110A, Ile291Ala), and ts2-active (Ile291Ala, Thr439Ser)^8,9,36–38^ proteins. The Asn177 mutants were generated in the ts2-active background. The expression and purification of SERT was carried out as previously described with minor modifications^8,9,37,38^, as described below. All SERT constructs were expressed as C-terminal GFP fusion using baculovirus-mediated transduction of HEK293S GnTI^−^ cells. Cells were solubilized in 20 mM Tris pH 8 with 150 mM NaCl, containing 20 mM n-dodecyl-β-D-maltoside (DDM) and 2.5 mM cholesteryl hemisuccinate (CHS), followed by purification using Strep-Tactin affinity chromatography in 20 mM Tris pH 8 with 100 mM NaCl (TBS), 1 mM DDM, and 0.2 mM CHS.

For cryo-EM of the ΔN72,ΔC13 SERT, 1 mM 5-HT was added during solubilization and affinity purification to stabilize SERT. GFP was cleaved from SERT by digestion with thrombin and the SERT-8B6 complex was made as described in the previous paragraph. The complex was separated from free Fab and GFP by SEC in TBS containing 1 mM DDM and 0.2 mM CHS, and the peak fractions were concentrated to 4 mg/ml followed by addition of either 200 μM paroxetine, Br-paroxetine or I-paroxetine.

For crystallization, no ligands were added during purification of ts2-inactive SERT, and 5% glycerol and 25 μM lipid (1-palmitoyl-2-oleoyl-sn-glycero-3-phosphocholine, 1-palmitoyl-2-oleoyl-sn-glycero-3-phosphoethanolamine, and 1-palmitoyl-2-oleoyl-sn-glycero-3-phosphoglycerol at a molar ratio of 1:1:1) were included in all the purification buffers. Following affinity purification, the fusion protein was digested by thrombin and EndoH and combined with recombinant 8B6 Fab at a molar ratio of 1:1.2. The SERT-8B6 complex was isolated by size-exclusion chromatography (SEC) on a Superdex 200 column in TBS containing 40 mM n-octyl β-D-maltoside, 0.5 mM CHS. The SERT-8B6 Fab complex was concentrated to 2 mg/ml and 1 μM 8B6 Fab and 50 μM Br-paroxetine or I-paroxetine was added prior to crystallization.

### Synthesis of Br- and I-paroxetine

All reactions were carried out under an inert atmosphere (argon) with flame-dried glassware using standard techniques, unless otherwise specified. Anhydrous solvents were obtained by filtration through drying columns (THF, MeCN, CH_2_Cl_2_ and DMF) or used as supplied (α,α,α-trifluorotoluene). Reactions in sealed tubes were run using Biotage microwave vials (2–5 ml or 10–20 ml recommended volumes). Aluminum caps equipped with molded butyl/PTFE septa were used for reactions in α,α,α-trifluorotoluene and toluene. Simple butyl septa were used for reactions in other solvents. Chromatographic purification was performed using 230–400 mesh silica with the indicated solvent system according to standard techniques. Analytical thin-layer chromatography (TLC) was performed on precoated, glass-backed silica gel plates. Visualization of the developed chromatogram was performed by UV absorbance (254 nm) and/or stained with a ninhydrin solution in ethanol. HPLC analyses were carried out on an Agilent 1260 Infinity Series system, employing Daicel Chiracel columns, under the indicated conditions. The high-resolution mass spectrometry (HRMS) analyses were performed using electrospray ion source (ESI). ESI was performed using a Waters LCT Premier equipped with an ESI source operated either in positive or negative ion mode. The software used was MassLynx 4.1; this software does not account for the electron and all the calibrations/references are calculated accordingly, *i.e.* [M+H]^+^ is detected and the mass is calibrated to output [M+H]. Melting points are uncorrected. Infrared spectra (FTIR) were recorded in reciprocal centimeters (cm^−1^).

Nuclear magnetic resonance spectra were recorded on 400 or 500 MHz spectrometers. The frequency used to record the NMR spectra is given in each assignment and spectrum (^1^H NMR at 400 or 500 MHz; ^13^C NMR at 101 MHz or 126 MHz). Chemical shifts for ^1^H NMR spectra were recorded in parts per million from tetramethylsilane with the residual protonated solvent resonance as the internal standard (CHCl_3_: δ 7.27 ppm, (CD_2_H)_2_SO: δ 2.50 ppm, CD_2_HOD: δ 3.31 ppm). Data was reported as follows: chemical shift (multiplicity [s = singlet, d = doublet, t = triplet, m = multiplet and br = broad], coupling constant, integration and assignment). *J* values are reported in Hz. All multiplet signals were quoted over a chemical shift range. ^13^C NMR spectra were recorded with complete proton decoupling. Chemical shifts were reported in parts per million from tetramethylsilane with the solvent resonance as the internal standard (^13^CDCl_3_: δ 77.0 ppm, (^13^CD_3_)_2_SO: δ 39.5 ppm, ^13^CD_3_OD: δ 49.0 ppm). Assignments of ^1^H and ^13^C spectra, as well as *cis-*or *trans*-configuration, were based upon the analysis of δ and *J* values, analogy with previously reported compounds^27^, as well as DEPT, COSY and HSQC experiments, where appropriate. All Boc containing compounds appeared as a mixture of rotamers in the NMR spectra at room temperature. In some cases, NMR experiments for these compounds were carried out at 373 K to coalesce the signals, which is indicated in parentheses where appropriate. For NMR analysis performed at room temperature, 2D NMR experiments (COSY and HSQC) are also presented when useful for the assignments. Observed optical rotation (α’) was measured at the indicated temperature (T °C) and values were converted to the corresponding specific rotations 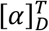 in deg cm^2^ g^−1^, concentration (*c*) in g per 100 mL. Full details of the synthetic route, using enantiopure and racemic substrates, and NMR spectra of all reaction intermediates, **2** and **3**, and HPLC analysis are cataloged in Supplementary File 1.

### Crystallization

Crystals of ts2-inactive SERT-8B6 Fab complex were grown by hanging-drop vapor diffusion at 4 °C at a ratio of 2:1 (v/v) protein:reservoir. Br-paroxetine crystals were grown using reservoir solution containing 50 mM Tris pH 8.5, 20 mM Na_2_(SO4), 20 mM LiCl_2_, 36% PEG 400, and 0.5% 6-aminohexanoic acid. I-paroxetine crystals were grown using a reservoir solution containing 100 mM HEPES pH 7.5, 40 mM MgCl_2_, and 32% PEG 400.

### X-ray data collection

Crystals were harvested and flash cooled in liquid nitrogen. Data was collected at the Advanced Photon Source (Argonne National Laboratory, beamline 24-ID-C). Data for Br-paroxetine was collected at a wavelength of 0.91840 Å and at 1.37760 Å for I-paroxetine. X-ray data sets were processed with XDS^40^. Molecular replacement was performed with coordinates from a previously determined SERT-paroxetine structure (Protein Data Bank (PDB) code: 6AWN)^8^ using PHASER^41^. The refinements were carried out in PHENIX^42^ followed by generation of the anomalous difference maps.

### Cryo-EM grid preparation

To promote the inclusion of particles in thin ice, 100 μM fluorinated octyl-maltoside (final concentration) from a 10 mM stock was added to SERT-8B6 complexes immediately prior to vitrification. Quantifoil holey carbon gold grids, 2.0/2.0 μm, size/hole space, 200 mesh) were glow discharged for 60 s at 15 mA. SERT-8B6 Fab complex (2.5 μl) was applied to the grid followed by blotting for 2 s in the vitrobot and plunging into liquid ethane cooled by liquid N_2_.

### Cryo-EM data collection and processing

Images were acquired using the automated program SerialEM^43^ on a FEI Titan Krios transmission electron microscope, operating at 300 keV and equipped with a Gatan Image Filter with the slit width set to 20 eV. A Gatan K3 direct electron detector was used to record movies in super-resolution counting mode with a binned pixel size of 0.648 Å per pixel. The defocus values ranged from −0.8 to −2.2 μm. Exposures of 1.0-1.5 s were dose fractioned into 40 frames, resulting in a total dose of 54-60 *e*^−^ Å^−2^. Movies were corrected for beam-induced motion using MotionCor2^44^ with 5×5 patching. The contrast transfer function (CTF) parameters for each micrograph was determined using ctffind4^45^ and particles were picked either using DoG-Picker^46^ or blob-based picking in cryoSPARC^47^. DoG or cryoSPARC picked particles were independently subjected to 3D classification against a low-resolution volume of the SERT-8B6 complex. After sorting, the DoG and cryoSPARC picked particles were combined in Relion^48^ and the duplicate picks were removed (particle picks that are less than 100 Å of one another were considered duplicates). Combined particles were further sorted using reference-free 2D classification in cryoSPARC, followed by refinement in Relion and further 3D classification. Particles were then re-extracted (box size 400, 0.648 Å per pixel) and subjected to non-uniform refinement in cryoSPARC. Local refinement was then performed in *cis*TEM^49^ with a mask that excludes the micelle and Fab constant domain to remove low-resolution features. The high-resolution refinement limit was incrementally increased while maintaining a correlation of 0.95 or better until no improvement in map quality was observed. The resolution of the reconstructions was accessed using the Fourier shell correlation (FSC) criterion and a threshold of 0.143^50^. Map sharpening was performed using local sharpening in PHENIX.

### Cryo-EM model building and refinement

A starting model was generated by fitting the x-ray structure of SERT-8B6 Fab paroxetine complex (PDB code: 6AWN) into the cryo-EM reconstruction in Chimera^51^. Several rounds of manual adjustment and rebuilding were performed in Coot^52^, followed by real space refinement in PHENIX. For cross-validation, the FSC curve between the refined model and half maps was calculated and compared to prevent overfitting. Molprobity was used to evaluate the stereochemistry and geometry of the structures^53^.

### Radioligand binding and uptake assays

Competition binding experiments were performed using scintillation proximity assays (SPA)^36,38^. The assays contained ~10 nM SERT, 0.5 mg/ml Cu-Ysi beads in TBS with 1 mM DDM, 0.2 mM CHS, and 10 nM [^3^H]citalopram and 0.01 nM–1 mM of the cold competitors. Experiments were measured in triplicate. The error bars for each data point represent the s.e.m. Ki values were determined with the Cheng–Prusoff equation^54^ in GraphPad Prism. Uptake was measured as described previously in 96-well plates with [^3^H]5-HT diluted 1:100 with unlabeled 5-HT. After 24 hrs, cells were washed into uptake buffer (25 mM HEPES-Tris, pH 7.0, 130 mM NaCl, 5.4 mM KCl, 1.2 mM CaCl_2_, 1.2 mM MgSO^4^, 1 mM ascorbic acid and 5 mM glucose) containing 0.001 – 10,000 nM of the inhibitor. [^3^H]5-HT was added to the cells and uptake was stopped by washing cells rapidly three times with uptake buffer. Cells were solubilized with 1% Triton-X100, followed by the addition of 200 μl of scintillation fluid to each well. The amount of labelled 5-HT was measured using a MicroBeta scintillation counter. Data were fit to a sigmoidal dose-response curve.

## ACKOWLEDGEMENTS

We thank L. Vaskalis for assistance with figures and H. Owen for help with manuscript preparation. We acknowledge the staff of the Northeastern Collaborative Access Team at the Advanced Photon Source. A portion of this research was supported by NIH grant U24GM129547 and performed at the PNCC at OHSU and accessed through EMSL (grid.436923.9), a DOE Office of Science User Facility sponsored by the Office of Biological and Environmental Research. We are particularly grateful to Bernard and Jennifer LaCroute for their generous support. This work was funded by the NIH (5R37MH070039). E.G. is an investigator of the Howard Hughes Medical Institute.

We gratefully acknowledge The Royal Society [University Research Fellowship, UF140161 (to J.A.B.), URF Appointed Grant RG150444].

---

**Supplemental Figure 2.**
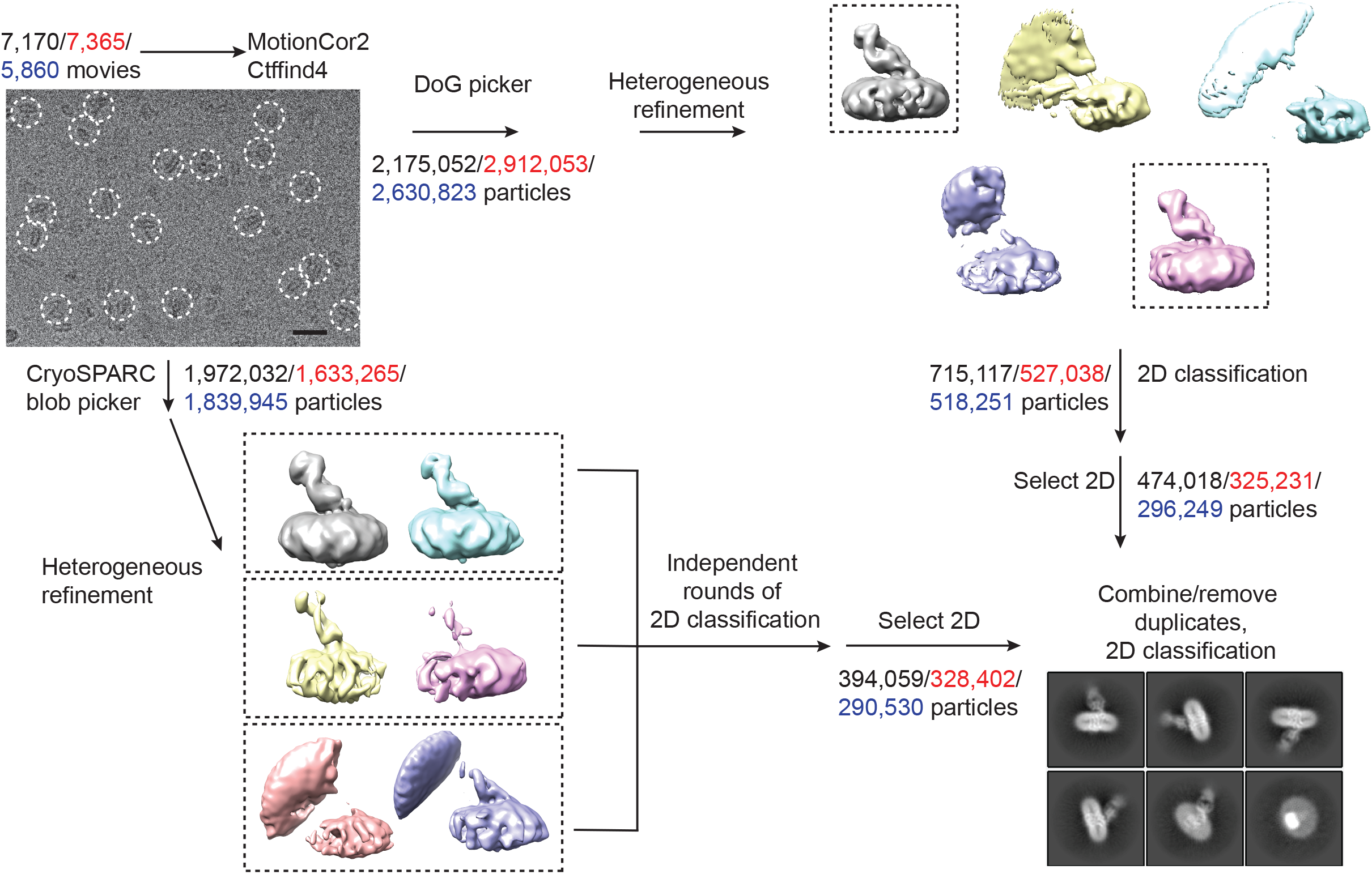
Work-flow of cryo-EM data processing of ΔN72, ΔC13 SERT/8B6 Fab/paroxetine complexes. A representative zoomed, motion-corrected micrograph with individual single particles circled in white. Bar equals 20 nm. Motion-correction and CTF estimation was performed using MotionCor2 and Ctffind4. The number of movies/particles collected for each data set are shown in black (paroxetine), red (Br-paroxetine), and blue (I-paroxetine). After particle picking using either DoG picker or the blob picker in cryoSPARC, particles were sorted using heterogeneous refinement in cryoSPARC followed by 2D classification. For the DoG-picked particles, 3D classes containing SERT-Fab features (boxed) were combined and subjected to 2D classification. For cryoSPARC-picked particles, heterogeneous refinement was also used to initially sort particles in cryoSPARC. Classes with similar features (boxed) were combined, subjected to three independent 2D classifications, and 2D classes containing SERT-Fab features were combined. Particles picked by both methods were combined and duplicate particle-picks were removed in Relion (particle picks that are less than 100 Å of one another were considered duplicates).

**Supplemental Figure 3.**
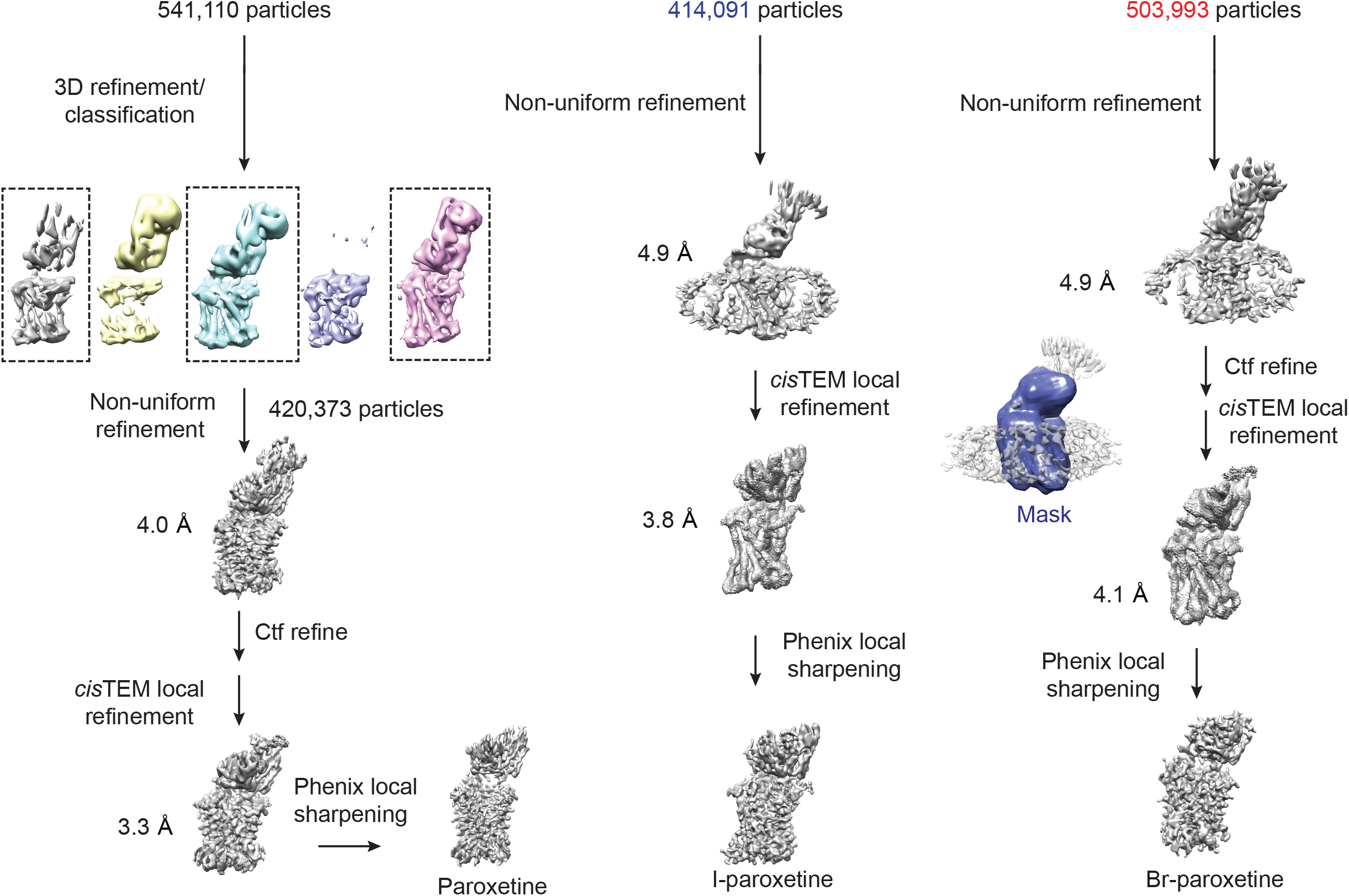
3D refinement of ΔN72, ΔC13 SERT/8B6 Fab/paroxetine complexes. For the paroxetine complex, 3D refinement was performed in Relion followed by 3D classification without alignment and a mask which isolated SERT and Fab. 3D classification was not performed on the Br-paroxetine and I-paroxetine particles. Particles were further refined using non-uniform refinement in cryoSPARC, followed by local refinement in cisTEM with a mask which isolated SERT and the Fab variable domain and removed the Fab constant domain and micelle (mask is shown overlaid in blue on top of the Br-paroxetine reconstruction). The final reconstructed volume was sharpened using Phenix local sharpening.

**Supplemental Figure 4.**
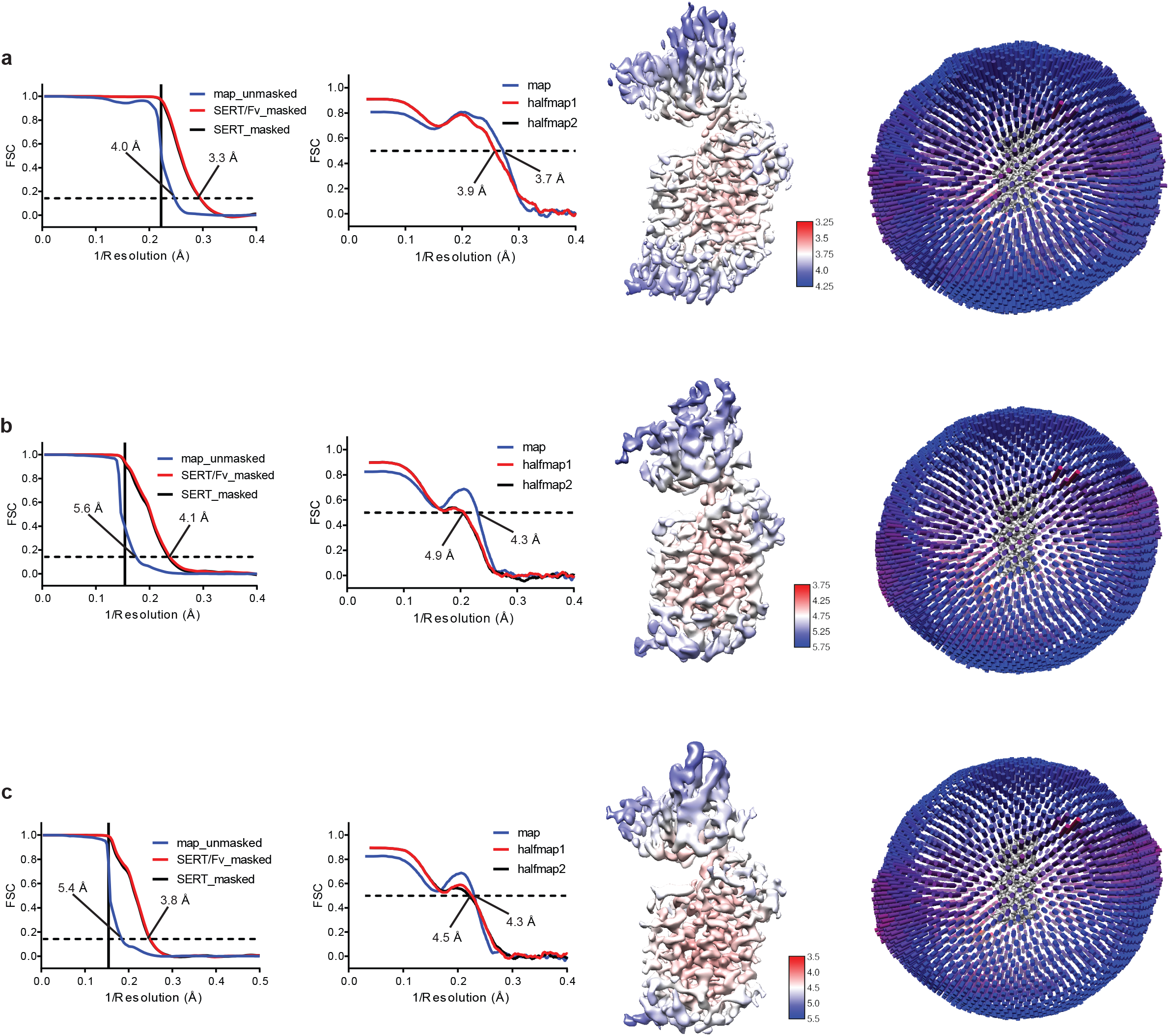
Cryo-EM reconstruction of ΔN72, ΔC13 SERT/8B6 Fab/paroxetine complexes. **a**, Reconstruction of SERT-8B6 paroxetine complex. Left panel, FSC curves for cross-validation, the final map (blue), masked SERT-Fv (red), and a mask which isolated SERT (black). The high-resolution limit cutoff for refinement was 4.5 Å. Middle left panel: model vs. half map 1 (working, red), half map 2 (free, black), model vs. final map (blue). Middle right panel: cryo-EM density map colored by local resolution estimation. Right panel: the angular distribution of particles used in the final reconstruction. **b**, Reconstruction of the SERT-8B6 Br-paroxetine complex. The high-resolution limit cutoff for refinement was 6.5 Å. **c**, Reconstruction of the SERT-8B6 I-paroxetine complex. The high-resolution limit cutoff for refinement was 6.5 Å.

**Supplemental Figure 5.**
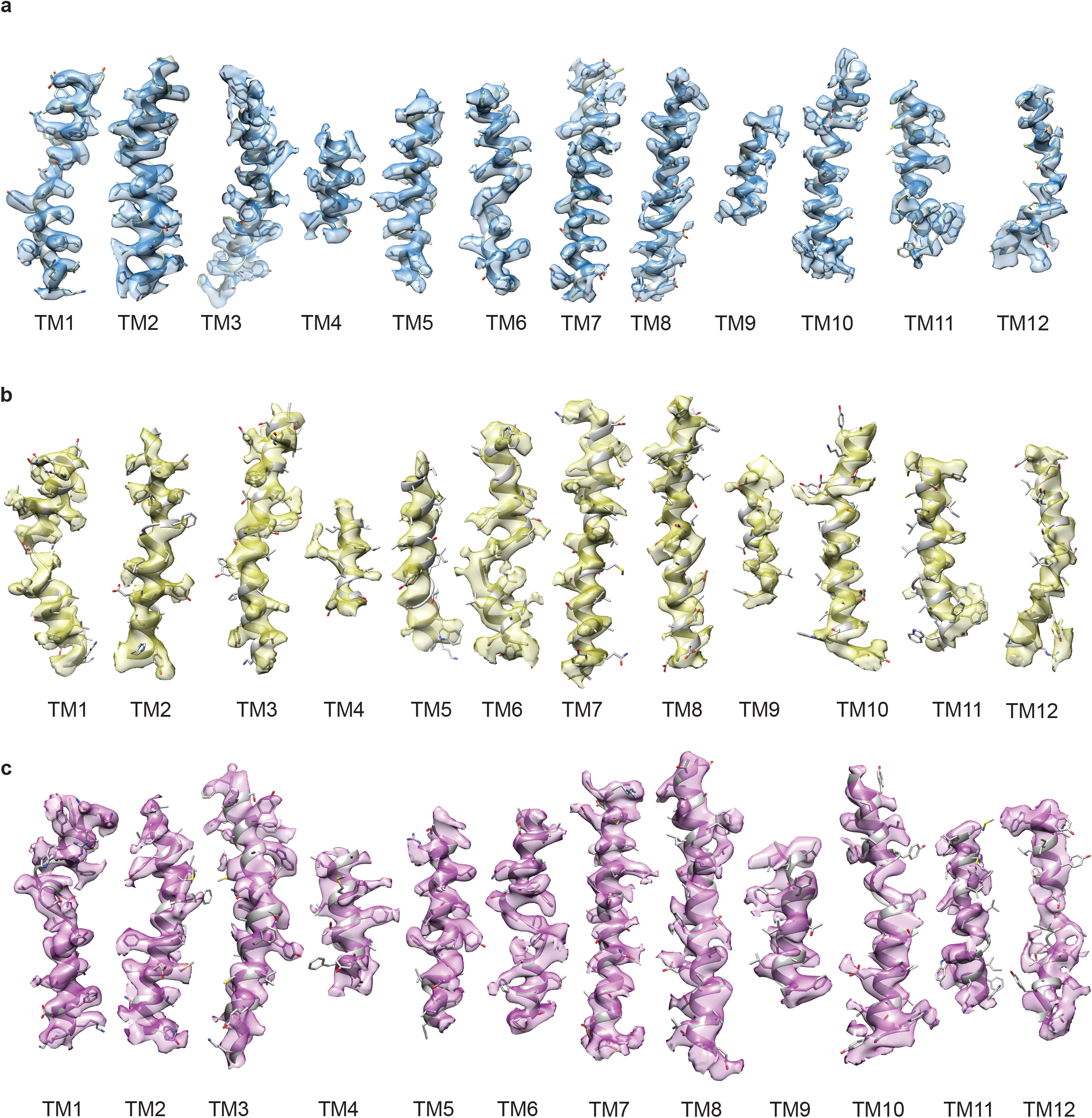
Cryo-EM density segments of the transmembrane helices. **a**, Density of TM1-12 of the paroxetine reconstruction, shown in blue. **b**, Density of TM1-12 of the Br-paroxetine reconstruction, shown in yellow. **c**, Density of TM1-12 of the I-paroxetine reconstruction, shown in purple.

**Supplemental Figure 6.**
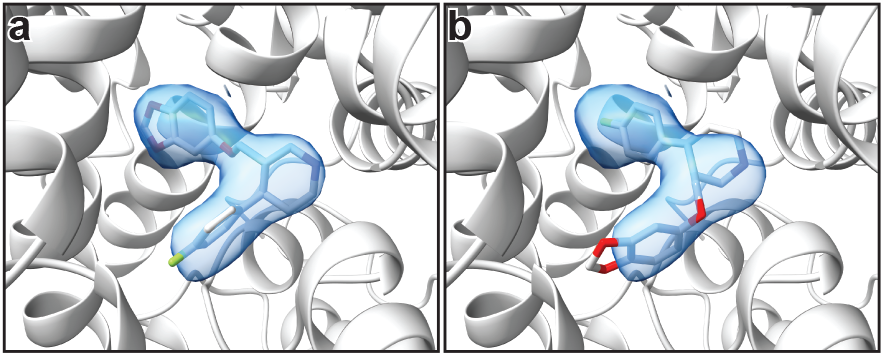
Comparison of the fit of paroxetine in the ABC and ACB poses. **a**, Shows the fit of paroxetine to the cryo-EM density in the ABC pose. **b**, Shows the fit in the ACB pose.

