## Supplementary File 1 for "Chemical and structural investigation of the paroxetine-human serotonin transporter complex"

**Supplementary file 1. Synthesis of paroxetine analogues.**

**Reagents**

Commercial reagents were used as supplied or purified by standard techniques where necessary.

$Pd(OAc)_2$, 8-Aminoquinoline, 1-(*tert*-butoxycarbonyl)piperidine-3-carboxylic acid and (*R*)-1-(*tert*-butoxycarbonyl)piperidine-3-carboxylic acid were purchased from Fluorochem Ltd and used as supplied.

PivOH and α,α,α-trifluorotoluene were purchased from Sigma-Aldrich Company Ltd and used as supplied.

$K_2CO_3$ was purchased from Sigma-Aldrich Company Ltd and flame-dried before use as part of reaction set-up.

Purity: $Pd(OAc)_2$, >98%; PivOH, 99%; $K_2CO_3$, ≥98% (powder, –325 mesh), α,α,α-trifluorotoluene, anhydrous, ≥99%.

Racemic and enantioenriched substrates *tert*-butyl (±)-3-(quinoline-8-ylcarbamoyl)piperidine-1-carboxylate (**(±)-S1**) and *tert*-butyl (–)-(*R*)-3-(quinolin-8-ylcarbamoyl)piperidine-1-carboxylate (**(–)-5**) were prepared by amide coupling of commercially available 8-aminoquinoline and the corresponding carboxylic acid (1-(*tert*-butoxycarbonyl)piperidine-3-carboxylic acid and (*R*)-1-(*tert*-butoxycarbonyl)piperidine-3-carboxylic acid, respectively) according to our previously reported procedures.[1]

**Structures of Additional Compounds in SI**

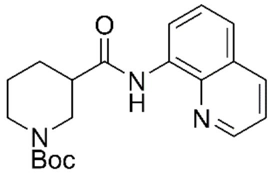

**(±)-S1**

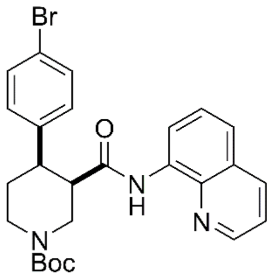

**(±)-S2a**

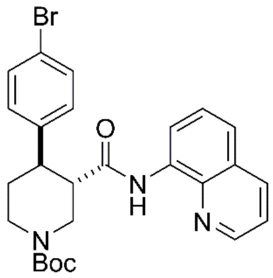

**(±)-S3a**

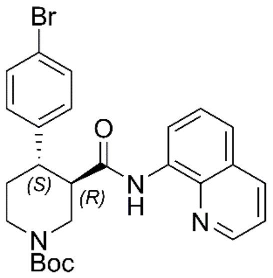

**(−)-S3a**

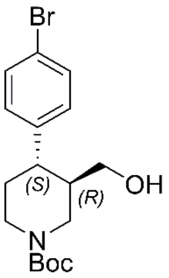

**(+)-S4a**

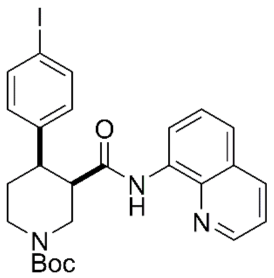

**(±)-S2b**

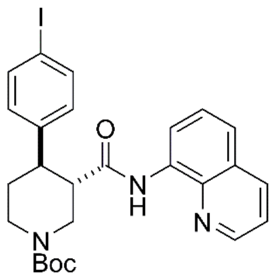

**(±)-S3b**

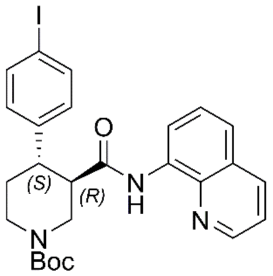

**(−)-S3b**

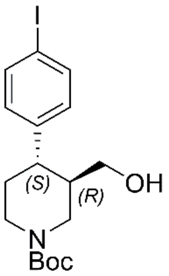

**(+)-S4b**

**Full Synthetic Route to Racemic and Enantioenriched Br-Piperidine Derivatives (±)-S2a, (±)-S3a, (+)-6a, (+)-7a, (–)-S3a, (–)-8a, (+)-S4a, (–)-9a and Br-(–)-paroxetine 2**

In order to evaluate the enantiomeric excess of key intermediates **(+)-6a** and **(+)-7a** by chiral HPLC, the C–H arylation with 4-bromo iodobenzene was performed on both racemic (**(±)-S1**) and enantioenriched **(–)-5** piperidine amide substrates (Scheme S1).

The racemic synthesis was performed on a 0.5 mmol scale according to our previously reported protocol,[1] and afforded *cis*-arylated derivatives **(±)-S2a** in 34% (Scheme S1a). A minor *trans*-functionalized product **(±)-S3a**, formed via a *trans*-palladacycle,[1] was also isolated in 14%.
C–H Arylation of enantioenriched substrate **(–)-5** proceed smoothly on a 4.0 mmol scale, and *cis*- and *trans*-piperidine products **(+)-6a** and **(–)-S3a** were isolated as single enantiomers in very similar yields (Scheme S1b). Subsequent treatment of enantiopure *cis*-derivative **(+)-6a** with DBU at 100 °C afforded the *trans*-diastereomer as the right-handed enantiomer **(+)-7a** in 94% yield.

**Scheme S1. Synthetic sequence, including the Pd-catalyzed C(4)–H arylation step, to access racemic and enantioenriched *cis*- and *trans*-piperidine amide derivatives (±)-S2a, (±)-S3a, (+)-6a, (+)-7a and (–)-S3a**

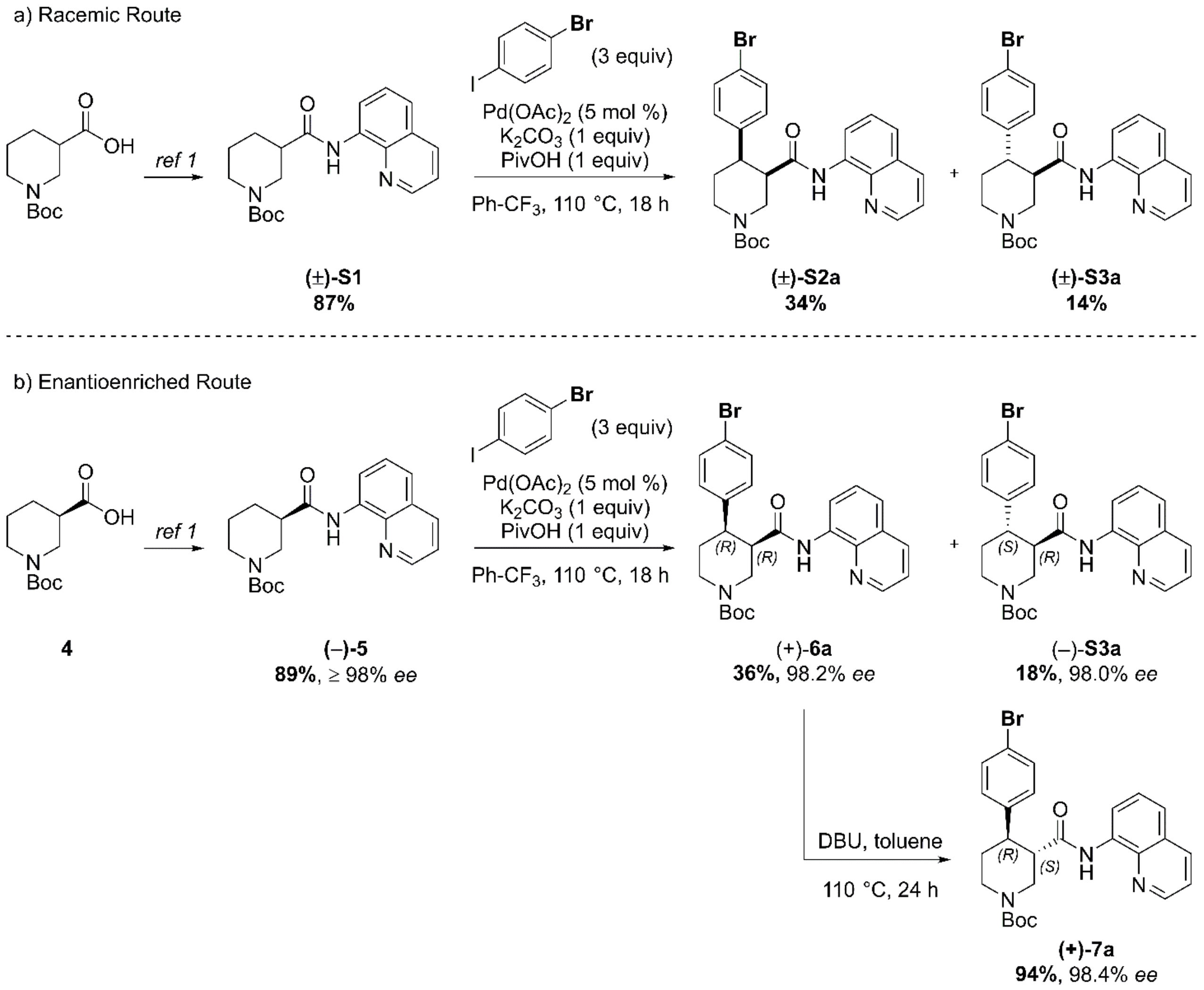


a) C–H Arylation conditions: **(±)-S1** (0.5 mmol, 1 equiv) Ph-$CF_3$ (500 μL, 1 M). b) C–H Arylation conditions: **(–)-5** (4.0 mmol, 1 equiv), Ph-$CF_3$ (2.0 mL, 2 M).

The enantiomeric excess of alcohol intermediates **(+)-S4a** and **(–)-8a** was evaluated after aminoquinoline removal on both enantiomeric *trans*-derivatives **(–)-S3a** and **(+)-7a** (Scheme S2).

No undesired debromination was observed for the reductive aminoquinoline removal, and enantiopure alcohols **(+)-S4a** and **(–)-8a** were obtained in 70% and 77% yield, respectively.
No erosion of enantiopurity should be expected after this step, given the literature precedents on the synthesis of (–)-paroxetine[2] and the absence of acidic protons in the substrate. Therefore, the synthesis was continued exclusively on alcohol derivative **(–)-8a**. O-Alkylation and Boc-deprotection with HCl finally afforded enantiopure Br-(–)-paroxetine analogue **2** as the corresponding hydrochloride salt in 12% yield over 8 steps from commercial material.

**Scheme S2. Reductive aminoquinoline removal and final steps in the synthesis of Br-(–)-paroxetine 2.**

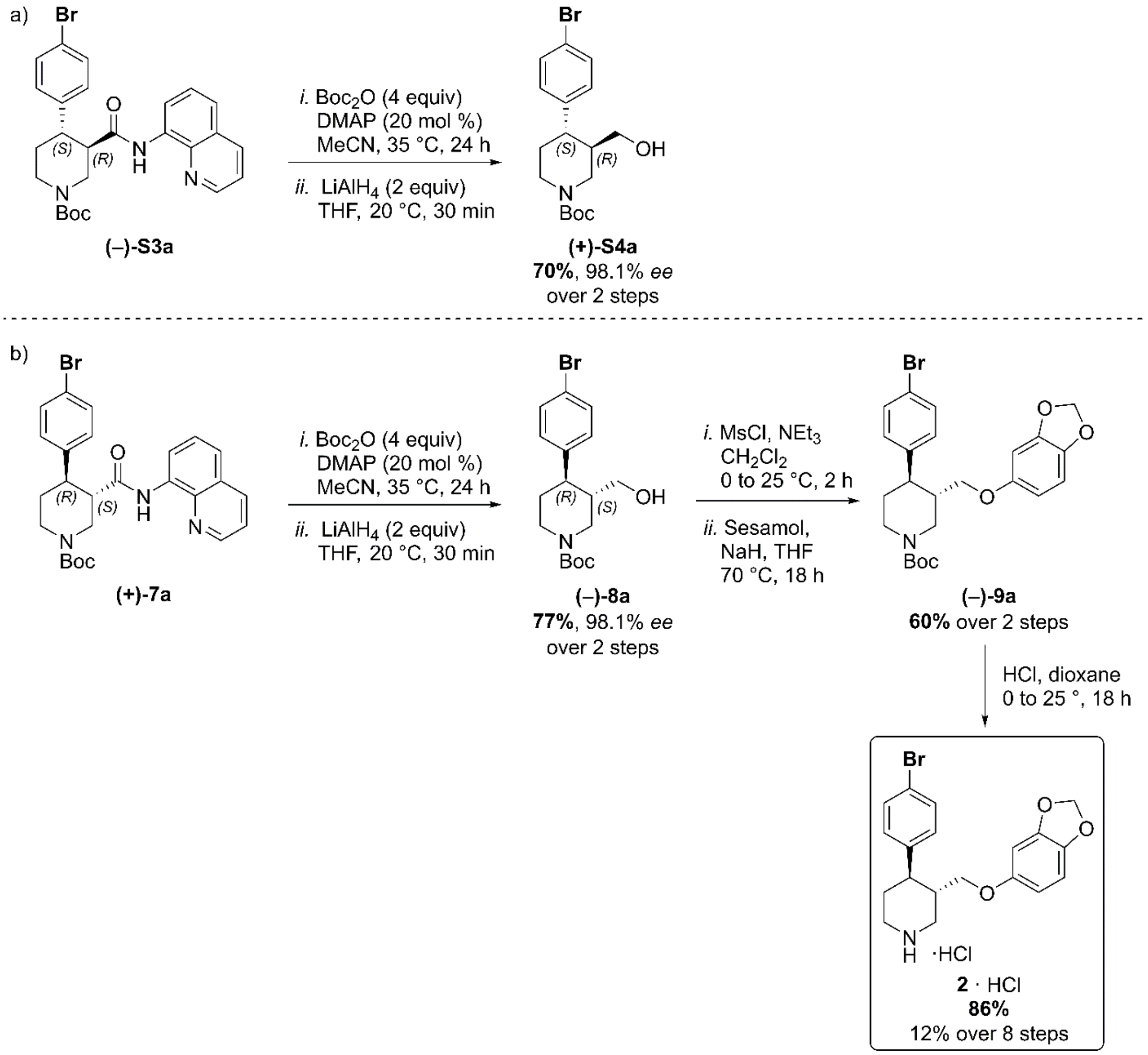


a) AQ removal on enantiomerically pure *trans*-piperidine **(–)-S3a** (0.2 mmol, 1 equiv). b) AQ removal on enantiomerically pure *trans*-piperidine **(+)-7a** (1.1 mmol, 1 equiv) and final steps in the synthesis of Br-(–)-paroxetine **2**.

**Full Synthetic Route to Racemic and Enantioenriched I-Piperidine Derivatives (±)-S2b, (±)-S3b, (+)-6b, (+)-7b, (–)-S3b, (–)-8b, (+)-S4b, (–)-9b and I-(–)-paroxetine 3**

Similarly to the Br-analogue, C–H arylation with 1,4-diiodobenzene was performed on both racemic (**(±)-S1**) and enantioenriched (**(–)-5**) piperidine amide substrates (Scheme S3).

The reaction proceeded well on both substrates affording racemic *cis*- and *trans*-arylated products **(±)-S2b** and **(±)-S3b** in 35% and 19% yield, and enantioenriched *cis*- and *trans*-derivatives **(+)-6b** and **(–)-S3b** in 35% and 20% yield respectively.

**Scheme S3. Synthetic sequence, including the Pd-catalyzed C(4)–H arylation step, to access racemic and enantioenriched *cis*- and *trans*-piperidine amide derivatives (±)-S2b, (±)-S3b, (+)-6b, (+)-7b and (–)-S3b**

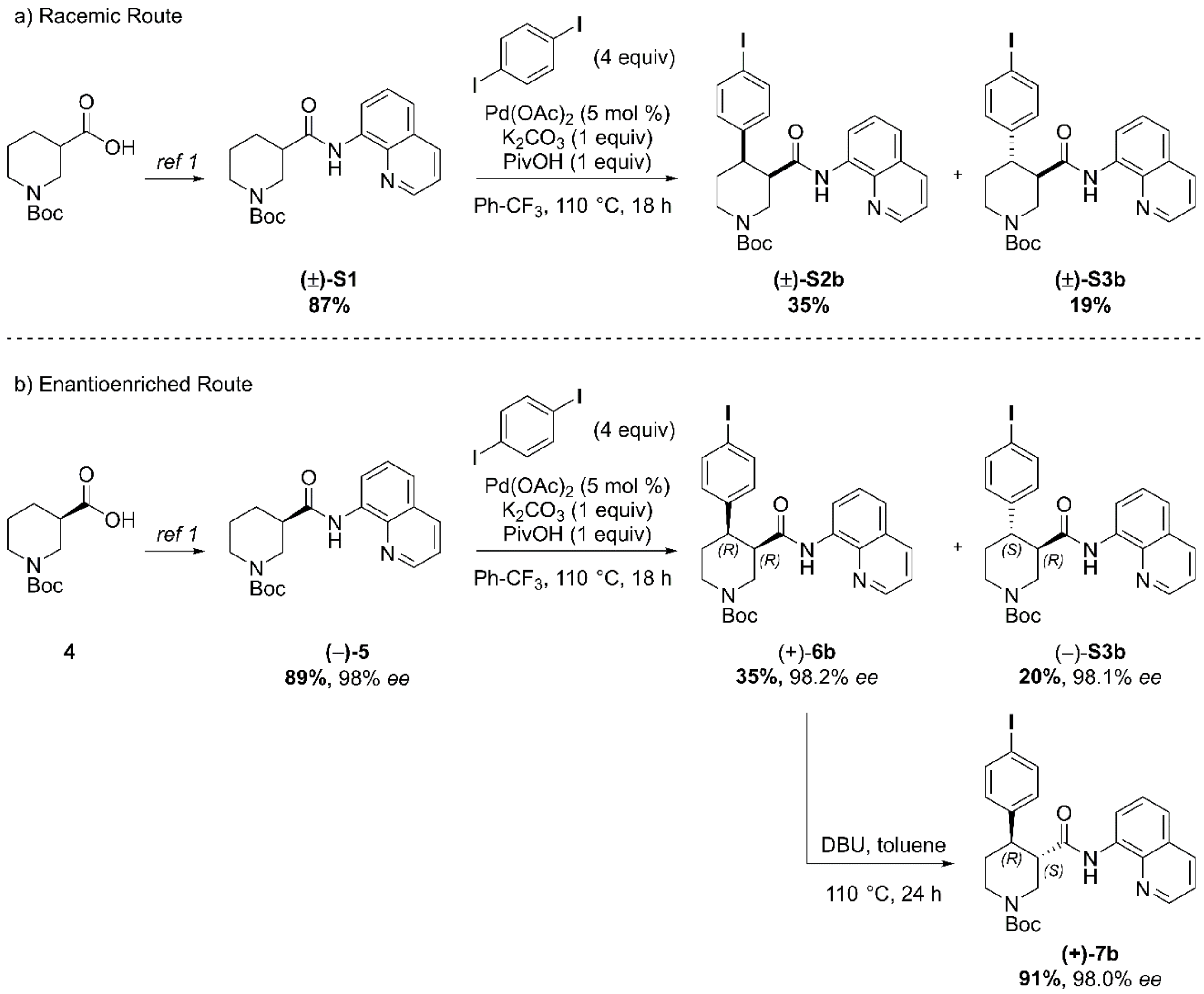


a) C–H Arylation conditions: **(±)-S1** (0.5 mmol, 1 equiv) Ph-$CF_3$ (500 μL, 1 M). b) C–H Arylation conditions: **(–)-5** (4.0 mmol, 1 equiv), Ph-$CF_3$ (2.0 mL, 2 M).

Reductive aminoquinoline cleavage was again performed to access enantiomeric *trans*-piperidine alcohols **(+)-S4b** and **(–)-8b** (Scheme S4).

In both cases a small degree of $LiAlH_4$-mediated dehalogenation was observed, and an inseparable mixture of the desired product and 10–15% of deiodinated material was isolated.[3] However, the contaminant could be effectively removed after O-Alkylation, affording the pure aryl ether derivative **(–)-9b** in 71% yield. Final HCl-mediated Boc deprotection formed the desired I-(–)-paroxetine **3** as the corresponding HCl salt in 81% yield (12% yield over 8 steps from commercial material).

**Scheme S4. Reductive aminoquinoline removal and final steps in the synthesis of I-(–)-paroxetine 3.**

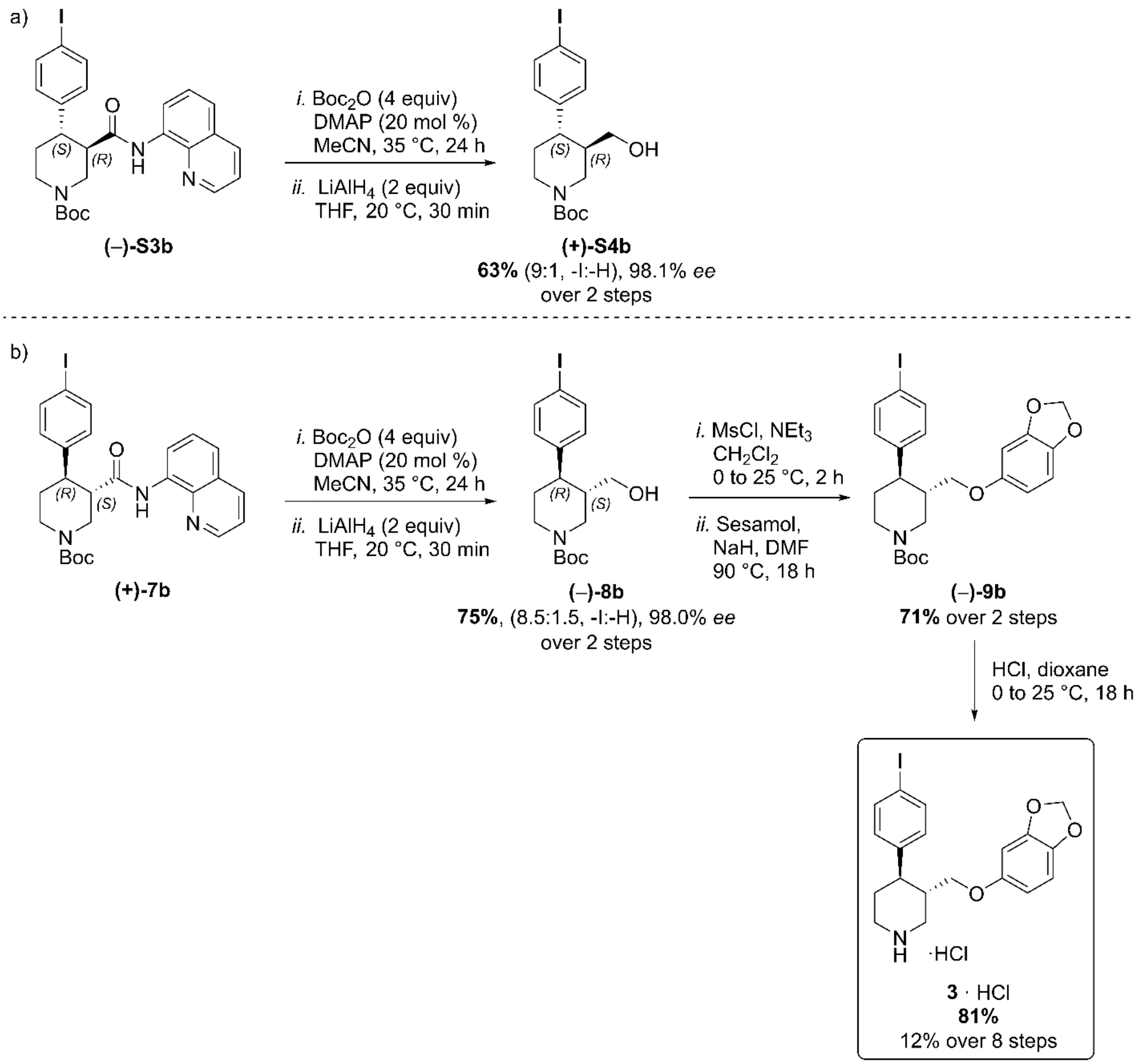


a) AQ removal on enantiomerically pure *trans*-piperidine **(–)-S3b** (0.2 mmol, 1 equiv). b) AQ removal on enantiomerically pure *trans*-piperidine **(+)-7b** (1.0 mmol, 1 equiv) and final steps in the synthesis of I-(–)-paroxetine **3**.

#### HPLC Traces for Racemic, Scalemic and Enantioenriched Br-Piperidine Derivatives (±)-S2a, (±)-S3a, (+)-6a, (+)-7a, (–)-S3a, (–)-8a and (+)-S4a

##### ***tert*-Butyl *cis*-(±)-4-(4-bromophenyl)-3-(quinolin-8-ylcarbamoyl)piperidine-1-carboxylate ((±)-S2a)**

**Conditions**: Chiralpak IA 3-column, 85:15 *n*-hexane:*i*-PrOH, flow rate: 1 mL·min$^{-1}$, 35 ºC, UV detection wavelength: 210.4 nm. Retention times: 11.9 min (3*S*,4*S* enantiomer), 17.3 min (3*R*,4*R* enantiomer).

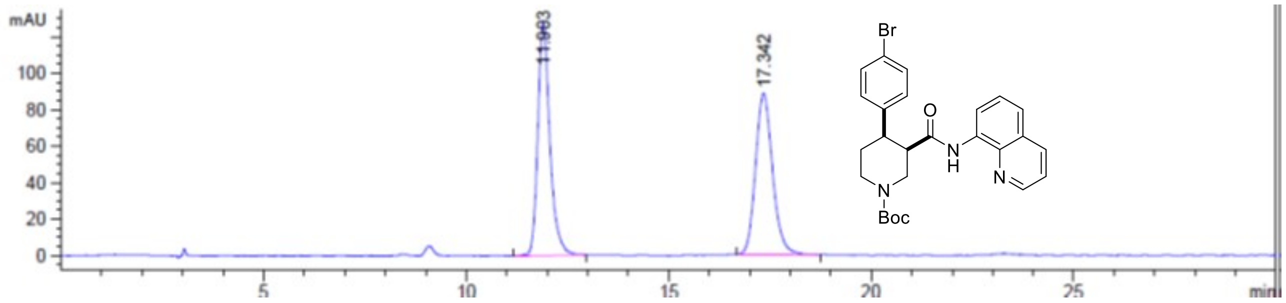


Signal 3: DAD1 C, Sig=210,10 Ref=off

| Peak # | RetTime [min] | Type | Width [min] | Area [mAU*s] | Height [mAU] | Area % |
|---|---|---|---|---|---|---|
| 1 | 11.903 | BB | 0.3134 | 2649.83203 | 128.07286 | 50.6199 |
| 2 | 17.342 | BB | 0.4515 | 2584.93091 | 88.31209 | 49.3801 |

Totals : 5234.76294 216.38495

##### ***tert*-Butyl (+)-(3*R*,4*R*)-4-(4-bromophenyl)-3-(quinolin-8-ylcarbamoyl)piperidine-1-carboxylate ((+)-6a)**

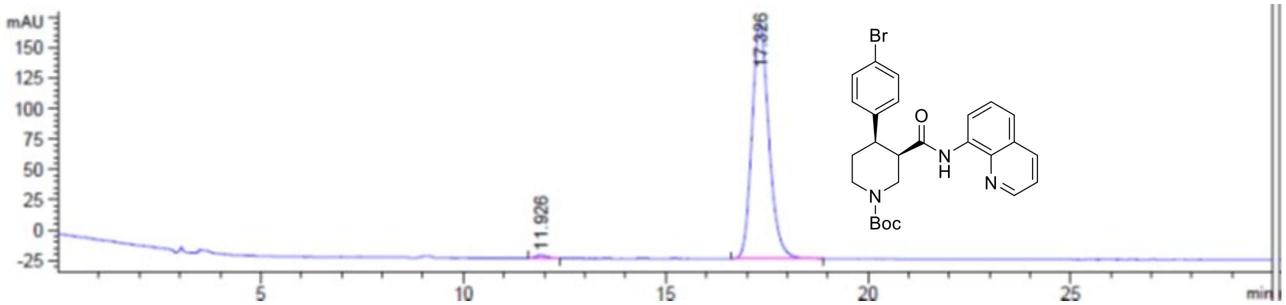


Signal 3: DAD1 C, Sig=210,10 Ref=off

| Peak # | RetTime [min] | Type | Width [min] | Area [mAU*s] | Height [mAU] | Area % |
|---|---|---|---|---|---|---|
| 1 | 11.926 | BB | 0.2556 | 51.12915 | 2.64391 | 0.8973 |
| 2 | 17.326 | BB | 0.4494 | 5647.02783 | 193.00262 | 99.1027 |

Totals : 5698.15698 195.64654

***ee* = 98.2%**

***tert*-Butyl *trans*-(±)-4-(4-bromophenyl)-3-(quinolin-8-ylcarbamoyl)piperidine-1-carboxylate ((±)-S3a)**

**Conditions**: Chiralpak IA 3-column, 85:15 *n*-hexane:*i*-PrOH, flow rate: 1 mL·min$^{-1}$, 35 ºC, UV detection wavelength: 254.1 nm. Retention times: 9.1 min (3*R*,4*S* enantiomer), 12.2 min (3*S*,4*R* enantiomer).

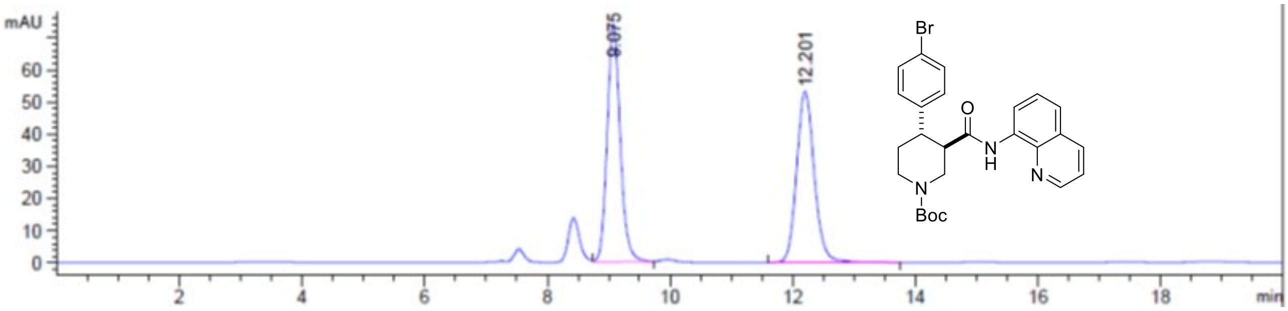


Signal 2: DAD1 B, Sig=254,10 Ref=off

| Peak # | RetTime [min] | Type | Width [min] | Area [mAU*s] | Height [mAU] | Area % |
|---|---|---|---|---|---|---|
| 1 | 9.075 | VB | 0.2266 | 1093.46033 | 73.93165 | 50.5693 |
| 2 | 12.201 | BB | 0.3080 | 1068.84167 | 53.29229 | 49.4307 |
| Totals : | | | | 2162.30200 | 127.22393 | |

***tert*-Butyl (–)-(*3R,4S*)-4-(4-bromophenyl)-3-(quinolin-8-ylcarbamoyl)piperidine-1-carboxylate ((–)-S3a)**

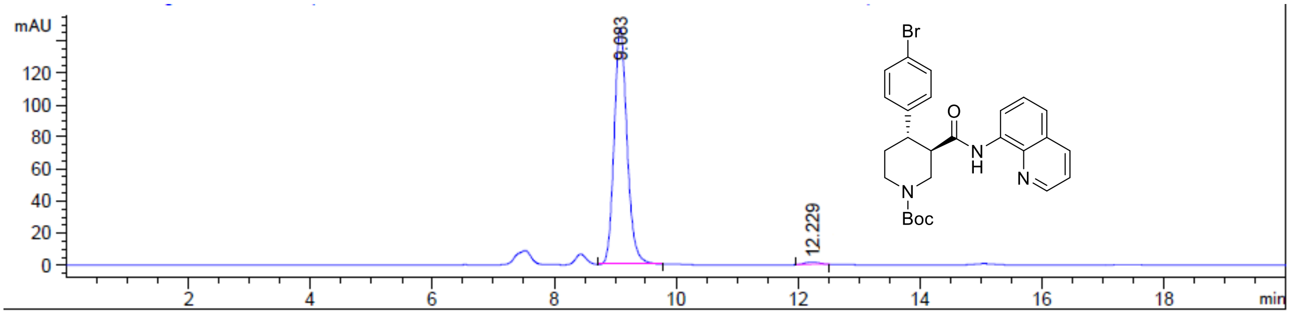


Signal 2: DAD1 B, Sig=254,10 Ref=off

| Peak # | RetTime [min] | Type | Width [min] | Area [mAU*s] | Height [mAU] | Area % |
|---|---|---|---|---|---|---|
| 1 | 9.083 | MM | 0.2445 | 2163.23486 | 147.46376 | 99.0118 |
| 2 | 12.229 | MM | 0.2733 | 21.59002 | 1.31660 | 0.9882 |
| Totals : | | | | 2184.82488 | 148.78036 | |

***ee* 98.0%**

***tert*-Butyl (+)-(*3S,4R*)-4-(4-bromophenyl)-3-(quinolin-8-ylcarbamoyl)piperidine-1-carboxylate ((+)-7a)**

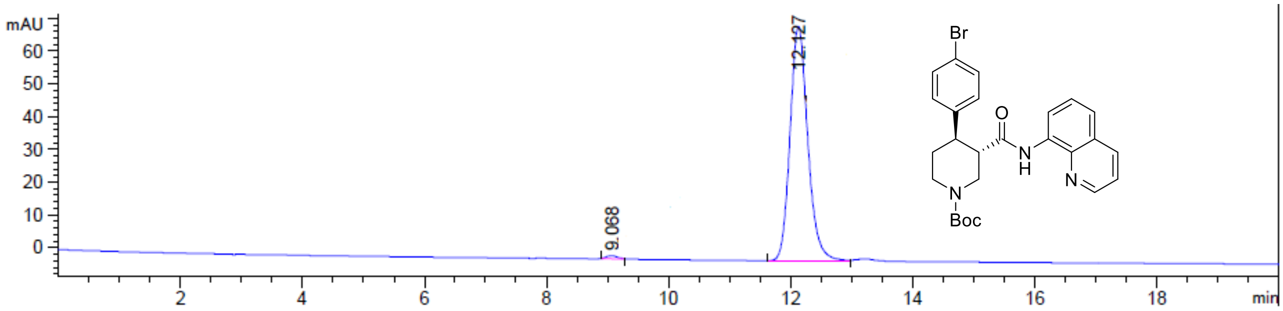


Signal 2: DAD1 B, Sig=254,10 Ref=off

| Peak # | RetTime [min] | Type | Width [min] | Area [mAU*s] | Height [mAU] | Area % |
|---|---|---|---|---|---|---|
| 1 | 9.068 | MM | 0.2108 | 11.79956 | 9.32985e-1 | 0.8026 |
| 2 | 12.127 | MM | 0.3388 | 1458.44690 | 71.75478 | 99.1974 |
| Totals : | | | | 1470.24646 | 72.68776 | |

***ee* = 98.4%**

**Scalemic Mixture of Enantioenriched Br-Piperidines (+)-S4a and (–)-8a**

**Conditions**: Chiralpak ID 3-column, 90:10 *n*-hexane:*i*-PrOH, flow rate: 1 mL·min$^{-1}$, 35 ºC, UV detection wavelength: 210.4 nm. Retention times: 8.0 min (3*R*,4*S* enantiomer), 8.6 min (3*S*,4*R* enantiomer).

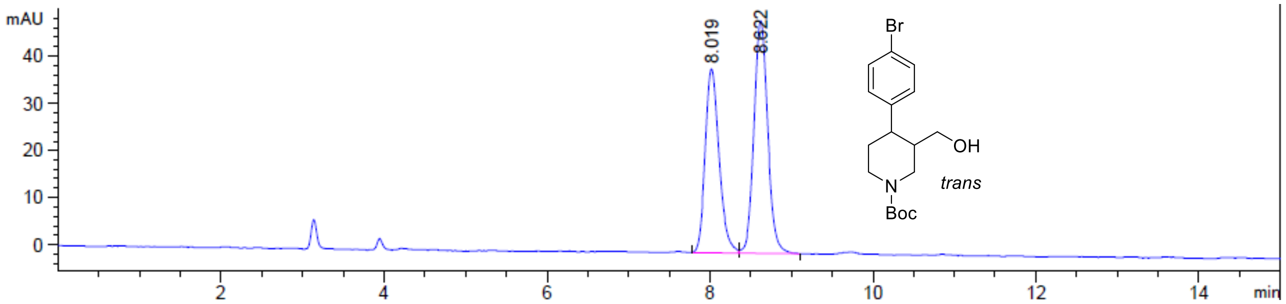


Signal 3: DAD1 C, Sig=210,10 Ref=off

| Peak # | RetTime [min] | Type | Width [min] | Area [mAU*s] | Height [mAU] | Area % |
|---|---|---|---|---|---|---|
| 1 | 8.019 | BV | 0.1793 | 458.78363 | 38.98658 | 44.3601 |
| 2 | 8.622 | VB | 0.1802 | 575.44318 | 49.30547 | 55.6399 |

Totals : 1034.22681 88.29205

***tert*-Butyl (+)-(*3R,4S*)-4-(4-bromophenyl)-3-(hydroxymethyl)piperidine-1-carboxylate ((+)-S4a)**

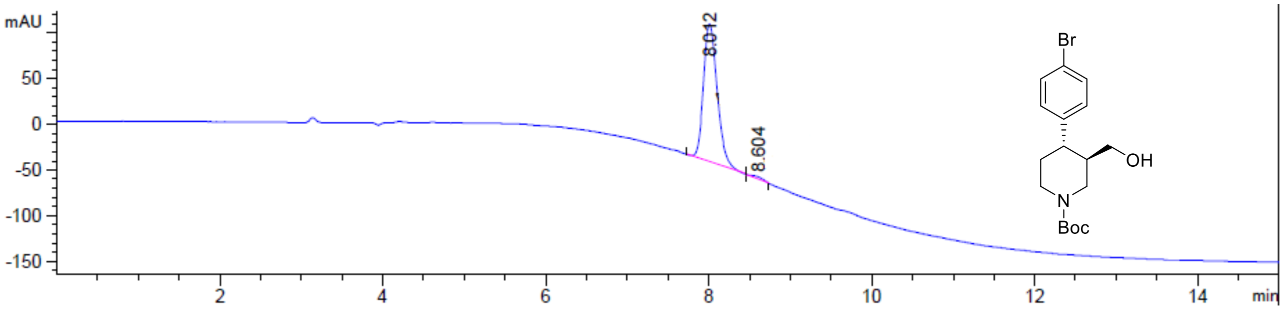


Signal 3: DAD1 C, Sig=210,10 Ref=off

| Peak # | RetTime [min] | Type | Width [min] | Area [mAU*s] | Height [mAU] | Area % |
|---|---|---|---|---|---|---|
| 1 | 8.012 | MM | 0.1924 | 1740.07422 | 150.75604 | 99.0603 |
| 2 | 8.604 | MM | 0.1355 | 16.50659 | 2.03064 | 0.9397 |

Totals : 1756.58081 152.78669

***ee* = 98.1%**

**_tert_-Butyl (–)-(_3S,4R_)-4-(4-bromophenyl)-3-(hydroxymethyl)piperidine-1-carboxylate ((–)-8a)**

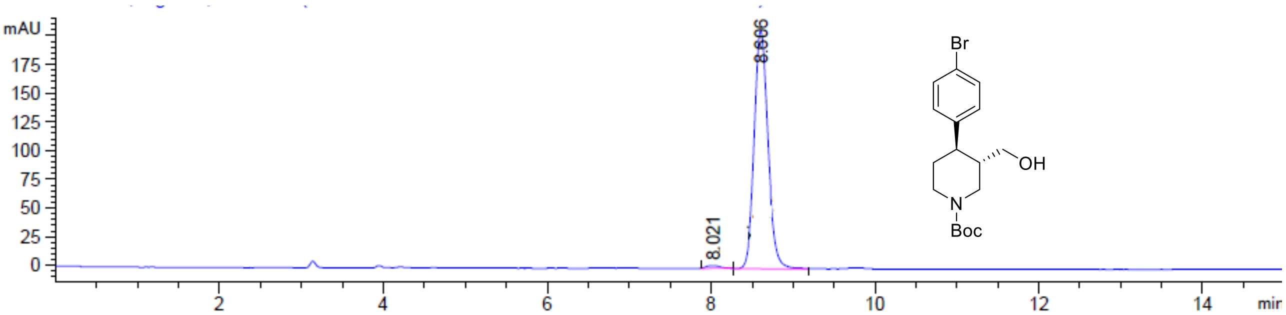


Signal 3: DAD1 C, Sig=210,10 Ref=off

| Peak # | RetTime [min] | Type | Width [min] | Area [mAU*s] | Height [mAU] | Area % |
|---|---|---|---|---|---|---|
| 1 | 8.021 | MM | 0.1640 | 23.16720 | 2.35474 | 0.9470 |
| 2 | 8.606 | MM | 0.1943 | 2423.17212 | 207.84224 | 99.0530 |
| Totals : | | | | 2446.33932 | 210.19698 | |

***ee* = 98.1%**

**HPLC Traces for Racemic, Scalemic and Enantioenriched I-Piperidine Derivatives (±)-S2b, (±)-S3b, (+)-6b, (+)-7b, (–)-S3b, (–)-8b and (+)-S4b**

***tert*-Butyl *cis*-(±)-4-(4-iodophenyl)-3-(quinolin-8-ylcarbamoyl)piperidine-1-carboxylate ((±)-S2b):**

**Conditions**: Chiralpak IA 3-column, 85:15 *n*-hexane:*i*-PrOH, flow rate: 1 mL·min$^{-1}$, 35 °C, UV detection wavelength: 210.4 nm. Retention times: 12.2 min (3*S*,4*S* enantiomer), 17.7 min (3*R*,4*R* enantiomer).

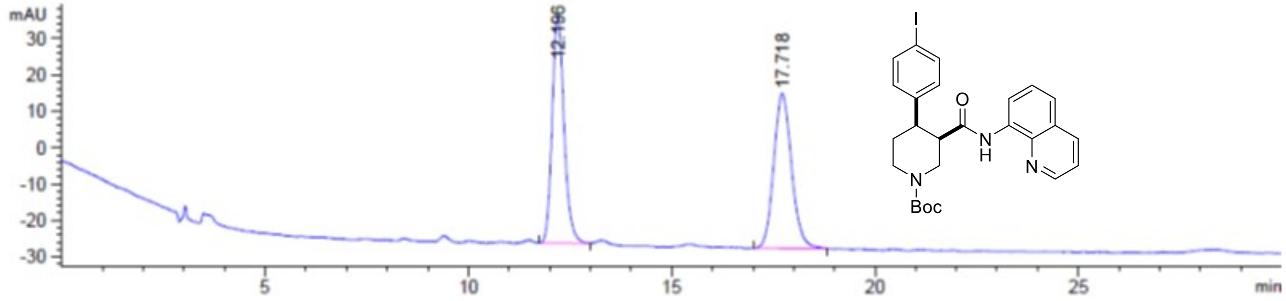


Signal 3: DAD1 C, Sig=210,10 Ref=off

| Peak # | RetTime [min] | Type | Width [min] | Area [mAU*s] | Height [mAU] | Area % |
|---|---|---|---|---|---|---|
| 1 | 12.196 | BB | 0.3093 | 1270.20874 | 62.45296 | 49.4311 |
| 2 | 17.718 | BB | 0.4698 | 1299.44507 | 42.60729 | 50.5689 |

Totals : 2569.65381 105.06025

***tert*-Butyl (+)-(*3R,4R*)-4-(4-iodophenyl)-3-(quinolin-8-ylcarbamoyl)piperidine-1-carboxylate ((+)-6b**

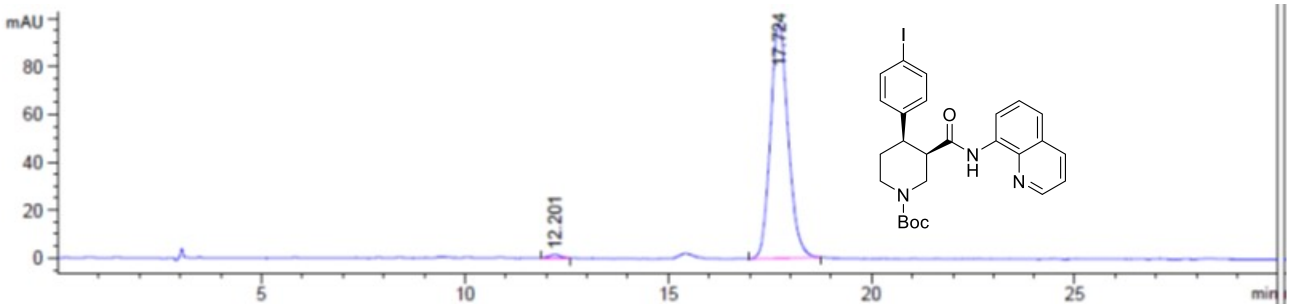


Signal 3: DAD1 C, Sig=210,10 Ref=off

| Peak # | RetTime [min] | Type | Width [min] | Area [mAU*s] | Height [mAU] | Area % |
|---|---|---|---|---|---|---|
| 1 | 12.201 | BB | 0.2287 | 26.94304 | 1.45051 | 0.8982 |
| 2 | 17.724 | BB | 0.4677 | 2972.60156 | 98.05776 | 99.1018 |

Totals : 2999.54460 99.50827

***ee* = 98.2%**

**tert-Butyl trans-(±)-4-(4-iodophenyl)-3-(quinolin-8-ylcarbamoyl)piperidine-1-carboxylate ((±)-S3b)**

**Conditions**: Chiralpak IA 3-column, 85:15 *n*-hexane:*i*-PrOH, flow rate: 1 mL·min$^{-1}$, 35 ºC, UV detection wavelength: 254.1 nm. Retention times: 9.4 min (3*R*,4*S* enantiomer), 13.3 min (3*S*,4*R* enantiomer).

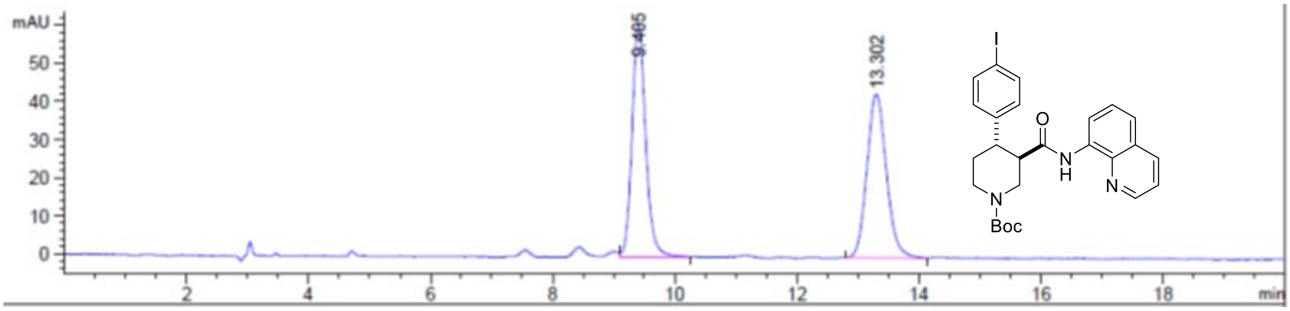


Signal 2: DAD1 B, Sig=254,10 Ref=off

| Peak # | RetTime [min] | Type | Width [min] | Area [mAU*s] | Height [mAU] | Area % |
|---|---|---|---|---|---|---|
| 1 | 9.405 | VB | 0.2368 | 464.95227 | 30.00297 | 49.8782 |
| 2 | 13.301 | BB | 0.3431 | 467.22318 | 21.03599 | 50.1218 |
| Totals : | | | | 932.17545 | 51.03896 | |

**tert-butyl (–)-(3R,4S)-4-(4-iodophenyl)-3-(quinolin-8-ylcarbamoyl)piperidine-1-carboxylate ((–)-S3b)**

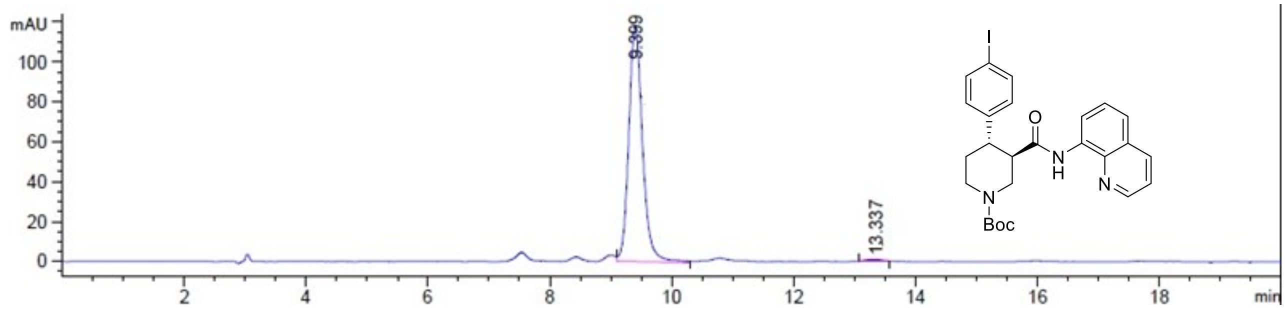


Signal 2: DAD1 B, Sig=254,10 Ref=off

| Peak # | RetTime [min] | Type | Width [min] | Area [mAU*s] | Height [mAU] | Area % |
|---|---|---|---|---|---|---|
| 1 | 9.400 | MM | 0.2584 | 902.97003 | 58.23014 | 99.0491 |
| 2 | 13.314 | MM | 0.2976 | 8.66880 | 4.85432e-1 | 0.9509 |
| Totals : | | | | 911.63883 | 58.71558 | |

***ee* = 98.1%**

***tert*-Butyl (+)-(*3S,4R*)-4-(4-iodophenyl)-3-(quinolin-8-ylcarbamoyl)piperidine-1-carboxylate ((+)-7b)**

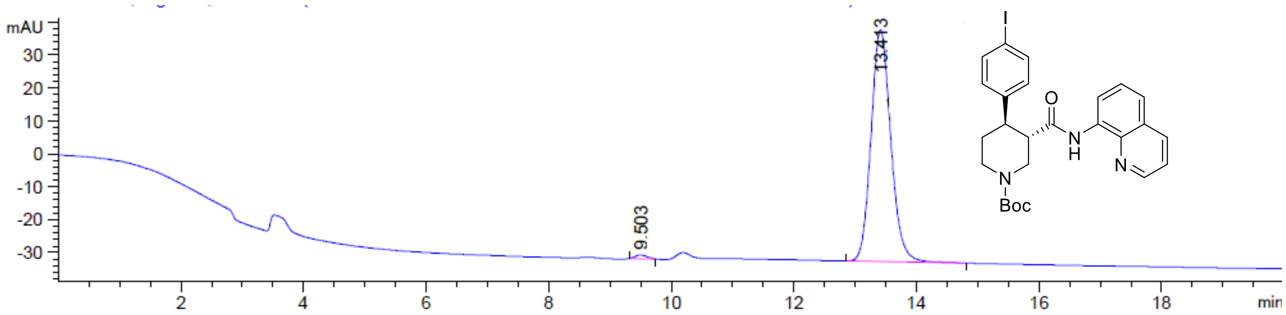


Signal 2: DAD1 B, Sig=254,10 Ref=off

| Peak # | RetTime [min] | Type | Width [min] | Area [mAU*s] | Height [mAU] | Area % |
|---|---|---|---|---|---|---|
| 1 | 9.503 | MM | 0.2272 | 15.83589 | 1.16162 | 0.9922 |
| 2 | 13.413 | MP | 0.3735 | 1580.24548 | 70.51000 | 99.0078 |

| | | |
|---|---|---|
| Totals : | 1596.08137 | 71.67163 |

***ee* = 98.0%**

**Scalemic Mixture of Enantioenriched I-Piperidines (+)-S4b and (–)-8b**

**Conditions**: Chiralpak ID 3-column, 90:10 *n*-hexane:*i*-PrOH, flow rate: 1 mL·min$^{-1}$, 35 ºC, UV detection wavelength: 230.1 nm. Retention times: 6.7 min (3*R*,4*S* enantiomer), 7.4 min (3*S*,4*R* enantiomer).

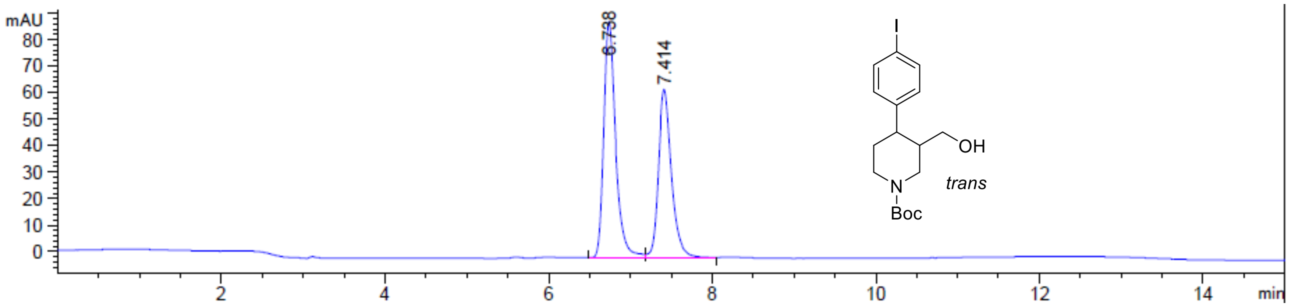


Signal 4: DAD1 D, Sig=230,10 Ref=off

| Peak # | RetTime [min] | Type | Width [min] | Area [mAU*s] | Height [mAU] | Area % |
|---|---|---|---|---|---|---|
| 1 | 6.738 | BV | 0.1527 | 891.19653 | 89.06178 | 56.0128 |
| 2 | 7.414 | VB | 0.1667 | 699.86224 | 63.44310 | 43.9872 |
| Totals : | | | | 1591.05878 | 152.50488 | |

***tert*-Butyl (+)-(*3R,4S*)-4-(4-iodophenyl)-3-(hydroxymethyl)piperidine-1-carboxylate ((+)-S4b)**

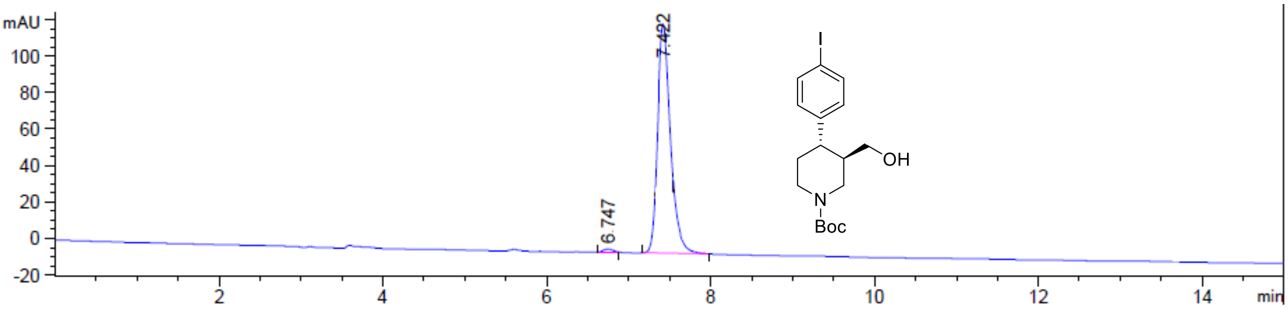


Signal 4: DAD1 D, Sig=230,10 Ref=off

| Peak # | RetTime [min] | Type | Width [min] | Area [mAU*s] | Height [mAU] | Area % |
|---|---|---|---|---|---|---|
| 1 | 6.747 | PP | 0.1304 | 13.28141 | 1.69711 | 0.9559 |
| 2 | 7.422 | PP | 0.1825 | 1376.08704 | 125.65760 | 99.0441 |
| Totals : | | | | 1389.36845 | 127.35471 | |

***ee* = 98.1%**

**_tert_-Butyl (–)-(_3S,4R_)-4-(4-iodophenyl)-3-(hydroxymethyl)piperidine-1-carboxylate ((–)-8b)**

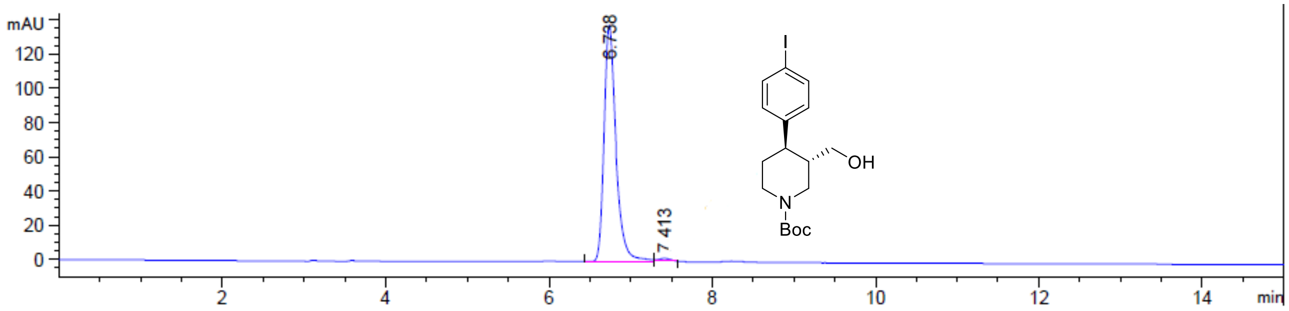


Signal 4: DAD1 D, Sig=230,10 Ref=off

| Peak # | RetTime [min] | Type | Width [min] | Area [mAU*s] | Height [mAU] | Area % |
|---|---|---|---|---|---|---|
| 1 | 6.738 | MM | 0.1672 | 1386.67920 | 138.19916 | 99.0161 |
| 2 | 7.413 | MP | 0.1557 | 13.77859 | 1.47463 | 0.9839 |
| Totals : | | | | 1400.45779 | 139.67379 | |

***ee* = 98.0%**

#### Experimental Details and Characterization Data

Synthesis of Br-analogue of (–)-paroxetine (compounds **(±)-S2a**, **(±)-S3a**, **(+)-6a**, **(+)-7a**, **(–)-S3a**, **(–)-8a**, **(+)-S4a**, **(–)-9a** and **2** · HCl)

***tert*-Butyl *cis*-(±)-4-(4-bromophenyl)-3-(quinolin-8-ylcarbamoyl)piperidine-1-carboxylate ((±)-S2a)** and ***tert*-butyl *trans*-(±)-4-(4-bromophenyl)-3-(quinolin-8-ylcarbamoyl)piperidine-1-carboxylate ((±)-S3a):**

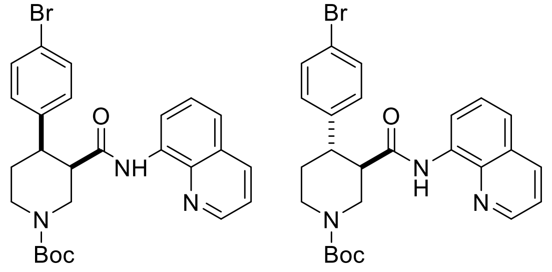


A reaction tube was charged with $K_2CO_3$ (69.1 mg, 0.50 mmol, 1 equiv), flame-dried, and allowed to cool under argon. *tert*-Butyl (±)-3-(quinoline-8-ylcarbamoyl)piperidine-1-carboxylate (**(±)-S1**) (178 mg, 0.50 mmol, 1 equiv), 4-bromoiodobenzene (424 mg, 1.50 mmol, 3 equiv), $Pd(OAc)_2$ (5.60 mg, 25.0 μmol, 5 mol %) and PivOH (51.2 mg, 0.50 mmol, 1 equiv) were added sequentially. The reaction vessel was sealed with an aluminum cap (with molded butyl/PTFE septa) and purged with argon, then anhydrous $PhCF_3$ (500 μL, 1.0 M) was added by syringe. The reaction tube was then placed in a preheated oil bath and stirred at 110 ºC for 18 h. The reaction mixture was allowed to cool to rt and EtOAc (10 mL) was added. The resulting mixture was filtered through a pad of Celite®, eluting with further EtOAc (2 × 10 mL). The solvent was removed under reduced pressure, and the crude material was purified by flash column chromatography (0% to 5% $CH_3CN/CH_2Cl_2$). The product containing fractions were combined and the solvent was removed under reduced pressure. $Et_2O$ (5 mL) and pentane (5 mL) were added and the solvent was removed under reduced pressure to afford the minor product *tert*–butyl *trans*-(±)-4-(4-bromophenyl)-3-(quinolin-8-ylcarbamoyl) piperidine-1-carboxylate **(±)-S3a** as a pale yellow solid (34.5 mg, 14%) followed by the major product *tert*-butyl *cis*-(±)-4-(4-bromophenyl)-3-(quinolin-8-ylcarbamoyl)piperidine-1-carboxylate **(±)-S2a** as an off-white solid (87.2 mg, 34%).

Major (**(±)-S2a**):
$R_f$ 0.31 (5% $CH_3CN/CH_2Cl_2$);
mp = 81–86 °C (from $Et_2O$/pentane);
$\nu_{max}$ (film)/cm$^{-1}$ 3343 (NH), 2859, 1684 (C=O), 1521, 1484, 1423, 1364, 1323, 1245, 1163, 1006, 827, 790, 757;
$^1$H NMR (500 MHz, $(CD_3)_2SO$, 373 K) δ 9.75 (br s, 1 H, NH), 8.83 (dd, *J* = 4.2, 1.7 Hz, 1 H, $HC_{Ar}$), 8.45 (dd, *J* = 7.7, 1.4 Hz, 1 H, $HC_{Ar}$), 8.31 (dd, *J* = 8.3, 1.7 Hz, 1 H, $HC_{Ar}$), 7.60–7.53 (m, 2 H, $HC_{Ar}$), 7.48 (t, *J* = 7.9 Hz, 1 H, $HC_{Ar}$), 7.40–7.34 (m, 2 H, $HC_{Ar}$), 7.33–7.26 (m, 2 H, $HC_{Ar}$), 4.42 (ddd, *J* = 14.8, 3.6, 1.7 Hz, 1 H, NC*H*HCHCO), 4.25 (ddt, *J* = 13.1, 4.6, 2.3 Hz, 1 H, NC*H*HCH$_2$), 3.36–3.28 (m, 2 H, NCH*H*CHCO, CHCO), 3.16 (dt, *J* = 12.2, 4.0 Hz, 1 H, CHAr), 3.02–2.92 (m, 1 H, NCH*H*CH$_2$), 2.68 (qd, *J* = 12.4, 4.7 Hz, 1 H, NCH$_2$C*H*H), 1.72 (dq, *J* = 12.9, 3.2 Hz, 1 H, NCH$_2$CH*H*), 1.25 (s, 9 H, $C(CH_3)_3$);
$^{13}$C NMR (126 MHz, $(CD_3)_2SO$, 373 K) δ 169.8 (C=O amide), 153.4 (C=O carbamate), 147.9 ($C_{Ar}$), 142.0 ($C_{Ar}$ quat), 137.6 ($C_{Ar}$ quat), 135.7 ($C_{Ar}$), 133.9 ($C_{Ar}$ quat), 130.3 (2 × $C_{Ar}$), 129.1 (2 × $C_{Ar}$), 127.2 ($C_{Ar}$ quat), 126.1 ($C_{Ar}$), 121.2 ($C_{Ar}$), 120.8 ($C_{Ar}$), 118.7 (Br$C_{Ar}$ quat), 115.7 ($C_{Ar}$), 77.9 (*C*(CH$_3$)$_3$), 46.2 (N*C*H$_2$CHCO), 45.6 (*C*HCO), 42.9 (N*C*H$_2$CH$_2$), 41.7 (*C*HAr), 27.4 (C(*C*H$_3$)$_3$), 25.0 (NCH$_2$*C*H$_2$);
HRMS (ESI$^+$) *m/z* Calculated for $C_{26}H_{29}N_3O_3{}^{79}Br$ [M+H] 510.1392; Found 510.1386.

SMILES:
O=C([C@H]1CN(C(OC(C)(C)C)=O)CC[C@H]1C2=CC=C(Br)C=C2)NC3=C(N=CC=C4)C4=CC=C3

*InChI=1S/C26H28BrN3O3/c1-26(2,3)33-25(32)30-15-13-20(17-9-11-19(27)12-10-17)21(16-30)24(31)29-22-8-4-6-18-7-5-14-28-23(18)22/h4-12,14,20-21H,13,15-16H2,1-3H3,(H,29,31)/t20-,21-/m0/s1*

Minor (**(±)-S3a**):
*R*f 0.41 (5% $CH_3CN/CH_2Cl_2$);
mp = 77–83 °C (from $Et_2O$/pentane);
$\nu_{max}$ (film)/cm$^{-1}$ 3340 (NH), 2926, 1677 (C=O), 1521, 1484, 1424, 1323, 1230, 1156, 1126, 999, 824, 757;
$^1$H NMR (500 MHz, $(CD_3)_2SO$, 373 K) δ 9.73 (br s, 1 H, NH), 8.85 (dd, *J* = 4.2, 1.7 Hz, 1 H, $HC_{Ar}$), 8.39 (dd, *J* = 7.7, 1.4 Hz, 1 H, $HC_{Ar}$), 8.31 (dd, *J* = 8.3, 1.7 Hz, 1 H, $HC_{Ar}$), 7.61–7.54 (m, 2 H, $HC_{Ar}$), 7.47 (t, *J* = 8.0 Hz, 1 H, $HC_{Ar}$), 7.39–7.32 (m, 2 H, $HC_{Ar}$), 7.34–7.28 (m, 2 H, $HC_{Ar}$), 4.36 (ddd, *J* = 12.9, 3.7, 1.8 Hz, 1 H, NC*H*HCHCO), 4.13 (ddt, *J* = 13.3, 4.3, 2.2 Hz, 1 H, NC*H*HCH$_2$), 3.18–3.00 (m, 3 H, NCH*H*CHCO, CHCO, CHAr), 2.99–2.90 (m, 1 H, NCH*H*CH$_2$), 1.81 (dq, *J* = 12.9, 2.8 Hz, 1 H, NCH$_2$C*H*H), 1.66 (qd, *J* = 12.8, 4.6 Hz, 1 H, NCH$_2$CH*H*), 1.49 (s, 9 H, $C(CH_3)_3$);
$^{13}$C NMR (126 MHz, $(CD_3)_2SO$, 373 K) δ 169.8 (C=O amide), 153.4 (C=O carbamate), 148.0 ($C_{Ar}$), 142.4 ($C_{Ar}$ quat), 137.7 ($C_{Ar}$ quat), 135.7 ($C_{Ar}$), 133.5 ($C_{Ar}$ quat), 130.6 (2 × $C_{Ar}$), 129.1 (2 × $C_{Ar}$), 127.2 ($C_{Ar}$ quat), 126.1 ($C_{Ar}$), 121.4 ($C_{Ar}$), 121.3 ($C_{Ar}$), 118.9 (Br$C_{Ar}$ quat), 116.3 ($C_{Ar}$), 78.6 (*C*(CH$_3$)$_3$), 49.2 (*C*HCO), 46.2 (N*C*H$_2$CHCO), 43.9 (*C*HAr), 43.3 (N*C*H$_2$CH$_2$), 32.0 (NCH$_2$*C*H$_2$), 27.7 (C(*C*H$_3$)$_3$);
HRMS (ESI$^+$) *m/z* Calculated for $C_{26}H_{29}N_3O_3{}^{79}Br$ [M+H] 510.1392; Found 510.1382.

SMILES:
O=C([C@@H]1CN(C(OC(C)(C)C)=O)CC[C@H]1C2=CC=C(Br)C=C2)NC3=C(N=CC=C4)C4=CC=C3

InChI=1S/C26H28BrN3O3/c1-26(2,3)33-25(32)30-15-13-20(17-9-11-19(27)12-10-17)21(16-30)24(31)29-22-8-4-6-18-7-5-14-28-23(18)22/h4-12,14,20-21H,13,15-16H2,1-3H3,(H,29,31)/t20-,21+/m0/s1

***tert*-Butyl (+)-(*3R,4R*)-4-(4-bromophenyl)-3-(quinolin-8-ylcarbamoyl)piperidine-1-carboxylate ((+)-6a)** and ***tert*-butyl (–)-(*3R,4S*)-4-(4-bromophenyl)-3-(quinolin-8-ylcarbamoyl)piperidine-1-carboxylate ((–)-S3a)**:

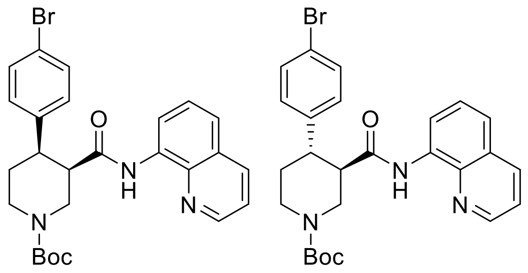


A large microwave vial (10–20 mL recommended volume) was charged with $K_2CO_3$ (553 mg, 4.0 mmol, 1 equiv), flame-dried, and allowed to cool under argon. *tert*-Butyl (*R*)-3-(quinolin-8-ylcarbamoyl)piperidine-1-carboxylate **(–)-5** (1.42 g, 4.0 mmol, 1 equiv), 4-bromoiodobenzene (3.40 g, 12.0 mmol, 3 equiv), $Pd(OAc)_2$ (45.1 mg, 0.2 mmol, 5 mol %) and PivOH (409 mg, 4.0 mmol, 1 equiv) were added sequentially. The reaction vessel was sealed with an aluminum cap (with molded butyl/PTFE septa) and purged with argon, then anhydrous $PhCF_3$ (2.0 mL, 2.00 M) was added by syringe. The reaction tube was then placed in a preheated oil bath and stirred at 110 ºC for 18 h. The reaction mixture was then allowed to cool to rt and EtOAc (20 mL) was added. The resulting mixture was filtered through a pad of Celite®, eluting with further EtOAc (2 × 50 mL). The solvent was removed under reduced pressure. The reaction mixture was purified by two consecutive chromatographic separations: one (0% to 5% $CH_3CN/CH_2Cl_2$) to isolate the minor *trans*-product *tert*-butyl (–)-(*3R,4S*)-4-(4-bromophenyl)-3-(quinolin-8-ylcarbamoyl)piperidine-1-carboxylate **(–)-S3a** followed by a second (10% to 15% acetone/pentane) to isolate the major *cis*-product *tert*-butyl (+)-(*3R,4R*)-4-(4-bromophenyl)-3-(quinolin-8-ylcarbamoyl)piperidine-1-carboxylate **(+)-6a**. The product containing fractions were combined and the solvent was removed under reduced pressure. $Et_2O$ (20 mL) and pentane (20 mL) were added and the solvent was removed under reduced pressure to afford the minor *trans*-product **(–)-S3a** as a pale yellow solid (371 mg, 18%, 98.0% *ee*) and the major *cis*-product **(+)-6a** as a white solid (730 mg, 36%, 98.2% *ee*).

Major (**(+)-6a**):
$[\alpha]_D^{23}$ + 15.4 (*c* 1.3, $CHCl_3$).

Characterization data identical to that reported for racemic *cis*-piperidine **(±)-S2a** (see S17).

**HPLC Conditions**: Chiralpak IA 3-column, 85:15 *n*-hexane:*i*-PrOH, flow rate: 1 mL·min$^{-1}$, 35 ºC, UV detection wavelength: 210.4 nm. Retention times: 11.9 min (3*S*,4*S* enantiomer), 17.3 min (3*R*,4*R* enantiomer).

SMILES:
O=C([C@H]1CN(C(OC(C)(C)C)=O)CC[C@H]1C2=CC=C(Br)C=C2)NC3=C(N=CC=C4)C4=CC=C3
InChI=1S/C26H28BrN3O3/c1-26(2,3)33-25(32)30-15-13-20(17-9-11-19(27)12-10-17)21(16-30)24(31)29-22-8-4-6-18-7-5-14-28-23(18)22/h4-12,14,20-21H,13,15-16H2,1-3H3,(H,29,31)/t20-,21-/m0/s1

Minor (**(–)-S3a**):
$[\alpha]_D^{23}$ – 35.4 (*c* 1.3, $CHCl_3$).
Characterization data identical to that reported for racemic *trans*-piperidine **(±)-S3a** (see S17).

**HPLC Conditions**: Chiralpak IA 3-column, 85:15 *n*-hexane:*i*-PrOH, flow rate: 1 mL·min$^{-1}$, 35 ºC, UV detection wavelength: 254.1 nm. Retention times: 9.1 min (3*R*,4*S* enantiomer), 12.2 min (3*S*,4*R* enantiomer).

SMILES:
O=C([C@H]1CN(C(OC(C)(C)C)=O)CC[C@@H]1C2=CC=C(Br)C=C2)NC3=C(N=CC=C4)C4=CC=C3

InChI=1S/C26H28BrN3O3/c1-26(2,3)33-25(32)30-15-13-20(17-9-11-19(27)12-10-17)21(16-30)24(31)29-22-8-4-6-18-7-5-14-28-23(18)22/h4-12,14,20-21H,13,15-16H2,1-3H3,(H,29,31)/t20-,21+/m1/s1

***tert*-Butyl (+)-(*3S,4R*)-4-(4-bromophenyl)-3-(quinolin-8-ylcarbamoyl)piperidine-1-carboxylate ((+)-7a)**

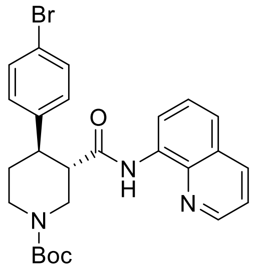

A flame-dried reaction tube was charged with *cis*-3,4-disubstituted piperidine **(+)-6a** (662 mg, 1.30 mmol, 1 equiv) and 1,8-diazabicyclo(5.4.0)undec-7-ene (DBU, 600 μL, 3.90 mmol, 3 equiv). The reaction vessel was sealed with an aluminum cap (with molded butyl/PTFE septa) and purged with argon, then anhydrous toluene (1.30 mL, 1.0 M) was added by syringe. The reaction tube was then placed in a preheated oil bath and stirred at 110 ºC for 24 h. The reaction mixture was then allowed to cool to rt and $CH_2Cl_2$ (5 mL) and sat. aq. $NH_4Cl$ (5 mL) were added. The phases were separated, and the aqueous layer was extracted with $CH_2Cl_2$ (3 × 10 mL). The combined organic extracts were dried over $Na_2SO_4$ and filtered. The solvent was removed under reduced pressure. The reaction mixture was purified by flash column chromatography (15% acetone/pentane). The product containing fractions were combined and the solvent was removed under reduced pressure. $Et_2O$ (10 mL) and pentane (10 mL) were added and the solvent was removed under reduced pressure to afford amide *tert*-butyl (+)-(*3S,4R*)-4-(4-bromophenyl)-3-(quinolin-8-ylcarbamoyl) piperidine-1-carboxylate **(+)-7a** as a white solid (621 mg, 94%, 98.4% *ee*).

$[\alpha]_D^{23}$ + 52.0 (*c* 1.0, $CHCl_3$).
Characterization data identical to that reported for racemic *trans*-piperidine **(±)-S3a** (see S17).

**HPLC Conditions**: Chiralpak IA 3-column, 85:15 *n*-hexane:*i*-PrOH, flow rate: 1 mL·min$^{-1}$, 35 ºC, UV detection wavelength: 254.1 nm. Retention times: 9.1 min (3*R*,4*S* enantiomer), 12.2 min (3*S*,4*R* enantiomer).

SMILES:
O=C([C@@H]1CN(C(OC(C)(C)C)=O)CC[C@H]1C2=CC=C(Br)C=C2)NC3=C(N=CC=C4)C4=CC=C3

InChI=1S/C26H28BrN3O3/c1-26(2,3)33-25(32)30-15-13-20(17-9-11-19(27)12-10-17)21(16-30)24(31)29-22-8-4-6-18-7-5-14-28-23(18)22/h4-12,14,20-21H,13,15-16H2,1-3H3,(H,29,31)/t20-,21+/m0/s1

***tert*-Butyl (+)-(*3R,4S*)-4-(4-bromophenyl)-3-(hydroxymethyl)piperidine-1-carboxylate ((+)-S4a)**

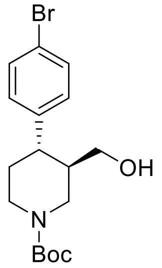


A flame-dried reaction tube was charged with amide **(–)-S3a** (102 mg, 0.20 mmol, 1 equiv), followed by di-*tert*-butyl dicarbonate ($Boc_2O$, 175 mg, 0.80 mmol, 4 equiv) and 4-(dimethylamino)pyridine (DMAP, 4.9 mg, 0.04 mmol, 20 mol %). The reaction vessel was sealed with an aluminum cap (with molded butyl septa) and purged with argon, then anhydrous MeCN (400 μL, 0.5 M) was added by syringe. The mixture was then stirred at 35 ºC for 22 h. The reaction mixture was then allowed to cool to rt and sat. aq. $NH_4Cl$ (1 mL) and $CH_2Cl_2$ (1 mL) were added. The phases were separated, and the aqueous layer was extracted with $CH_2Cl_2$ (3 × 5 mL). The combined organic extracts were dried over $Na_2SO_4$ and filtered. The solvent was removed under reduced pressure to afford the crude *N*-Boc protected piperidine derivative.

This crude was solubilized in anhydrous THF (800 μL, 0.2 M) and the resulting solution was added dropwise to a suspension of $LiAlH_4$ (15.2 mg, 0.40 mmol, 2 equiv) in anhydrous THF (200 μL, 2.0 M) at 0 °C under argon atmosphere. The mixture was then stirred at 20 °C for 30 min. The reaction mixture was then quenched by slow addition of sat. aq. $NH_4Cl$ (2 mL) at 0 °C and stirred at rt for 30 min. The resulting suspension was filtered through a pad of Celite®, eluting with EtOAc (3 × 5 mL). The phases were separated, and the aqueous layer was extracted with EtOAc (3 × 5 mL). The combined organic extracts were dried over $Na_2SO_4$ and filtered. The solvent was removed under reduced pressure. Purification by flash column chromatography (10% to 20% acetone/hexane) afforded primary alcohol **(+)-S4a** as a yellow solid (52.0 mg, 70% over 2 steps, 98.1% *ee*).

$[\alpha]_D^{23}$ + 5.0 (*c* 0.8, $CHCl_3$).
$R_f$ 0.21 (20% acetone/hexane);
mp = 49–54 °C;
$\nu_{max}$ (film)/cm$^{-1}$ 3407 (OH), 2922, 1662 (C=O), 1476, 1424, 1230, 1159, 1129, 1059, 1006, 816, 769;
$^1$H NMR (400 MHz, $CDCl_3$, 298 K) δ 7.47–7.40 (m, 2 H, $HC_{Ar}$), 7.11–7.05 (m, 2 H, $HC_{Ar}$), 4.36 (br d, *J* = 13.2 Hz, 1 H, NC*H*HCHCH$_2$OH), 4.20 (br s, 1 H, NC*H*HCH$_2$), 3.43 (dd, *J* = 11.0, 3.1 Hz, 1 H, C*H*HOH), 3.26 (dd, *J* = 11.0, 6.4 Hz, 1 H, CH*H*OH), 2.87–2.62 (m, 2 H, NCH*H*CHCH$_2$OH, NCH*H*CH$_2$), 2.59–2.47 (m, 1 H, CHAr), 1.88–1.59 (m, 4 H, C*H*CH$_2$OH, NCH$_2$C*H*$_2$, OH), 1.49 (s, 9 H, C(CH$_3$)$_3$);
$^{13}$C NMR (101 MHz, $CDCl_3$, 298 K, observed as a mixture of rotamers) δ 154.8 (C=O), 142.8 ($C_{Ar}$ quat), 131.8 (2 × $C_{Ar}$), 129.1 (2 × $C_{Ar}$), 120.3 (Br$C_{Ar}$ quat), 79.7 (*C*(CH$_3$)$_3$), 62.9 (*C*H$_2$OH), 46.5 (br m, N*C*H$_2$CHCH$_2$OH), 44.2 and 43.6 (N*C*H$_2$CH$_2$, *C*HAr, *C*HCH$_2$OH), 33.8 (NCH$_2$*C*H$_2$), 28.5 (C(*C*H$_3$)$_3$);
HRMS (ESI$^+$) *m/z* Calculated for $C_{19}H_{27}N_2O_3Na^{79}Br$ [M+CH$_3$CN+Na Adduct] 433.1103; Found 433.1110.

**HPLC Conditions**: Chiralpak ID 3-column, 90:10 *n*-hexane:*i*-PrOH, flow rate: 1 mL·min$^{-1}$, 35 ºC, UV detection wavelength: 210.4 nm. Retention times: 8.0 min (3*R*,4*S* enantiomer), 8.6 min (3*S*,4*R* enantiomer).

SMILES: BrC1=CC=C([C@@H]2[C@@H](CO)CN(C(OC(C)(C)C)=O)CC2)C=C1

InChI=1S/C17H24BrNO3/c1-17(2,3)22-16(21)19-9-8-15(13(10-19)11-20)12-4-6-14(18)7-5-12/h4-7,13,15,20H,8-11H2,1-3H3/t13-,15-/m1/s1

***tert*-Butyl (–)-(3*S*,4*R*)-4-(4-bromophenyl)-3-(hydroxymethyl)piperidine-1-carboxylate ((–)-8a)**

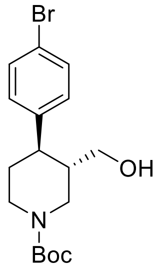

A flame-dried round-bottom flask was charged with amide **(+)-7a** (565 mg, 1.11 mmol, 1 equiv), followed by di-*tert*-butyl dicarbonate ($Boc_2O$, 969 mg, 4.44 mmol, 4 equiv) and 4-(dimethylamino)pyridine (DMAP, 26.9 mg, 0.22 mmol, 20 mol %). The reaction vessel was sealed with an aluminum cap (with molded butyl septa) and purged with argon, then anhydrous MeCN (3.7 mL) and anhydrous $CH_2Cl_2$ (0.5 mL) were added by syringe. The mixture (0.3 M) was then stirred at 35 ºC for 22 h. The reaction mixture was then allowed to cool to rt and sat. aq. $NH_4Cl$ (5 mL) and $CH_2Cl_2$ (5 mL) were added. The phases were separated, and the aqueous layer was extracted with $CH_2Cl_2$ (3 × 10 mL). The combined organic extracts were dried over $Na_2SO_4$ and filtered. The solvent was removed under reduced pressure to afford the crude *N*-Boc protected piperidine derivative.

This crude was solubilized in anhydrous THF (3.5 mL, 0.3 M) and the resulting solution was added dropwise to a suspension of $LiAlH_4$ (84.2 mg, 2.22 mmol, 2 equiv) in anhydrous THF (2.0 mL, 1.0 M) at 0 °C under argon atmosphere. The mixture was then stirred at 20 °C for 30 min. The reaction mixture was then quenched by slow addition of sat. aq. $NH_4Cl$ (5 mL) at 0 °C and stirred at rt for 30 min. The resulting suspension was filtered through a pad of Celite®, eluting with EtOAc (3 × 10 mL). The phases were separated, and the aqueous layer was extracted with EtOAc (3 × 10 mL). The combined organic extracts were dried over $Na_2SO_4$ and filtered. The solvent was removed under reduced pressure. Purification by flash column chromatography (10% to 20% acetone/hexane) afforded primary alcohol **(–)-8a** as a white solid (316 mg, 77% over 2 steps, 98.1% *ee*).

$[\alpha]_D^{23}$ – 8.0 (*c* 1.0, $CHCl_3$).

Characterization data identical to that reported for enantiomeric alcohol **(+)-S4a** (see S20).

**HPLC Conditions**: Chiralpak ID 3-column, 90:10 *n*-hexane:*i*-PrOH, flow rate: 1 mL·min$^{-1}$, 35 ºC, UV detection wavelength: 210.4 nm. Retention times: 8.0 min (3*R*,4*S* enantiomer), 8.6 min (3*S*,4*R* enantiomer).

SMILES: BrC1=CC=C([C@H]2[C@H](CO)CN(C(OC(C)(C)C)=O)CC2)C=C1

InChI=1S/C17H24BrNO3/c1-17(2,3)22-16(21)19-9-8-15(13(10-19)11-20)12-4-6-14(18)7-5-12/h4-7,13,15,20H,8-11H2,1-3H3/t13-,15-/m0/s1

***tert*-Butyl (3*S*,4*R*)-3-((benzo[*d*][1,3]dioxol-5-yloxy)methyl)-4-(4-bromophenyl)piperidine-1-carboxylate ((–)-9a)**

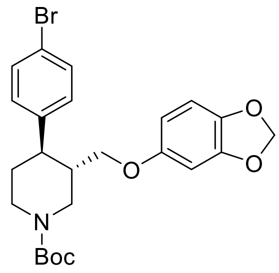

Alcohol **(–)-8a** (280 mg, 0.76 mmol, 1 equiv) and triethylamine (147 μL, 1.10 mmol, 1.4 equiv) were added to a flame-dried round-bottom flask, dissolved in anhydrous $CH_2Cl_2$ (4.0 mL, 0.2 M) and cooled down to 0 °C. Methanesulfonyl chloride (75 μL, 0.97 mmol, 1.3 equiv) was then added by Gilson pipette. After stirring 5 min at 0 °C, the reaction mixture was stirred at 25 °C for 2 h, then diluted with $CH_2Cl_2$ (5 mL) and sat. aq. $NaHCO_3$ (5 mL). The phases were separated, and the aqueous layer was extracted with $CH_2Cl_2$ (3 × 10 mL). The combined organic extracts were dried over $Na_2SO_4$ and filtered. The solvent was removed under reduced pressure to afford the crude mesylated alcohol derivative.

NaH (60% dispersion in mineral oil, 51.8 mg, 1.30 mmol, 1.7 equiv) was added to a solution of sesamol (168 mg, 1.20 mmol, 1.6 equiv) in anhydrous THF (4.0 mL, 0.3 M) at 0 °C. The mixture was then stirred at 25 °C for 1 h. A solution of the crude mesylated alcohol in anhydrous THF (5.0 mL, 0.1 M) was then added dropwise to this suspension. The resulting mixture was stirred at 70 °C for 18 h. The reaction mixture was then quenched by addition of $H_2O$ (5 mL) and diluted with EtOAc (5 mL). The phases were separated, and the aqueous layer was extracted with EtOAc (4 × 10 mL). The combined organic extracts were dried over $Na_2SO_4$ and filtered. The solvent was removed under reduced pressure. Purification by flash column chromatography (5% acetone/pentane) afforded piperidine **(–)-9a** as a white solid (225 mg, 60% over 2 steps).

$[\alpha]_D^{23}$ – 36.0 (*c* 1.0, $CHCl_3$).
$R_f$ 0.20 (5% acetone/pentane);
mp = 53–58 °C;
$\nu_{max}$ (film)/cm$^{-1}$ 2915, 1685 (C=O), 1483, 1424, 1230, 1163, 1129, 1036, 928, 816, 769;
$^1$H NMR (400 MHz, $CDCl_3$, 298 K) δ 7.45–7.38 (m, 2 H, $HC_{Ar}$), 7.10–7.03 (m, 2 H, $HC_{Ar}$), 6.64 (d, *J* = 8.5 Hz, 1 H, $HC_{Ar}$), 6.36 (d, *J* = 2.5 Hz, 1 H, $HC_{Ar}$), 6.14 (dd, *J* = 8.5, 2.5 Hz, 1 H, $HC_{Ar}$), 5.89 (s, 2 H, $OCH_2O$), 4.44 (br s, 1 H, NC*H*HCHCH$_2$OAr), 4.25 (br s, 1 H, NC*H*HCH$_2$), 3.61 (dd, *J* = 9.4, 2.8 Hz, 1 H, C*H*HOAr), 3.45 (dd, *J* = 9.4, 6.4 Hz, 1 H, CH*H*OAr), 2.92–2.73 (br m, 2 H, NCH*H*CHCH$_2$OAr, NCH*H*CH$_2$), 2.67 (td, *J* = 11.7, 3.9 Hz, 1 H, CHAr), 2.08–1.97 (br m, 1 H, C*H*CH$_2$OAr), 1.85–1.77 (br m, 1 H, NCH$_2$C*H*H), 1.72 (td, *J* = 12.6, 4.3 Hz, 1 H, NCH$_2$CH*H*), 1.50 (s, 9 H, $C(CH_3)_3$);
$^{13}$C NMR (101 MHz, $CDCl_3$, 298 K) δ 154.7 (C=O), 154.2 ($OC_{Ar}$ quat), 148.1 ($OC_{Ar}$ quat), 142.4 ($C_{Ar}$ quat), 141.7 ($OC_{Ar}$ quat), 131.8 (2 × $C_{Ar}$), 129.1 (2 × $C_{Ar}$), 120.4 ($BrC_{Ar}$ quat), 107.8 ($C_{Ar}$), 105.5 ($C_{Ar}$), 101.1 ($OCH_2O$), 98.0 ($C_{Ar}$), 79.7 (*C*(CH$_3$)$_3$), 68.7 (*C*H$_2$OAr), 47.3 (br m, N*C*H$_2$CHCH$_2$OAr), 44.2 (N*C*H$_2$CH$_2$, *C*HAr), 41.7 (*C*HCH$_2$OAr), 33.7 (NCH$_2$*C*H$_2$), 28.4 (C(*C*H$_3$)$_3$);
HRMS (ESI$^+$) *m/z* Calculated for $C_{24}H_{29}NO_5{}^{79}Br$ [M+H] 490.1229; Found 490.1240.

SMILES: BrC1=CC=C([C@H]2[C@H](COC3=CC(OCO4)=C4C=C3)CN(C(OC(C)(C)C)=O)CC2)C=C1

InChI=1S/C24H28BrNO5/c1-24(2,3)31-23(27)26-11-10-20(16-4-6-18(25)7-5-16)17(13-26)14-28-19-8-9-21-22(12-19)30-15-29-21/h4-9,12,17,20H,10-11,13-15H2,1-3H3/t17-,20-/m0/s1

**(3*S*,4*R*)-3-((Benzo[d][1,3]dioxol-5-yloxy)methyl)-4-(4-bromophenyl)piperidine-1-ium chloride (2 · HCl)**

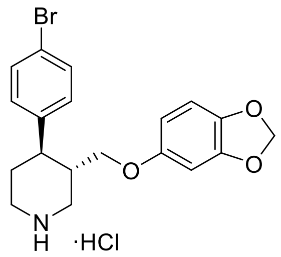


4 N HCl in 1,4-dioxane (500 μL, 2.00 mmol, 10 equiv) was added to a solution of *N*-Boc protected piperidine **(–)-9a** (98.1 mg, 0.20 mmol, 1 equiv) in 1,4-dioxane (500 μL, 0.4 M) at 0 °C under air. The solution was stirred at 25 ºC for 18 h, then an ice-cold 1:1 mixture of $Et_2O$/pentane (1 mL) was added and formation of a solid precipitate was observed. This was filtered and washed with further ice-cold $Et_2O$/pentane mixture (2 × 5 mL). The solid precipitate was dried under reduced pressure to afford (3*S*,4*R*)-3-((benzo[d][1,3]dioxol-5-yloxy)methyl)-4-(4-bromophenyl) piperidine-1-ium chloride **2** · HCl as an off-white solid (73.5 mg, 86%).

$[\alpha]_D^{23}$ – 82.0 (*c* 1.0, MeOH);
mp = 206–209 ºC;
$\nu_{max}$ (film)/cm$^{-1}$ 3317 (NH), 2926, 2687, 1484, 1182, 1103, 1033, 932, 846, 813, 787;
$^1$H NMR (400 MHz, $CD_3OD$, 298 K) δ 7.50–7.44 (m, 2 H, $HC_{Ar}$), 7.24–7.18 (m, 2 H, $HC_{Ar}$), 6.63 (d, *J* = 8.4 Hz, 1 H, $HC_{Ar}$), 6.39 (d, *J* = 2.5 Hz, 1 H, $HC_{Ar}$), 6.18 (dd, *J* = 8.5, 2.5 Hz, 1 H, $HC_{Ar}$), 5.87–5.84 (m, 2 H, $OCH_2O$), 3.71–3.62 (m, 2 H, C*H*HOAr, NC*H*HCHCH$_2$OAr), 3.59–3.49 (m, 2 H, CH*H*OAr, NC*H*HCH$_2$), 2.21–2.11 (m, 2 H, NCH*H*CHCH$_2$OAr, NCH*H*CH$_2$), 3.03–2.91 (m, 1 H, CHAr), 2.49–2.37 (m, 1 H, C*H*CH$_2$OAr), 2.10–2.01 (m, 2 H, NCH$_2$C*H*$_2$);
$^{13}$C NMR (101 MHz, $CD_3OD$, 298 K) δ 155.3 ($OC_{Ar}$ quat), 149.7 ($OC_{Ar}$ quat), 143.5 ($C_{Ar}$ quat), 142.4 ($OC_{Ar}$ quat), 133.0 (2 × $C_{Ar}$), 130.5 (2 × $C_{Ar}$), 122.0 ($BrC_{Ar}$ quat), 108.8 ($C_{Ar}$), 106.7 ($C_{Ar}$), 102.5 ($OCH_2O$), 98.9 ($C_{Ar}$), 69.0 (*C*H$_2$OAr), 47.7 (N*C*H$_2$CHCH$_2$OAr), 45.5 (N*C*H$_2$CH$_2$), 42.9 (*C*HAr), 40.6 (*C*HCH$_2$OAr), 31.4 (NCH$_2$*C*H$_2$);

HRMS (ESI$^+$) *m/z* Calculated for $C_{19}H_{21}NO_3{}^{79}Br$ [M–Cl] 390.0705; Found 390.0698.

SMILES: BrC1=CC=C([C@H]2[C@H](COC3=CC(OCO4)=C4C=C3)CNCC2)C=C1.Cl

InChI=1S/C19H20BrNO3.ClH/c20-15-3-1-13(2-4-15)17-7-8-21-10-14(17)11-22-16-5-6-18-19(9-16)24-12-23-18;/h1-6,9,14,17,21H,7-8,10-12H2;1H/t14-,17-;/m0./s1

Synthesis of I-analogue of (–)-paroxetine (compounds **(±)-S2b**, **(±)-S3b**, **(+)-6b**, **(+)-7b**, **(–)-S3b**, **(–)-8b**, **(+)-S4b**, **(–)-9b** and **3** · HCl)

***tert*-Butyl *cis*-(±)-4-(4-iodophenyl)-3-(quinolin-8-ylcarbamoyl)piperidine-1-carboxylate ((±)-S2b)** and ***tert*-butyl *trans*-(±)-4-(4-iodophenyl)-3-(quinolin-8-ylcarbamoyl)piperidine-1-carboxylate ((±)-S3b):**

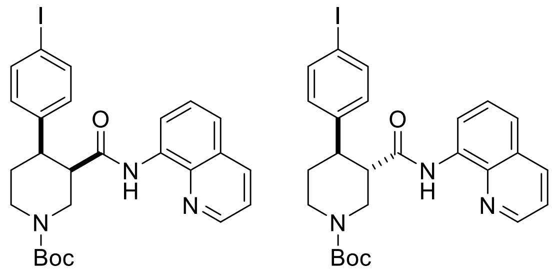


A reaction tube was charged with $K_2CO_3$ (69.1 mg, 0.50 mmol, 1 equiv), flame-dried, and allowed to cool under argon. *tert*-Butyl (±)-3-(quinoline-8-ylcarbamoyl)piperidine-1-carboxylate (**(±)-S1**) (178 mg, 0.50 mmol, 1 equiv), 1,4-diiodobenzene (660 mg, 2.00 mmol, 4 equiv), $Pd(OAc)_2$ (5.60 mg, 25.0 μmol, 5 mol %) and PivOH (51.2 mg, 0.50 mmol, 1 equiv) were added sequentially. The reaction vessel was sealed with an aluminum cap (with molded butyl/PTFE septa) and purged with argon, then anhydrous $PhCF_3$ (500 μL, 1.0 M) was added by syringe. The reaction tube was then placed in a preheated oil bath and stirred at 110 ºC for 18 h. The reaction mixture was allowed to cool to rt and EtOAc (10 mL) was added. The resulting mixture was filtered through a pad of Celite®, eluting with further EtOAc (2 × 10 mL). The solvent was removed under reduced pressure, and the crude material was purified by flash column chromatography (0% to 5% $CH_3CN/CH_2Cl_2$). The product containing fractions were combined and the solvent was removed under reduced pressure. $Et_2O$ (5 mL) and pentane (5 mL) were added and the solvent was removed under reduced pressure to afford the minor product *tert*–butyl *trans*-(±)-4-(4-iodophenyl)-3-(quinolin-8-ylcarbamoyl) piperidine-1-carboxylate **(±)-S3b** as a pale yellow solid (52.2 mg, 19%) followed by the major product *tert*-butyl *cis*-(±)-4-(4-iodophenyl)-3-(quinolin-8-ylcarbamoyl)piperidine-1-carboxylate **(±)-S2b** as a pale yellow solid (97.9 mg, 35%).

Major (**(±)-S2b**):
$R_f$ 0.30 (5% $CH_3CN/CH_2Cl_2$);
mp = 91–95 °C (from $Et_2O$/pentane);
$\nu_{max}$ (film)/cm$^{-1}$ 3343 (NH), 2926, 1685 (C=O), 1521, 1483, 1424, 1364, 1323, 1245, 1159, 1118, 1003, 824, 790, 757;
$^1$H NMR (500 MHz, $(CD_3)_2SO$, 373 K) δ 9.75 (br s, 1 H, NH), 8.83 (dd, *J* = 4.2, 1.7 Hz, 1 H, $HC_{Ar}$), 8.45 (dd, *J* = 7.6, 1.4 Hz, 1 H, $HC_{Ar}$), 8.31 (dd, *J* = 8.3, 1.7 Hz, 1 H, $HC_{Ar}$), 7.60–7.53 (m, 4 H, $HC_{Ar}$), 7.48 (t, *J* = 8.0 Hz, 1 H, $HC_{Ar}$), 7.19–7.12 (m, 2 H, $HC_{Ar}$), 4.42 (ddd, *J* = 14.9, 3.7, 1.8 Hz, 1 H, NC*H*HCHCO), 4.25 (ddt, *J* = 13.2, 4.7, 2.4 Hz, 1 H, NC*H*HCH$_2$), 3.35–3.28 (m, 2 H, NCH*H*CHCO, CHCO), 3.14 (dt, *J* = 12.4, 4.2 Hz, 1 H, CHAr), 3.01–2.92 (m, 1 H, NCH*H*CH$_2$), 2.67 (qd, *J* = 12.4, 4.6 Hz, 1 H, NCH$_2$C*H*H), 1.71 (dq, *J* = 13.0, 3.4 Hz, 1 H, NCH$_2$CH*H*), 1.25 (s, 9 H, C(CH$_3$)$_3$);
$^{13}$C NMR (126 MHz, $(CD_3)_2SO$, 373 K) δ 169.8 (C=O amide), 153.4 (C=O carbamate), 147.9 ($C_{Ar}$), 142.5 ($C_{Ar}$ quat), 137.6 ($C_{Ar}$ quat), 136.3 (2 × $C_{Ar}$), 135.7 ($C_{Ar}$), 133.9 ($C_{Ar}$ quat), 129.3 (2 × $C_{Ar}$), 127.2 ($C_{Ar}$ quat), 126.1 ($C_{Ar}$), 121.2 ($C_{Ar}$), 120.8 ($C_{Ar}$), 115.7 ($C_{Ar}$), 90.9 ($IC_{Ar}$ quat), 77.9 (*C*(CH$_3$)$_3$), 46.2 (N*C*H$_2$CHCO), 45.6 (*C*HCO), 42.9 (N*C*H$_2$CH$_2$), 41.8 (*C*HAr), 27.4 (C(*C*H$_3$)$_3$), 25.0 (NCH$_2$*C*H$_2$);
HRMS (ESI$^+$) *m/z* Calculated for $C_{26}H_{29}N_3O_3{}^{127}I$ [M+H] 558.1254; Found 558.1260.

SMILES:
O=C([C@H]1CN(C(OC(C)(C)C)=O)CC[C@H]1C2=CC=C(I)C=C2)NC3=C(N=CC=C4)C4=CC=C3

InChI=1S/C26H28IN3O3/c1-26(2,3)33-25(32)30-15-13-20(17-9-11-19(27)12-10-17)21(16-30)24(31)29-22-8-4-6-18-7-5-14-28-23(18)22/h4-12,14,20-21H,13,15-16H2,1-3H3,(H,29,31)/t20-,21-/m0/s1

Minor (**(±)-S3b**):
$R_f$ 0.41 (5% $CH_3CN/CH_2Cl_2$);
mp = 93–96 °C (from $Et_2O$/pentane); $\nu_{max}$ (film)/cm$^{-1}$ 3336 (NH), 2922, 1677 (C=O), 1521, 1483, 1424, 1323, 1230, 1156, 1062, 1003, 824, 757;
$^1$H NMR (500 MHz, $(CD_3)_2SO$, 373 K) δ 9.73 (br s, 1 H, NH), 8.85 (dd, *J* = 4.2, 1.7 Hz, 1 H, $HC_{Ar}$), 8.39 (dd, *J* = 7.7, 1.3 Hz, 1 H, $HC_{Ar}$), 8.31 (dd, *J* = 8.3, 1.7 Hz, 1 H, $HC_{Ar}$), 7.62–7.55 (m, 2 H, $HC_{Ar}$), 7.55–7.51 (m, 2 H, $HC_{Ar}$), 7.47 (t, *J* = 8.0 Hz, 1 H, $HC_{Ar}$), 7.19–7.14 (m, 2 H, $HC_{Ar}$), 4.35 (ddd, *J* = 12.8, 3.8, 1.8 Hz, 1 H, NC*H*HCHCO), 4.12 (ddt, *J* = 13.3, 4.4, 2.1 Hz, 1 H, NC*H*HCH$_2$), 3.17–2.99 (m, 3 H, NCH*H*CHCO, CHCO, CHAr), 2.98–2.90 (m, 1 H, NCH*H*CH$_2$), 1.80 (dq, *J* = 13.3, 3.0 Hz, 1 H, NCH$_2$C*H*H), 1.65 (qd, *J* = 12.7, 4.6 Hz, 1 H, NCH$_2$CH*H*), 1.48 (s, 9 H, $C(CH_3)_3$);
$^{13}$C NMR (126 MHz, $(CD_3)_2SO$, 373 K) δ 169.8 (C=O amide), 153.4 (C=O carbamate), 148.1 ($C_{Ar}$), 142.8 ($C_{Ar}$ quat), 137.7 ($C_{Ar}$ quat), 136.6 (2 × $C_{Ar}$), 135.7 ($C_{Ar}$), 133.5 ($C_{Ar}$ quat), 129.3 (2 × $C_{Ar}$), 127.2 ($C_{Ar}$ quat), 126.1 ($C_{Ar}$), 121.4 ($C_{Ar}$), 121.3 ($C_{Ar}$), 116.3 ($C_{Ar}$), 91.1 ($IC_{Ar}$ quat), 78.6 (*C*(CH$_3$)$_3$), 49.1 (*C*HCO), 46.2 (N*C*H$_2$CHCO), 44.0 (*C*HAr), 43.3 (N*C*H$_2$CH$_2$), 32.0 (NCH$_2$*C*H$_2$), 27.7 (C(*C*H$_3$)$_3$);
HRMS (ESI$^+$) *m/z* Calculated for $C_{26}H_{29}N_3O_3{}^{127}I$ [M+H] 558.1254; Found 558.1247.

SMILES:
O=C([C@@H]1CN(C(OC(C)(C)C)=O)CC[C@H]1C2=CC=C(I)C=C2)NC3=C(N=CC=C4)C4=CC=C3

InChI=1S/C26H28IN3O3/c1-26(2,3)33-25(32)30-15-13-20(17-9-11-19(27)12-10-17)21(16-30)24(31)29-22-8-4-6-18-7-5-14-28-23(18)22/h4-12,14,20-21H,13,15-16H2,1-3H3,(H,29,31)/t20-,21+/m0/s1

***tert*-Butyl (+)-(*3R,4R*)-4-(4-iodophenyl)-3-(quinolin-8-ylcarbamoyl)piperidine-1-carboxylate ((+)-6b)** and ***tert*-butyl (–)-(*3R,4S*)-4-(4-iodophenyl)-3-(quinolin-8-ylcarbamoyl)piperidine-1-carboxylate ((–)-S3b)**:

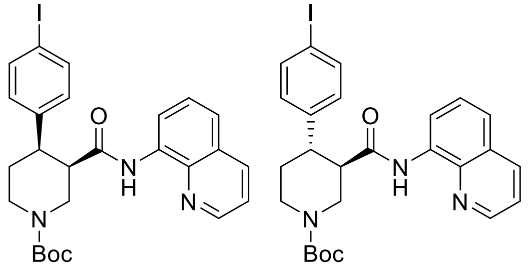

A large microwave vial (10–20 mL recommended volume) was charged with $K_2CO_3$ (553 mg, 4.0 mmol, 1 equiv), flame-dried, and allowed to cool under argon. *tert*-Butyl (*R*)-3-(quinolin-8-ylcarbamoyl)piperidine-1-carboxylate **(–)-5** (1.42 g, 4.0 mmol 1 equiv), 1,4-diiodobenzene (5.28 g, 16.0 mmol, 4 equiv), $Pd(OAc)_2$ (45.1 mg, 0.2 mmol, 5 mol %) and PivOH (409 mg, 4.0 mmol, 1 equiv) were added sequentially. The reaction vessel was sealed with an aluminum cap (with molded butyl/PTFE septa) and purged with argon, then anhydrous $PhCF_3$ (2.0 mL, 2.00 M) was added by syringe. The reaction tube was then placed in a preheated oil bath and stirred at 110 ºC for 18 h. The reaction mixture was then allowed to cool to rt and EtOAc (20 mL) was added. The resulting mixture was filtered through a pad of Celite®, eluting with further EtOAc (2 × 50 mL). The solvent was removed under reduced pressure. The reaction mixture was purified by two consecutive chromatographic separations: one (0% to 5% $CH_3CN/CH_2Cl_2$) to isolate the minor *trans*-product *tert*-butyl (–)-(*3R,4S*)-4-(4-iodophenyl)-3-(quinolin-8-ylcarbamoyl)piperidine-1-carboxylate **(–)-S3b** followed by a second (10% to 15% acetone/pentane) to isolate the major *cis*-product *tert*-butyl (+)-(*3R,4R*)-4-(4-iodophenyl)-3-(quinolin-8-ylcarbamoyl)piperidine-1-carboxylate **(+)-6b**. The product containing fractions were combined and the solvent was removed under reduced pressure. $Et_2O$ (20 mL) and pentane (20 mL) were added and the solvent was removed under reduced pressure to afford the minor *trans*-product **(–)-S3b** as a pale orange solid (441 mg, 20%, 98.1% *ee*) and the major *cis*-product **(+)-6b** (775 mg, 35%, 98.2% *ee*).

Major (**(+)-6b**):
$[\alpha]_D^{23}$ + 9.1 (*c* 1.1, $CHCl_3$).
Characterization data identical to that reported for racemic *cis*-piperidine **(±)-S2b** (see S24).

**HPLC Conditions**: Chiralpak IA 3-column, 85:15 *n*-hexane:*i*-PrOH, flow rate: 1 mL·min$^{-1}$, 35 ºC, UV detection wavelength: 210.4 nm. Retention times: 12.2 min (3*S*,4*S* enantiomer), 17.7 min (3*R*,4*R* enantiomer).

SMILES:
O=C([C@H]1CN(C(OC(C)(C)C)=O)CC[C@H]1C2=CC=C(I)C=C2)NC3=C(N=CC=C4)C4=CC=C3

InChI=1S/C26H28IN3O3/c1-26(2,3)33-25(32)30-15-13-20(17-9-11-19(27)12-10-17)21(16-30)24(31)29-22-8-4-6-18-7-5-14-28-23(18)22/h4-12,14,20-21H,13,15-16H2,1-3H3,(H,29,31)/t20-,21-/m0/s1

Minor (**(–)-S3b**):
$[\alpha]_D^{23}$ – 45.5 (*c* 1.1, $CHCl_3$).
Characterization data identical to that reported for racemic *trans*-piperidine **(±)-S3b** (see S24).

**HPLC Conditions**: Chiralpak IA 3-column, 85:15 *n*-hexane:*i*-PrOH, flow rate: 1 mL·min$^{-1}$, 35 ºC, UV detection wavelength: 254.1 nm. Retention times: 9.4 min (3*R*,4*S* enantiomer), 13.3 min (3*S*,4*R* enantiomer).

SMILES:
O=C([C@H]1CN(C(OC(C)(C)C)=O)CC[C@@H]1C2=CC=C(I)C=C2)NC3=C(N=CC=C4)C4=CC=C3

InChI=1S/C26H28IN3O3/c1-26(2,3)33-25(32)30-15-13-20(17-9-11-19(27)12-10-17)21(16-30)24(31)29-22-8-4-6-18-7-5-14-28-23(18)22/h4-12,14,20-21H,13,15-16H2,1-3H3,(H,29,31)/t20-,21+/m1/s1

***tert*-Butyl (+)-(*3S,4R*)-4-(4-iodophenyl)-3-(quinolin-8-ylcarbamoyl)piperidine-1-carboxylate ((+)-7b)**

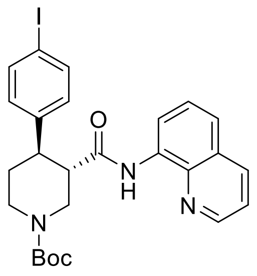


A flame-dried reaction tube was charged with *cis*-3,4-disubstituted piperidine **(+)-6b** (687 mg, 1.23 mmol, 1 equiv) and 1,8-diazabicyclo(5.4.0)undec-7-ene (DBU, 550 μL, 3.70 mmol, 3 equiv). The reaction vessel was sealed with an aluminum cap (with molded butyl/PTFE septa) and purged with argon, then anhydrous toluene (1.20 mL, 1.0 M) was added by syringe. The reaction tube was then placed in a preheated oil bath and stirred at 110 ºC for 24 h. The reaction mixture was then allowed to cool to rt and $CH_2Cl_2$ (5 mL) and sat. aq. $NH_4Cl$ (5 mL) were added. The phases were separated, and the aqueous layer was extracted with $CH_2Cl_2$ (3 × 10 mL). The combined organic extracts were dried over $Na_2SO_4$ and filtered. The solvent was removed under reduced pressure. The reaction mixture was purified by flash column chromatography (10% acetone/pentane). The product containing fractions were combined and the solvent was removed under reduced pressure. $Et_2O$ (10 mL) and pentane (10 mL) were added and the solvent was removed under reduced pressure to afford amide *tert*-butyl (+)-(*3S,4R*)-4-(4-iodophenyl)-3-(quinolin-8-ylcarbamoyl) piperidine-1-carboxylate **(+)-7b** as a white solid (626 mg, 91%, 98.0% *ee*).

$[\alpha]_D^{23}$ + 48.0 (*c* 1.0, $CHCl_3$).
Characterization data identical to that reported for racemic *trans*-piperidine **(±)-S3b** (see S24).

**HPLC Conditions**: Chiralpak IA 3-column, 85:15 *n*-hexane:*i*-PrOH, flow rate: 1 mL·min$^{-1}$, 35 ºC, UV detection wavelength: 254.1 nm. Retention times: 9.4 min (3*R*,4*S* enantiomer), 13.3 min (3*S*,4*R* enantiomer).

SMILES:
O=C([C@@H]1CN(C(OC(C)(C)C)=O)CC[C@H]1C2=CC=C(I)C=C2)NC3=C(N=CC=C4)C4=CC=C3

InChI=1S/C26H28IN3O3/c1-26(2,3)33-25(32)30-15-13-20(17-9-11-19(27)12-10-17)21(16-30)24(31)29-22-8-4-6-18-7-5-14-28-23(18)22/h4-12,14,20-21H,13,15-16H2,1-3H3,(H,29,31)/t20-,21+/m0/s1

**_tert_-Butyl (+)-(_3R,4S_)-4-(4-iodophenyl)-3-(hydroxymethyl)piperidine-1-carboxylate ((+)-S4b)**

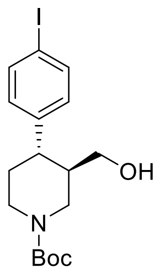


A flame-dried reaction tube was charged with amide **(–)-S3b** (111 mg, 0.20 mmol, 1 equiv), followed by di-*tert*-butyl dicarbonate ($Boc_2O$, 175 mg, 0.80 mmol, 4 equiv) and 4-(dimethylamino)pyridine (DMAP, 4.9 mg, 0.04 mmol, 20 mol %). The reaction vessel was sealed with an aluminum cap (with molded butyl septa) and purged with argon, then anhydrous MeCN (400 μL, 0.5 M) was added by syringe. The mixture was then stirred at 40 ºC for 22 h. The reaction mixture was then allowed to cool to rt and sat. aq. $NH_4Cl$ (1 mL) and $CH_2Cl_2$ (1 mL) were added. The phases were separated, and the aqueous layer was extracted with $CH_2Cl_2$ (3 × 5 mL). The combined organic extracts were dried over $Na_2SO_4$ and filtered. The solvent was removed under reduced pressure to afford the crude *N*-Boc protected piperidine derivative.

This crude was solubilized in anhydrous THF (800 μL, 0.2 M) and the resulting solution was added dropwise to a suspension of $LiAlH_4$ (15.2 mg, 0.40 mmol, 2 equiv) in anhydrous THF (200 μL, 2.0 M) at 0 °C under argon atmosphere. The mixture was then stirred at 20 °C for 30 min. The reaction mixture was then quenched by slow addition of sat. aq. $NH_4Cl$ (2 mL) at 0 °C and stirred at rt for 30 min. The resulting suspension was filtered through a pad of Celite®, eluting with EtOAc (3 × 5 mL). The phases were separated, and the aqueous layer was extracted with EtOAc (3 × 5 mL). The combined organic extracts were dried over $Na_2SO_4$ and filtered. The solvent was removed under reduced pressure. Purification by flash column chromatography (10% to 15% acetone/pentane) afforded primary alcohol **(+)-S4b** as a white solid (52.3 mg, 63% over 2 steps, 98.1% *ee*, containing approx. 10% deiodinated derivative).

$[\alpha]_D^{23}$ + 2.0 (*c* 1.0, $CHCl_3$).
$R_f$ 0.24 (15% acetone/pentane);
mp = 53–59 °C;
$\nu_{max}$ (film)/cm$^{-1}$ 3422 (OH), 2922, 1662 (C=O), 1479, 1424, 1364, 1234, 1163, 1129, 1059, 1006, 816, 764; $^1$H NMR (400 MHz, $CDCl_3$, 298 K) δ 7.66–7.61 (m, 2 H, $HC_{Ar}$), 6.99–6.93 (m, 2 H, $HC_{Ar}$), 4.36 (br d, *J* = 13.2 Hz, 1 H, NC*H*HCHCH$_2$OH), 4.20 (br s, 1 H, NC*H*HCH$_2$), 3.44 (dt, *J* = 11.0, 3.5 Hz, 1 H, C*H*HOH), 3.26 (dt, *J* = 11.3, 5.8 Hz, 1 H, CH*H*OH), 2.87–2.63 (m, 2 H, NCH*H*CHCH$_2$OH, NCH*H*CH$_2$), 2.51 (dt, *J* = 10.2, 5.2 Hz, 1 H, CHAr), 1.87–1.72 (m, 2 H, C*H*CH$_2$OH, NCH$_2$C*H*H), 1.71–1.58 (m, 2 H, NCH$_2$CH*H*, OH), 1.49 (s, 9 H, C(CH$_3$)$_3$);
$^{13}$C NMR (101 MHz, $CDCl_3$, 298 K, observed as a mixture of rotamers) δ 154.8 (C=O), 143.5 ($C_{Ar}$ quat), 137.7 (2 × $C_{Ar}$), 129.5 (2 × $C_{Ar}$), 91.7 (I$C_{Ar}$ quat), 79.7 (*C*(CH$_3$)$_3$), 63.0 (*C*H$_2$OH), 46.4 (br m, N*C*H$_2$CHCH$_2$OH), 44.4 and 43.5 (N*C*H$_2$CH$_2$, *C*HAr, *C*HCH$_2$OH), 33.8 (NCH$_2$*C*H$_2$), 28.5 (C(*C*H$_3$)$_3$);
HRMS (ESI$^+$) *m/z* Calculated for $C_{17}H_{25}NO_3{}^{127}I$ [M+H] 418.0879; Found 418.0886.

**HPLC Conditions**: Chiralpak ID 3-column, 90:10 *n*-hexane:*i*-PrOH, flow rate: 1 mL·min$^{-1}$, 35 ºC, UV detection wavelength: 230.1 nm. Retention times: 6.7 min (3*R*,4*S* enantiomer), 7.4 min (3*S*,4*R* enantiomer).

SMILES: IC1=CC=C([C@@H]2[C@@H](CO)CN(C(OC(C)(C)C)=O)CC2)C=C1

InChI=1S/C17H24INO3/c1-17(2,3)22-16(21)19-9-8-15(13(10-19)11-20)12-4-6-14(18)7-5-12/h4-7,13,15,20H,8-11H2,1-3H3/t13-,15-/m1/s1

***tert*-Butyl (–)-(*3S,4R*)-4-(4-iodophenyl)-3-(hydroxymethyl)piperidine-1-carboxylate ((–)-8b)**

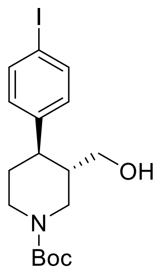

A flame-dried round-bottom flask was charged with amide **(+)-7b** (558 mg, 1.00 mmol, 1 equiv), followed by di-*tert*-butyl dicarbonate ($Boc_2O$, 873 mg, 4.00 mmol, 4 equiv) and 4-(dimethylamino)pyridine (DMAP, 24.4 mg, 0.20 mmol, 20 mol %). The reaction vessel was sealed with an aluminum cap (with molded butyl septa) and purged with argon, then anhydrous MeCN (3.3 mL) and anhydrous $CH_2Cl_2$ (0.5 mL) were added by syringe. The mixture (0.3 M) was then stirred at 40 ºC for 22 h. The reaction mixture was then allowed to cool to rt and sat. aq. $NH_4Cl$ (5 mL) and $CH_2Cl_2$ (5 mL) were added. The phases were separated, and the aqueous layer was extracted with $CH_2Cl_2$ (3 × 10 mL). The combined organic extracts were dried over $Na_2SO_4$ and filtered. The solvent was removed under reduced pressure to afford the crude *N*-Boc protected piperidine derivative.
This crude solubilized in anhydrous THF (3.5 mL, 0.3 M) and the resulting solution was added dropwise to a suspension of $LiAlH_4$ (75.9 mg, 2.00 mmol, 2 equiv) in anhydrous THF (1.5 mL, 1.0 M) at 0 °C under argon atmosphere. The mixture was then stirred at 20 °C for 30 min. The reaction mixture was then quenched by slow addition of sat. aq. $NH_4Cl$ (5 mL) at 0 °C and stirred at rt for 30 min. The resulting suspension was filtered through a pad of Celite®, eluting with EtOAc (3 × 10 mL). The phases were separated, and the aqueous layer was extracted with EtOAc (3 × 10 mL). The combined organic extracts were dried over $Na_2SO_4$ and filtered. The solvent was removed under reduced pressure. Purification by flash column chromatography (10% to 15% acetone/pentane) afforded primary alcohol **(–)-8b** as a white solid (315 mg, 68% over 2 steps, 98.0% *ee*, containing approx. 15% deiodinated derivative).

$[\alpha]_D^{23}$ – 8.0 (*c* 1.0, $CHCl_3$).
Characterization data identical to that reported for enantiomeric alcohol **(+)-S4b** (see S27).

**HPLC Conditions**: Chiralpak ID 3-column, 90:10 *n*-hexane:*i*-PrOH, flow rate: 1 mL·min$^{-1}$, 35 ºC, UV detection wavelength: 230.1 nm. Retention times: 6.7 min (3*R,*4*S* enantiomer), 7.4 min (3*S,*4*R* enantiomer).

SMILES: IC1=CC=C([C@H]2[C@H](CO)CN(C(OC(C)(C)C)=O)CC2)C=C1

InChI=1S/C17H24INO3/c1-17(2,3)22-16(21)19-9-8-15(13(10-19)11-20)12-4-6-14(18)7-5-12/h4-7,13,15,20H,8-11H2,1-3H3/t13-,15-/m0/s1

***tert*-Butyl (3*S*,4*R*)-3-((benzo[d][1,3]dioxol-5-yloxy)methyl)-4-(4-iodophenyl)piperidine-1-carboxylate ((–)-9b)**

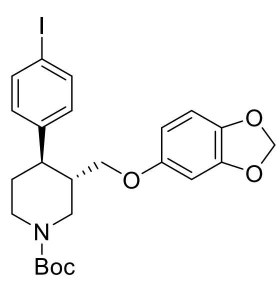

Alcohol **(–)-8b** (203 mg, 0.49 mmol, 1 equiv) and triethylamine (96 μL, 0.69 mmol, 1.4 equiv) were added to a flame-dried round-bottom flask, dissolved in anhydrous $CH_2Cl_2$ (2.5 mL, 0.2 M) and cooled down to 0 °C. Methanesulfonyl chloride (49 μL, 0.64 mmol, 1.3 equiv) was then added by Gilson pipette. After stirring 5 min at 0 °C, the reaction mixture was stirred at 25 °C for 2 h, then diluted with $CH_2Cl_2$ (5 mL) and sat. aq. $NaHCO_3$ (5 mL). The phases were separated, and the aqueous layer was extracted with $CH_2Cl_2$ (3 × 10 mL). The combined organic extracts were dried over $Na_2SO_4$ and filtered. The solvent was removed under reduced pressure to afford the crude mesylated alcohol derivative.
NaH (60% dispersion in mineral oil, 45.2 mg, 1.10 mmol, 2.2 equiv) was added to a solution of sesamol (135 mg, 0.98 mmol, 2 equiv) in anhydrous DMF (3.0 mL, 0.3 M) at 0 °C. The mixture was then stirred at 25 °C for 1 h. A solution of the crude mesylated alcohol in dry DMF (2.0 mL, 0.2 M) was then added dropwise to this suspension. The resulting mixture was stirred at 90 °C for 20 h. The reaction mixture was quenched by addition of $H_2O$ (5 mL) and aq NaOH 1 N (5 mL) and EtOAc (10 mL) were then added.

The phases were separated, and the aqueous layer was extracted with EtOAc (4 × 20 mL). The combined organic extracts were washed with brine (2 × 50 mL), dried over $Na_2SO_4$ and filtered. The solvent was removed under reduced pressure. Purification by flash column chromatography (5% acetone/pentane) afforded piperidine **(–)-9b** as a white solid (188 mg, 71% over 2 steps).

$[\alpha]_D^{23}$ – 43.3 (*c* 1.2, $CHCl_3$).
$R_f$ 0.15 (5% acetone/pentane);
mp = 51–54 °C; $\nu_{max}$ (film)/cm$^{-1}$ 2919, 1685 (C=O), 1483, 1424, 1230, 1163, 1129, 1036, 1106, 928, 813, 764; $^1$H NMR (400 MHz, $CDCl_3$, 298 K) δ 7.65–7.59 (m, 2 H, $HC_{Ar}$), 6.67–6.91 (m, 2 H, $HC_{Ar}$), 6.64 (d, *J* = 8.4 Hz, 1 H, $HC_{Ar}$), 6.36 (d, *J* = 2.5 Hz, 1 H, $HC_{Ar}$), 6.14 (dd, *J* = 8.5, 2.5 Hz, 1 H, $HC_{Ar}$), 5.89 (s, 2 H, $OCH_2O$), 4.43 (br s, 1 H, NC*H*HCHCH$_2$OAr), 4.25 (br s, 1 H, NC*H*HCH$_2$), 3.61 (dd, *J* = 9.4, 2.9 Hz, 1 H, C*H*HOAr), 3.45 (dd, *J* = 9.4, 6.4 Hz, 1 H, CH*H*OAr), 2.91–2.71 (br m, 2 H, NCH*H*CHCH$_2$OAr, NCH*H*CH$_2$), 2.65 (td, *J* = 11.8, 3.8 Hz, 1 H, CHAr), 2.08–1.96 (br m, 1 H, C*H*CH$_2$OAr), 1.86–1.76 (br m, 1 H, NCH$_2$C*H*H), 1.76–1.63 (m, 1 H, NCH$_2$CH*H*), 1.50 (s, 9 H, $C(CH_3)_3$);
$^{13}$C NMR (101 MHz, $CDCl_3$, 298 K) δ 154.7 (C=O), 154.2 ($OC_{Ar}$ quat), 148.1 ($OC_{Ar}$ quat), 143.1 ($C_{Ar}$ quat), 141.7 ($OC_{Ar}$ quat), 137.7 (2 × $C_{Ar}$), 129.4 (2 × $C_{Ar}$), 107.8 ($C_{Ar}$), 105.5 ($C_{Ar}$), 101.1 ($OCH_2O$), 98.0 ($C_{Ar}$), 91.8 ($IC_{Ar}$ quat), 79.7 (*C*(CH$_3$)$_3$), 68.7 (*C*H$_2$OAr), 47.0 (br m, N*C*H$_2$CHCH$_2$OAr), 44.3 (N*C*H$_2$CH$_2$, *C*HAr), 41.6 (*C*HCH$_2$OAr), 33.6 (NCH$_2$*C*H$_2$), 28.4 (C(*C*H$_3$)$_3$);
HRMS (ESI$^+$) *m/z* Calculated for $C_{24}H_{29}NO_5{}^{127}I$ [M+H] 538.1090; Found 538.1104.

SMILES: IC1=CC=C([C@H]2[C@H](COC3=CC(OCO4)=C4C=C3)CN(C(OC(C)(C)C)=O)CC2)C=C1

InChI=1S/C24H28INO5/c1-24(2,3)31-23(27)26-11-10-20(16-4-6-18(25)7-5-16)17(13-26)14-28-19-8-9-21-22(12-19)30-15-29-21/h4-9,12,17,20H,10-11,13-15H2,1-3H3/t17-,20-/m0/s1

**(3*S*,4*R*)-3-((Benzo[d][1,3]dioxol-5-yloxy)methyl)-4-(4-iodophenyl)piperidine-1-ium chloride (3** · HCl)

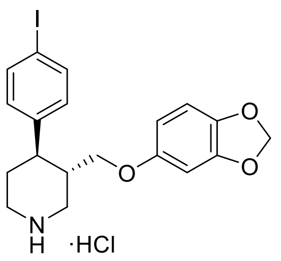


4 N HCl in 1,4-dioxane (250 μL, 1.00 mmol, 10 equiv) was added to a solution of *N*-Boc protected piperidine **(–)-9b** (56.9 mg, 0.10 mmol) in 1,4-dioxane (250 μL, 0.4 M). at 0 °C under air. The solution was stirred at 25 ºC for 18 h, then an ice-cold 1:1 mixture of $Et_2O$/pentane (1 mL) was added and formation of a solid precipitate was observed. This was filtered and washed with further ice-cold $Et_2O$/pentane mixture (2 × 5 mL). The solid precipitate was dried under reduced pressure to afford (3*S*,4*R*)-3-((benzo[d][1,3]dioxol-5-yloxy)methyl)-4-(4-iodophenyl)piperidine-1-ium chloride **3** · HCl (38.5 mg, 81%) as an off-white solid.

$[\alpha]_D^{23}$ – 86.0 (*c* 0.9, MeOH).
mp = 203–205 ºC;
$\nu_{max}$ (film)/cm$^{-1}$ 3321 (NH), 2926, 2807, 1618, 1484, 1185, 1103, 1033, 1003, 932, 846, 813, 787;
$^1$H NMR (400 MHz, $CD_3OD$, 298 K) δ 7.71–7.64 (m, 2 H, $HC_{Ar}$), 7.11–7.04 (m, 2 H, $HC_{Ar}$), 6.63 (d, *J* = 8.5 Hz, 1 H, $HC_{Ar}$), 6.39 (d, *J* = 2.5 Hz, 1 H, $HC_{Ar}$), 6.18 (dd, *J* = 8.5, 2.5 Hz, 1 H, $HC_{Ar}$), 5.89–5.82 (m, 2 H, $OCH_2O$), 3.71–3.62 (m, 2 H, C*H*HOAr, NC*H*HCHCH$_2$OAr), 3.60–3.48 (m, 2 H, CH*H*OAr, NC*H*HCH$_2$), 2.21–2.11 (m, 2 H, NCH*H*CHCH$_2$OAr, NCH*H*CH$_2$), 3.00–2.90 (m, 1 H, CHAr), 2.49–2.37 (m, 1 H, C*H*CH$_2$OAr), 2.09–2.00 (m, 2 H, NCH$_2$C*H*$_2$);
$^{13}$C NMR (101 MHz, $CD_3OD$, 298 K) δ 155.2 ($OC_{Ar}$ quat), 149.7 ($OC_{Ar}$ quat), 143.5 ($C_{Ar}$ quat), 143.0 ($OC_{Ar}$ quat), 139.1 (2 × $C_{Ar}$), 130.7 (2 × $C_{Ar}$), 108.8 ($C_{Ar}$), 106.6 ($C_{Ar}$), 102.5 ($OCH_2O$), 98.9 ($C_{Ar}$), 93.1 ($IC_{Ar}$ quat), 68.9 (*C*H$_2$OAr), 47.7 (N*C*H$_2$CHCH$_2$OAr), 45.4 (N*C*H$_2$CH$_2$), 43.0 (*C*HAr), 40.5 (*C*HCH$_2$OAr), 31.3 (NCH$_2$*C*H$_2$);
HRMS (ESI$^+$) *m/z* Calculated for $C_{19}H_{21}NO_3{}^{127}I$ [M–Cl] 438.0566; Found 438.0571.

SMILES: IC1=CC=C([C@H]2[C@H](COC3=CC(OCO4)=C4C=C3)CNCC2)C=C1.Cl

InChI=1S/C19H20INO3.ClH/c20-15-3-1-13(2-4-15)17-7-8-21-10-14(17)11-22-16-5-6-18-19(9-16)24-12-23-18;/h1-6,9,14,17,21H,7-8,10-12H2;1H/t14-,17-;/m0./s1

### NMR Spectra for Novel Compounds

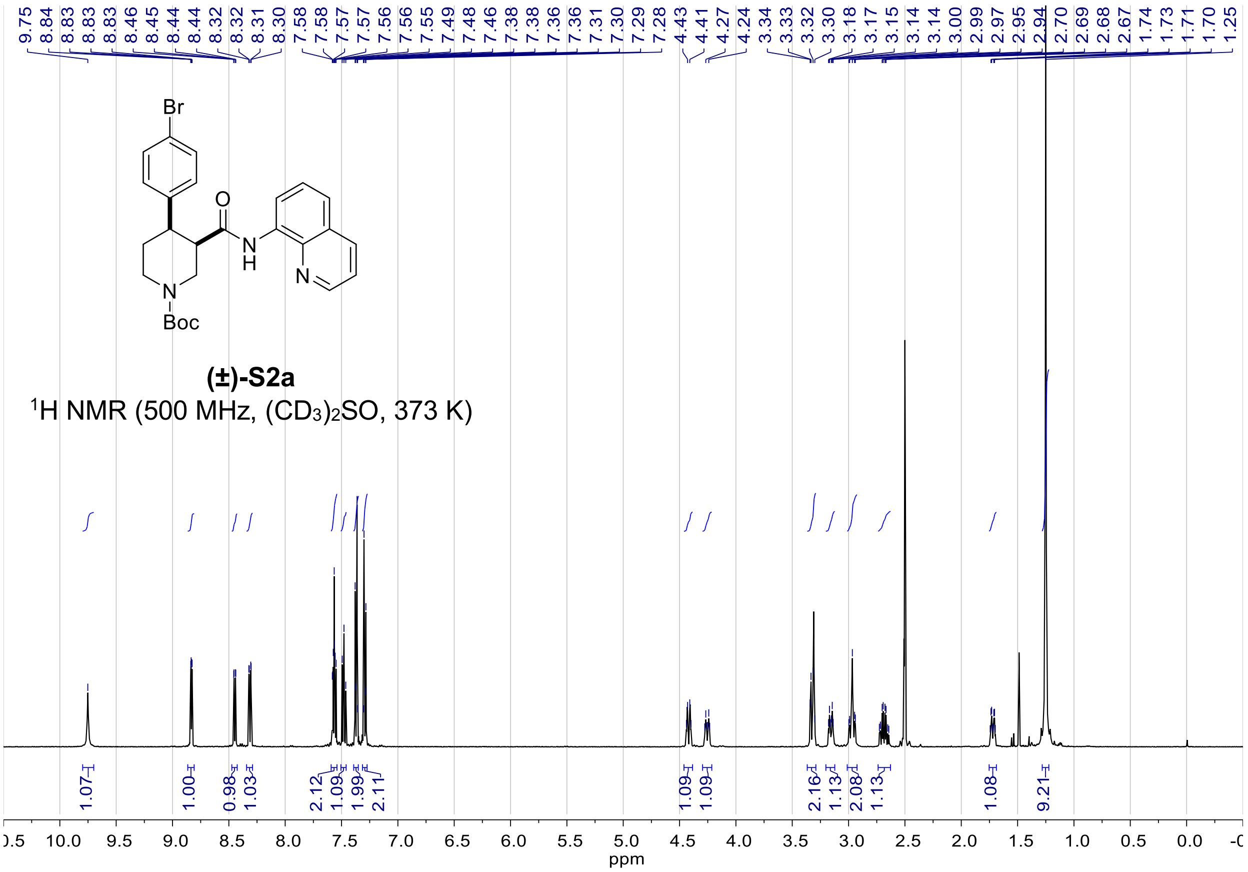


**(±)-S2a**

$^{1}$H NMR (500 MHz, $(CD_3)_2SO$, 373 K)

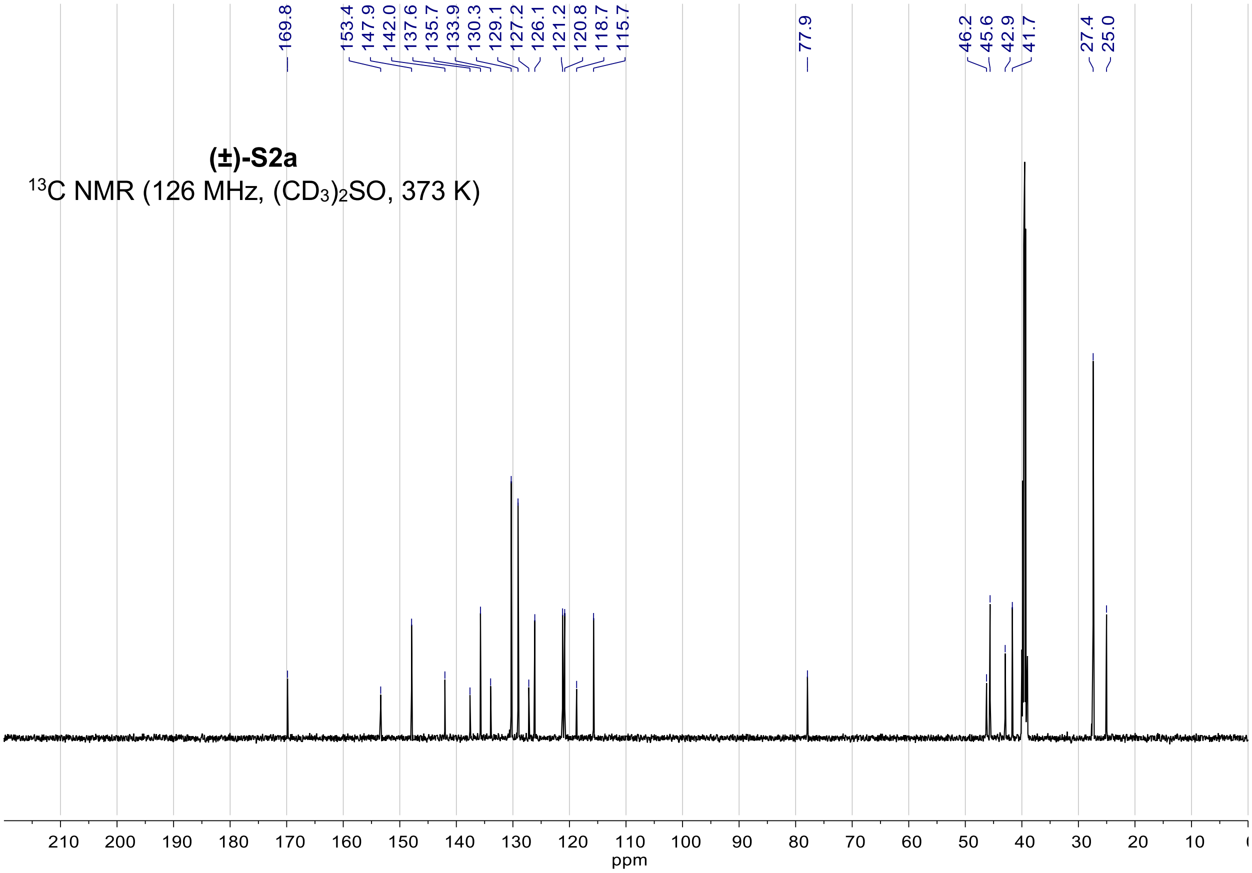


**(±)-S2a**

$^{13}$C NMR (126 MHz, $(CD_3)_2SO$, 373 K)

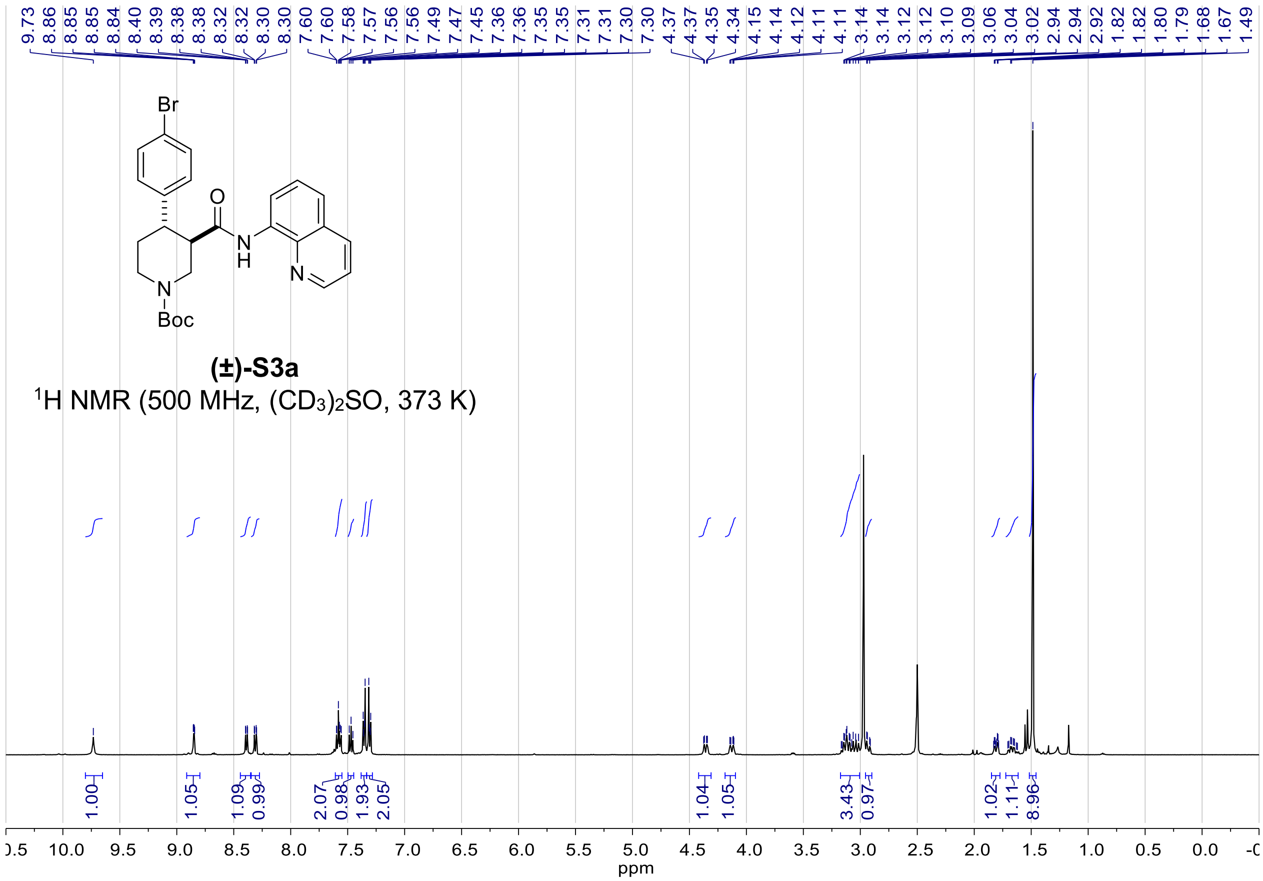

**(±)-S3a**

$^1$H NMR (500 MHz, $(CD_3)_2SO$, 373 K)

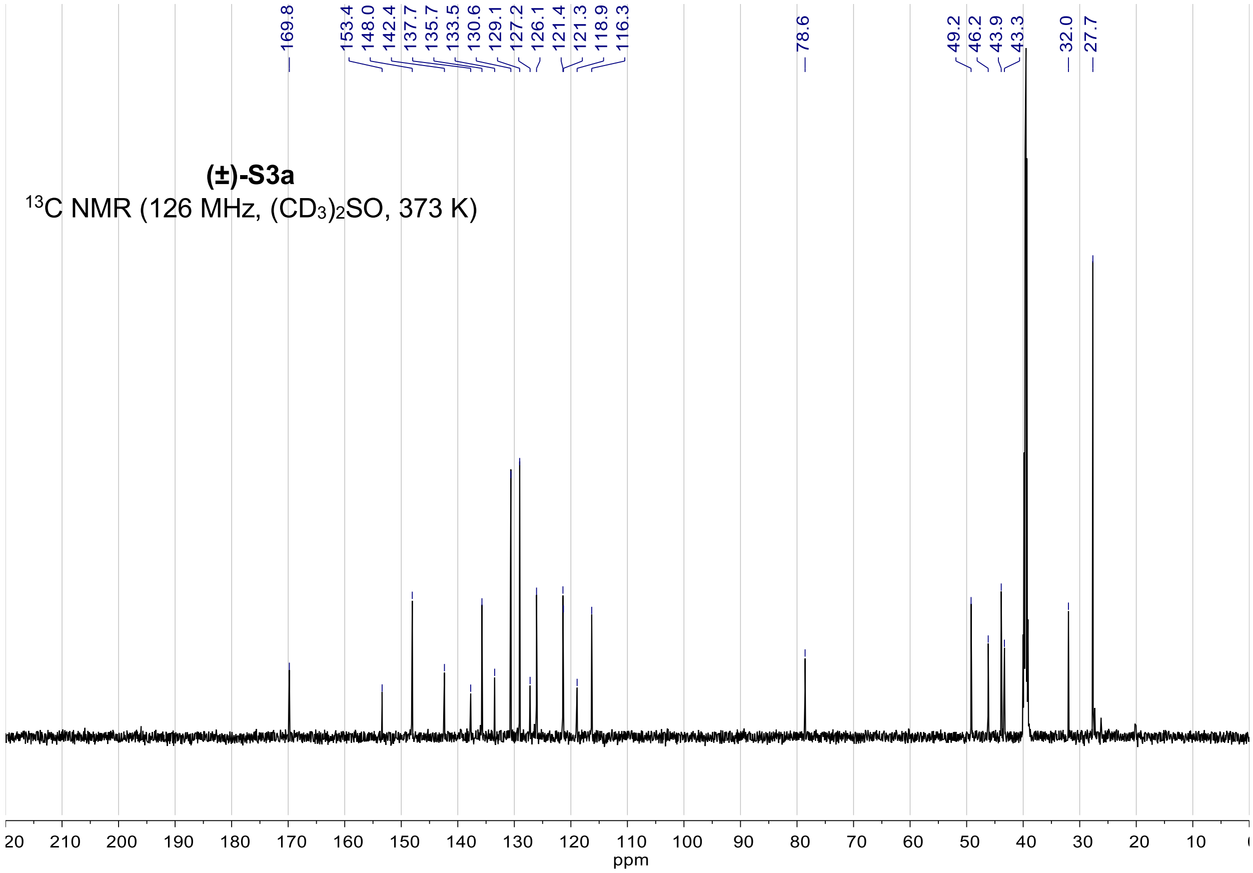

**(±)-S3a**

$^{13}$C NMR (126 MHz, $(CD_3)_2SO$, 373 K)

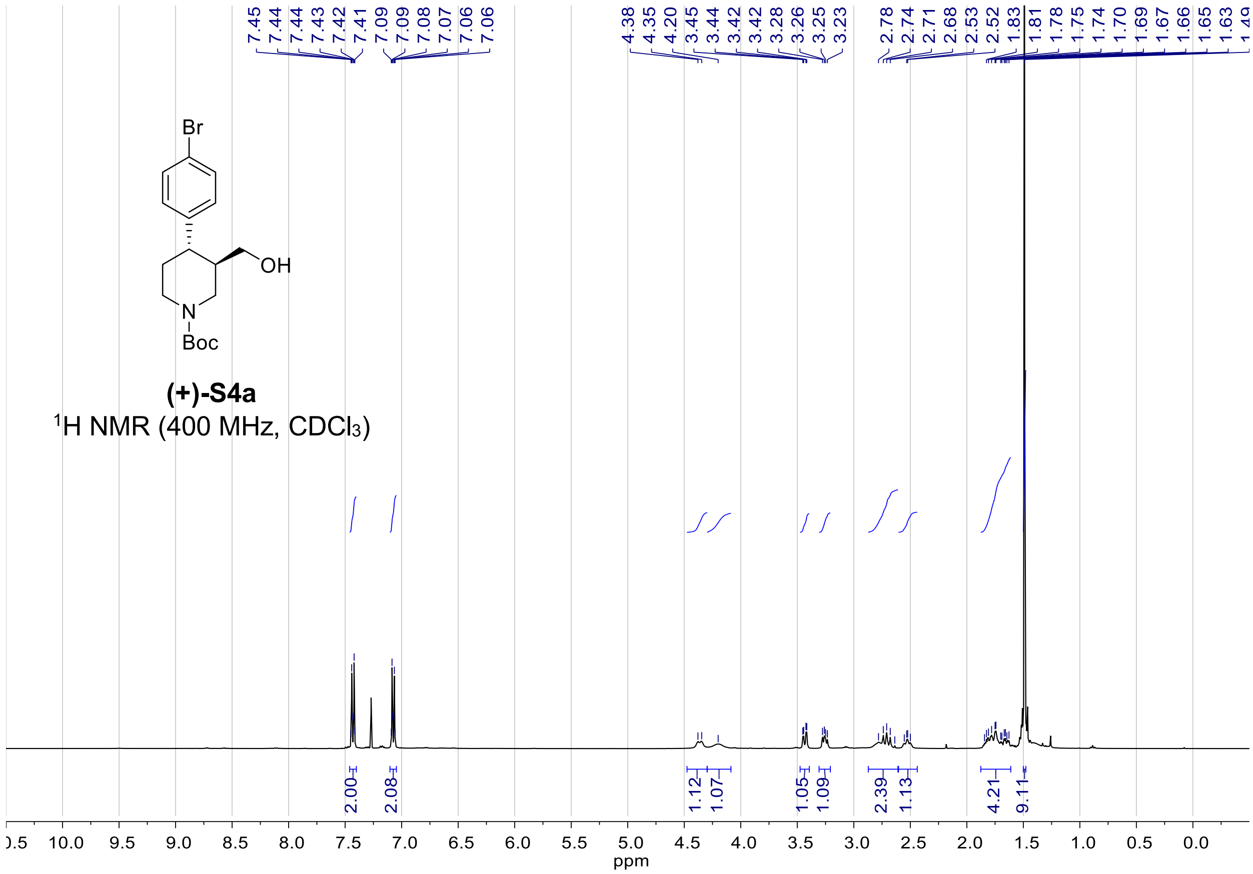
7.45
7.44
7.44
7.43
7.42
7.41
7.09
7.09
7.08
7.07
7.06
7.06
4.38
4.35
4.20
3.45
3.44
3.42
3.42
3.28
3.26
3.25
3.23
2.78
2.74
2.71
2.68
2.53
2.52
1.83
1.81
1.78
1.75
1.74
1.70
1.69
1.67
1.66
1.65
1.63
Br
OH
N
Boc
(+)-S4a
1H NMR (400 MHz, CDCl3)
2.00
2.08
1.12
1.07
1.05
1.09
2.39
1.13
4.21
9.11
10.0
9.5
9.0
8.5
8.0
7.5
7.0
6.5
6.0
5.5
5.0
4.5
4.0
3.5
3.0
2.5
2.0
1.5
1.0
0.5
0.0
ppm

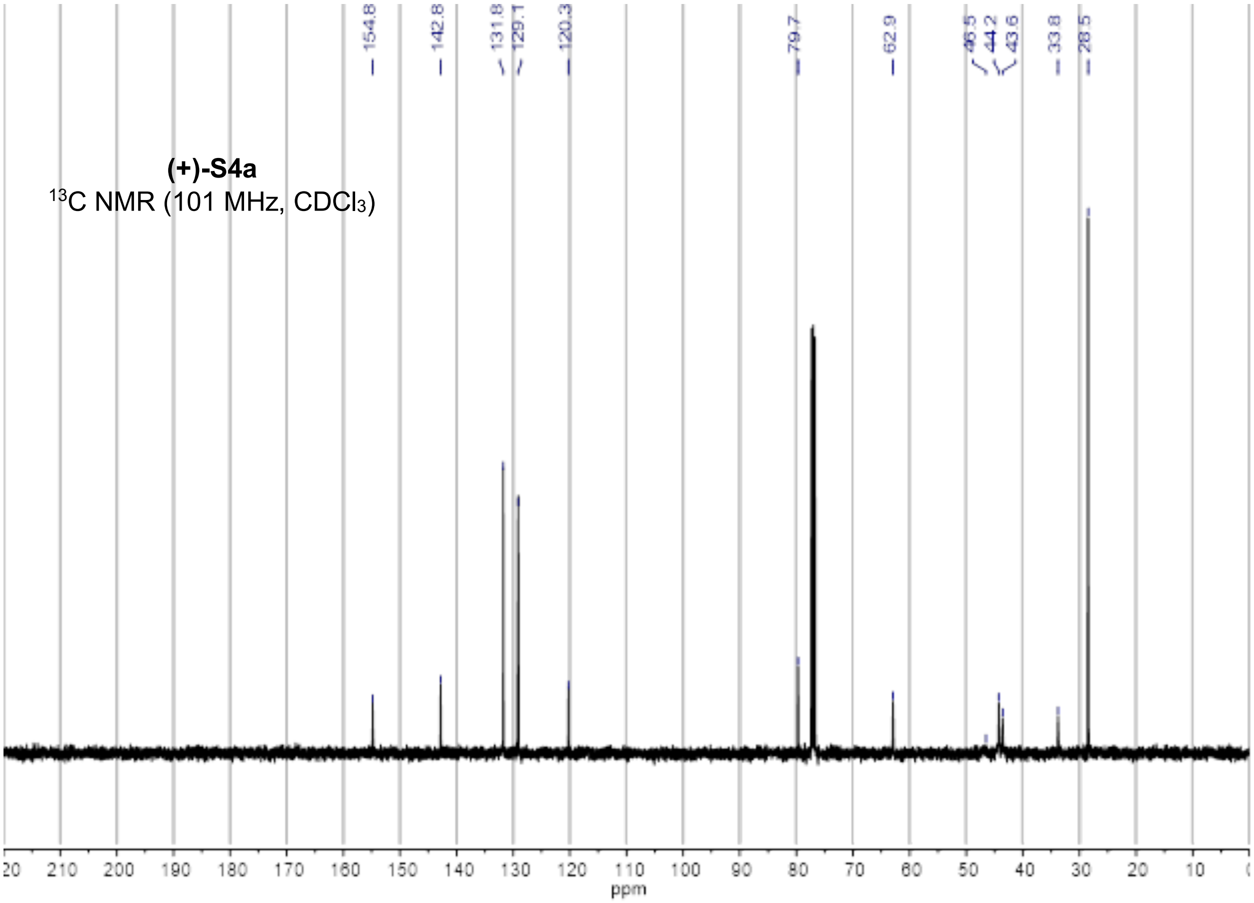
154.8
142.8
131.8
129.1
120.3
79.7
62.9
46.5
44.2
43.6
33.8
28.5
(+)-S4a
13C NMR (101 MHz, CDCl3)
210
200
190
180
170
160
150
140
130
120
110
100
90
80
70
60
50
40
30
20
10
ppm

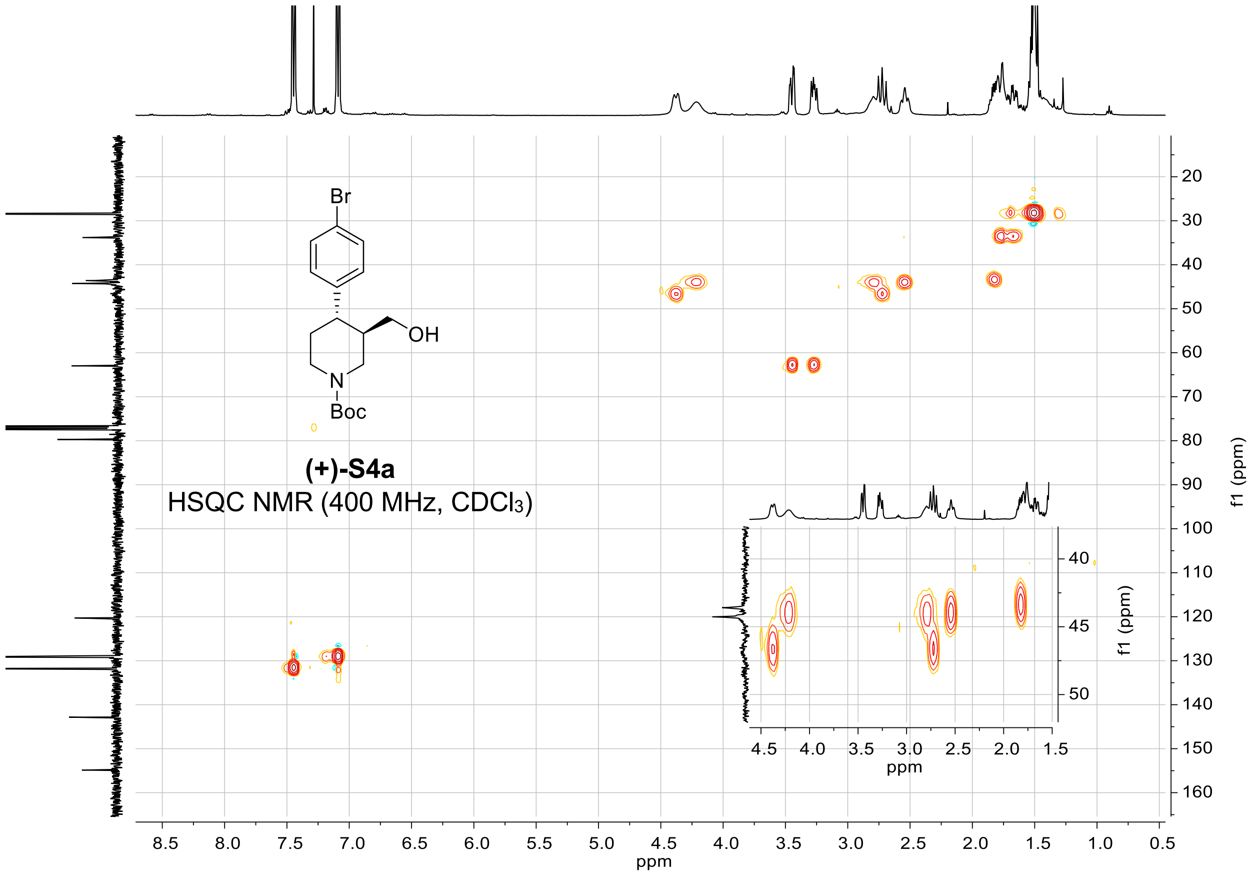
Br
OH
N
Boc
(+)-S4a
HSQC NMR (400 MHz, CDCl3)
ppm
f1 (ppm)

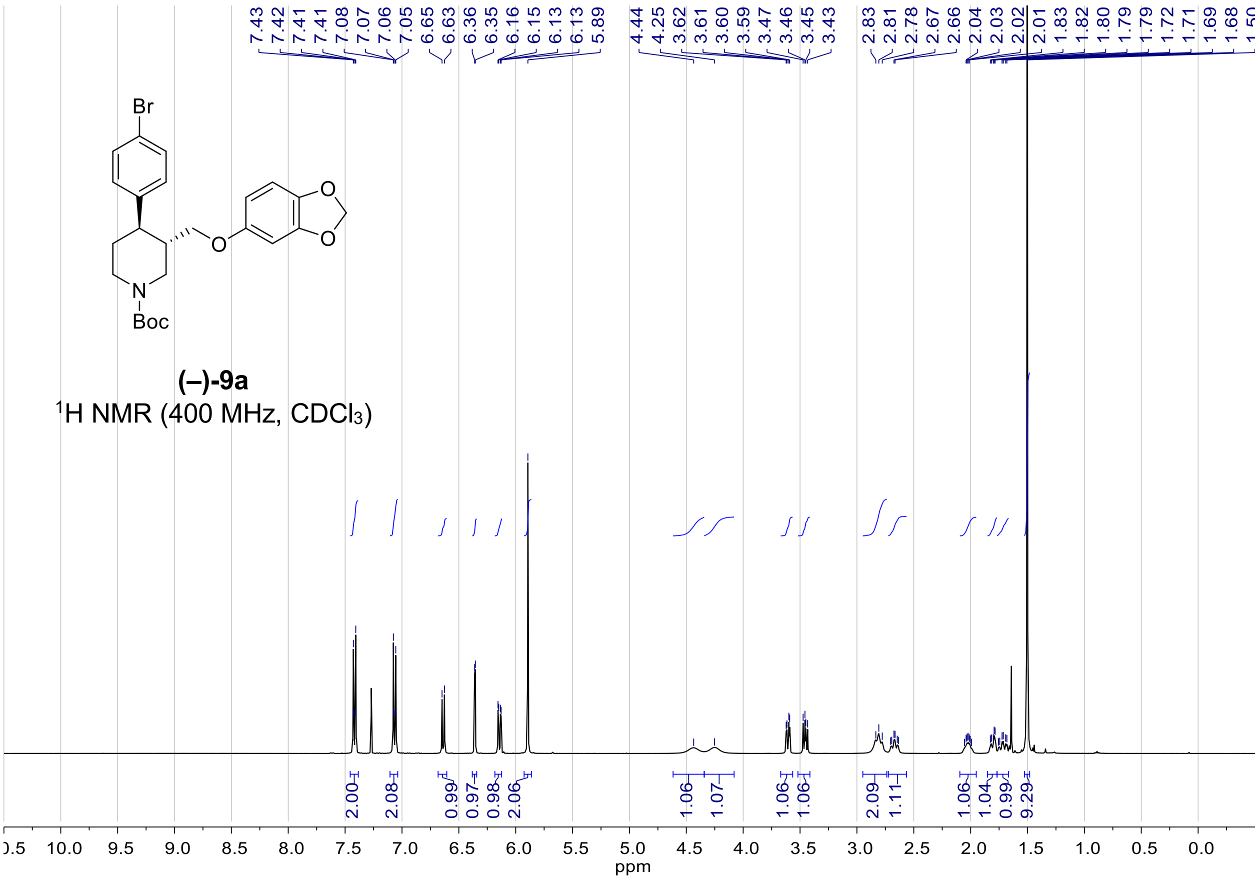
Br
O
O
O
N
Boc
(–)-9a
1H NMR (400 MHz, CDCl3)
7.43 7.42 7.41 7.41 7.08 7.07 7.06 7.05 6.65 6.63 6.36 6.35 6.16 6.15 6.13 6.13 5.89 4.44 4.25 3.62 3.61 3.60 3.59 3.47 3.46 3.45 3.43 2.83 2.81 2.78 2.67 2.66 2.04 2.03 2.02 2.01 1.83 1.82 1.80 1.79 1.79 1.72 1.71 1.69 1.68
2.00 2.08 0.99 0.97 0.98 2.06 1.06 1.07 1.06 1.06 2.09 1.11 1.06 1.04 0.99 9.29
10.0 9.5 9.0 8.5 8.0 7.5 7.0 6.5 6.0 5.5 5.0 4.5 4.0 3.5 3.0 2.5 2.0 1.5 1.0 0.5 0.0
ppm

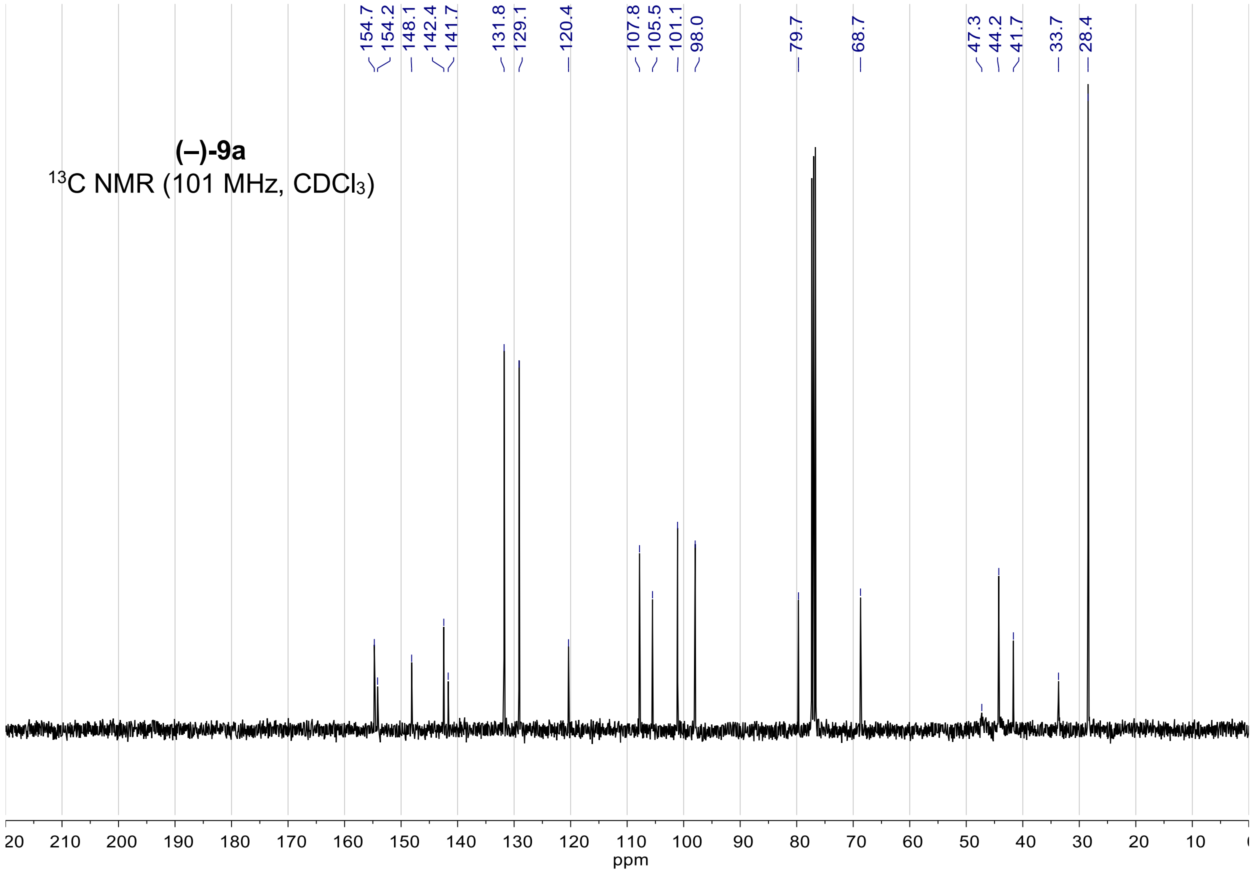
(–)-9a
13C NMR (101 MHz, CDCl3)
154.7 154.2 148.1 142.4 141.7 131.8 129.1 120.4 107.8 105.5 101.1 98.0 79.7 68.7 47.3 44.2 41.7 33.7 28.4
210 200 190 180 170 160 150 140 130 120 110 100 90 80 70 60 50 40 30 20 10
ppm

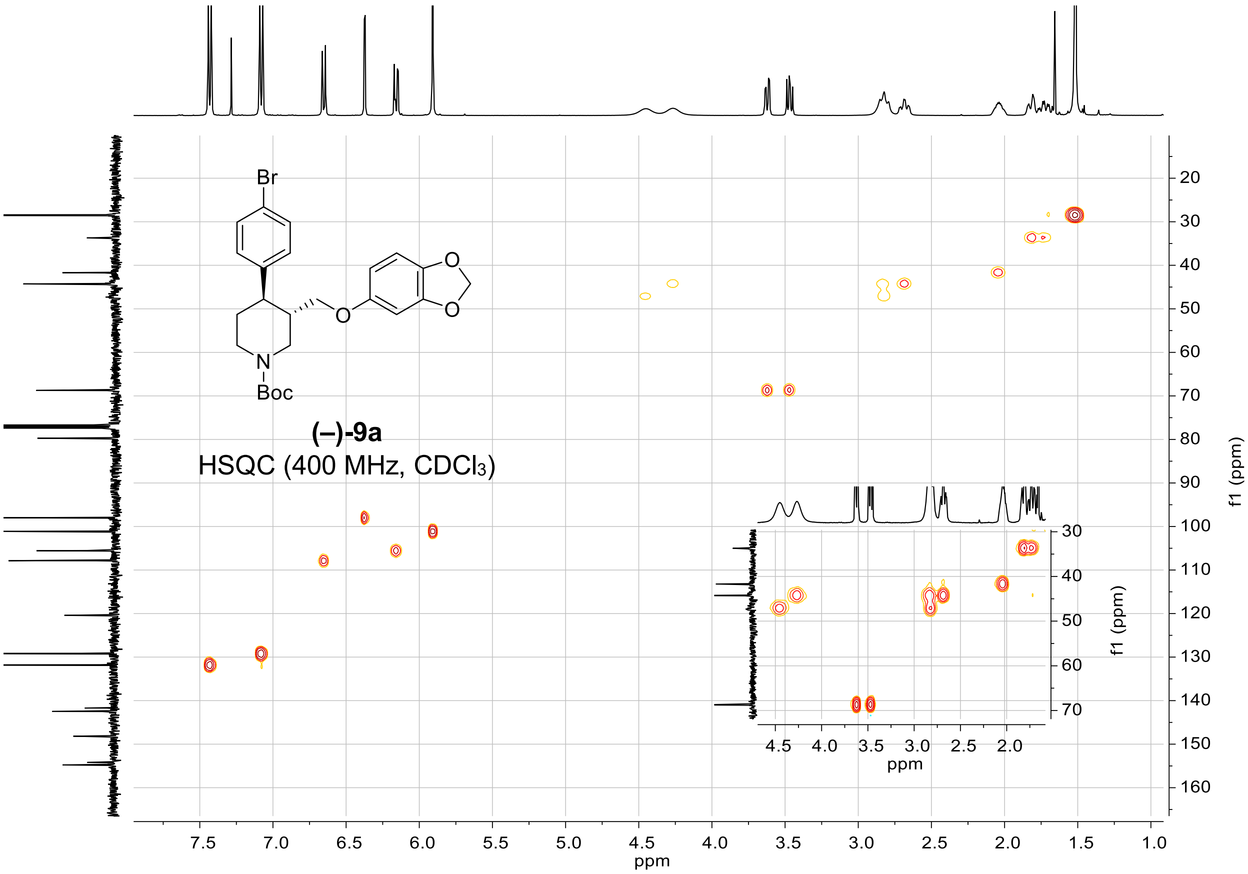
Br
O
O
O
N
Boc
(–)-9a
HSQC (400 MHz, CDCl3)
f1 (ppm)
ppm

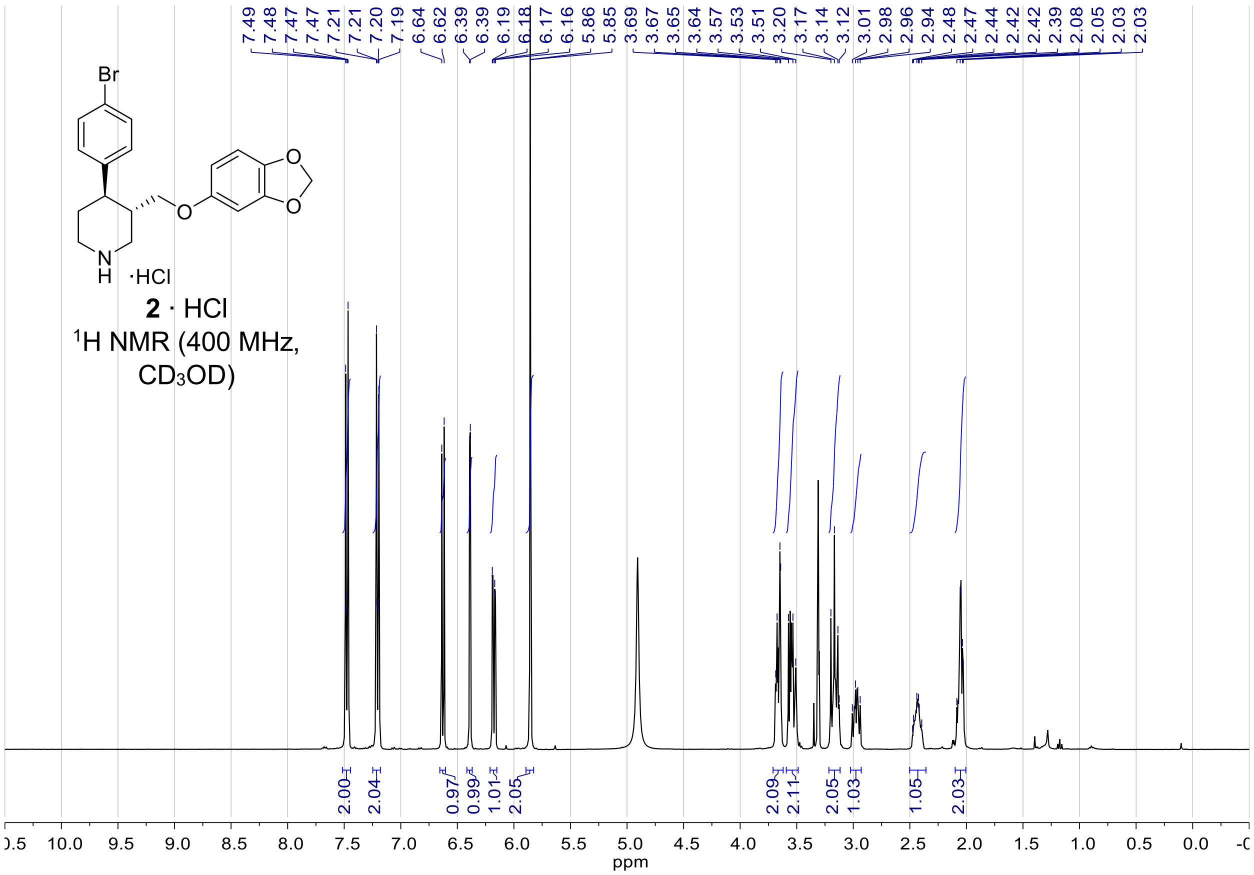

2 · HCl
$^1$H NMR (400 MHz, CD$_3$OD)
ppm


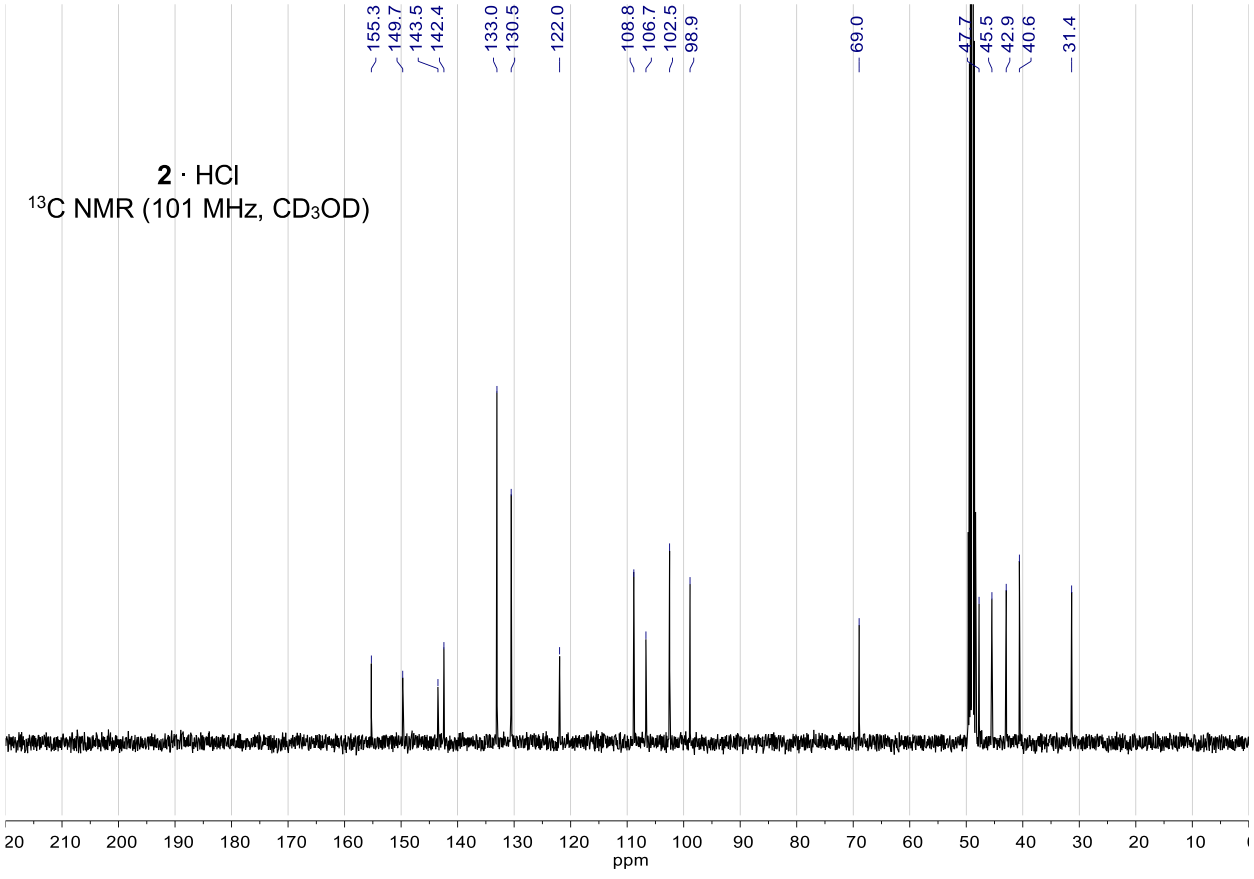

2 · HCl
$^{13}$C NMR (101 MHz, CD$_3$OD)
155.3
149.7
143.5
142.4
133.0
130.5
122.0
108.8
106.7
102.5
98.9
69.0
47.7
45.5
42.9
40.6
31.4
ppm

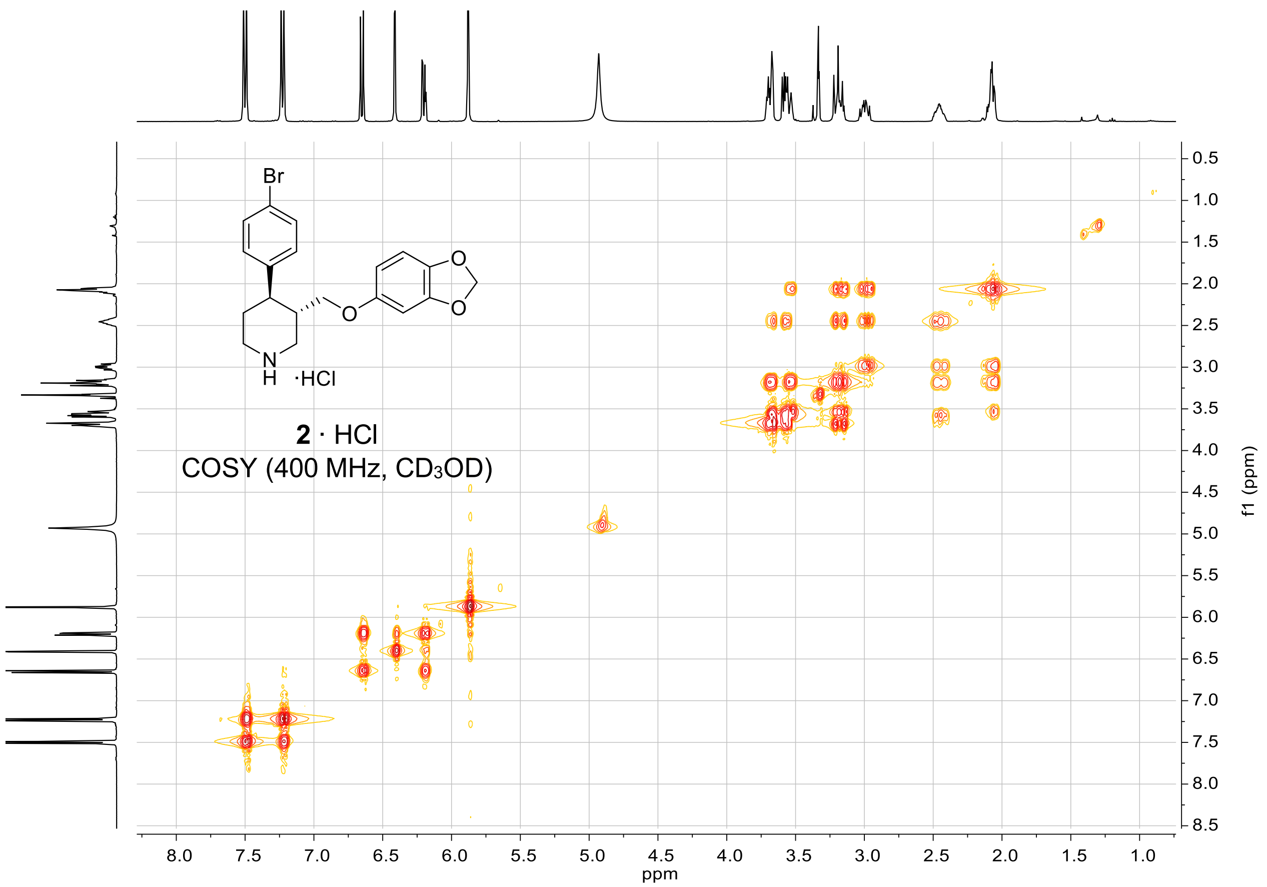
Br
N
H ·HCl
2 · HCl
COSY (400 MHz, CD3OD)
f1 (ppm)
ppm

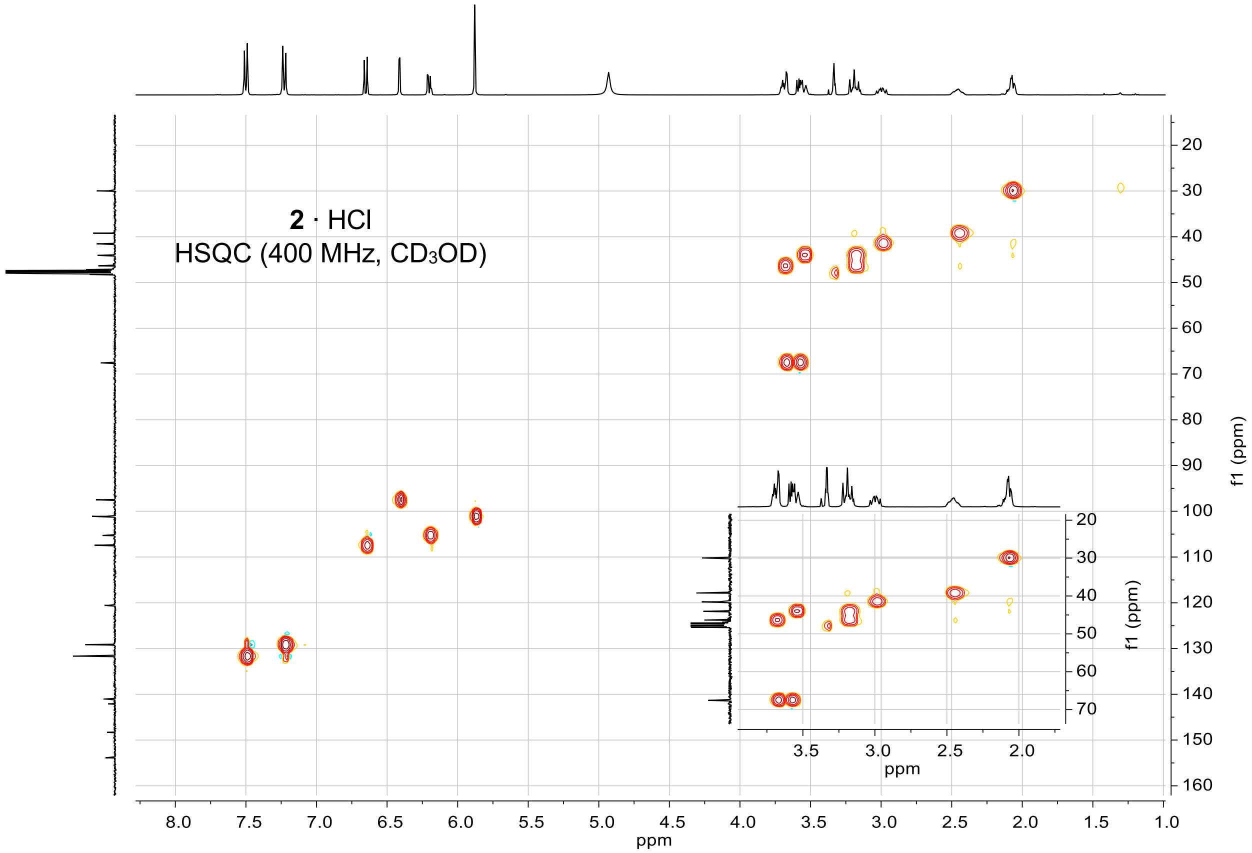
2 · HCl
HSQC (400 MHz, CD3OD)
f1 (ppm)
ppm

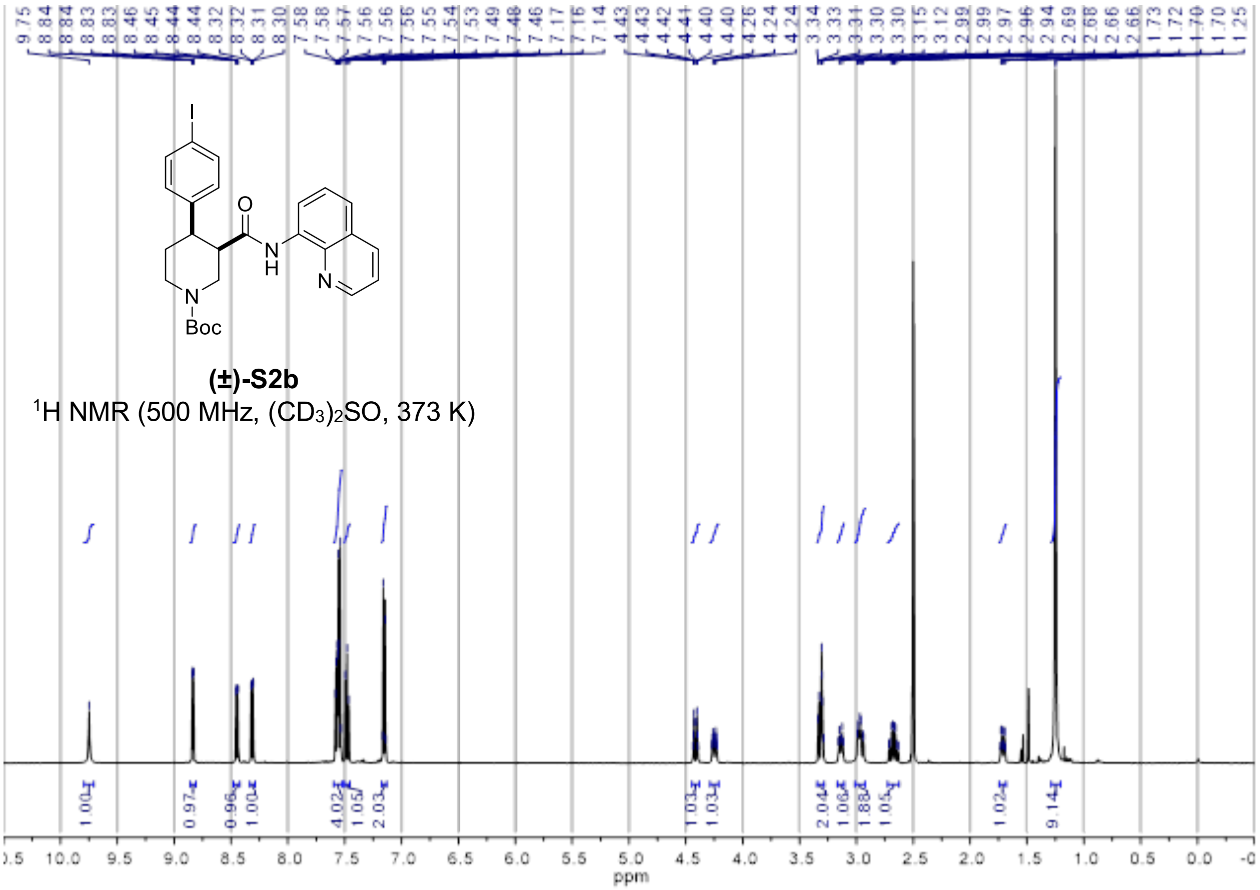


**(±)-S2b**

$^{1}$H NMR (500 MHz, $(CD_3)_2SO$, 373 K)

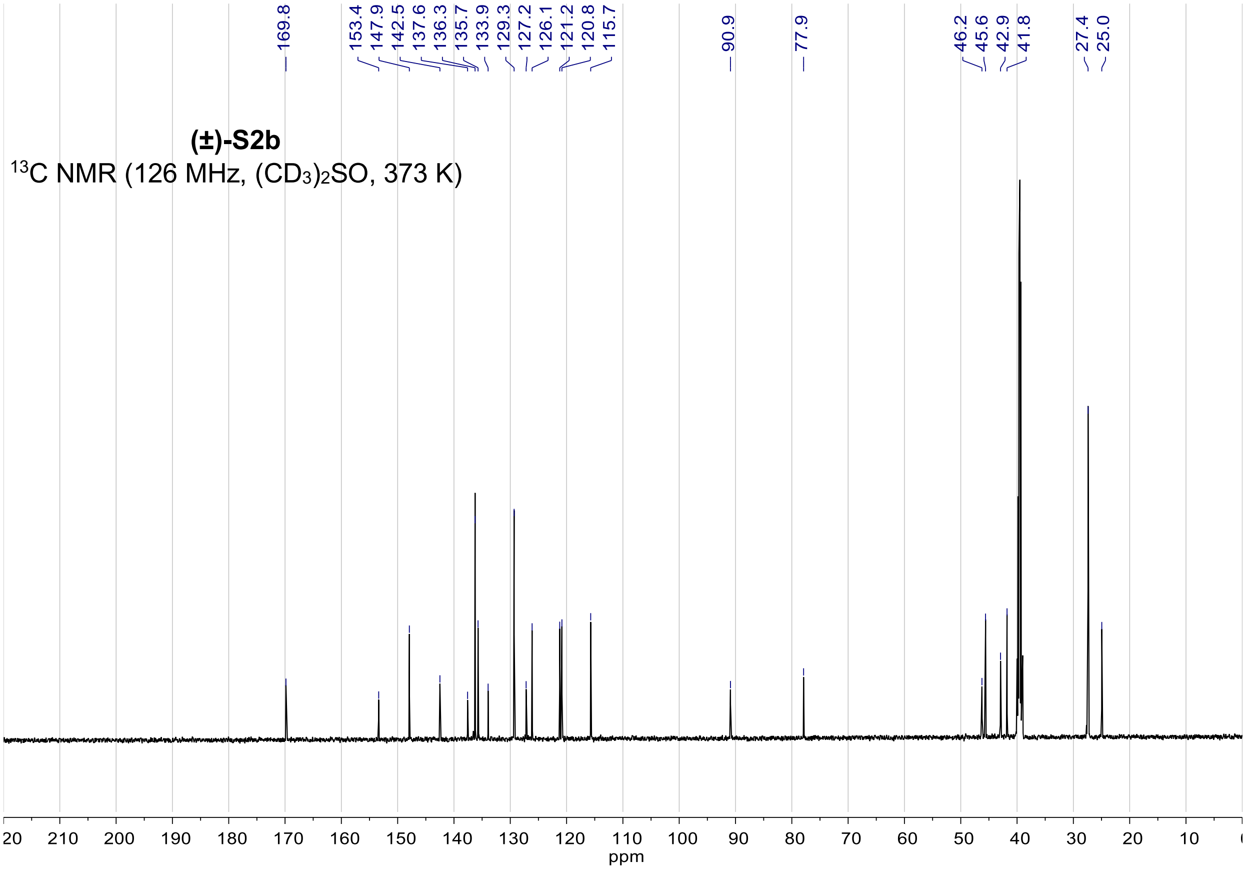


**(±)-S2b**

$^{13}$C NMR (126 MHz, $(CD_3)_2SO$, 373 K)

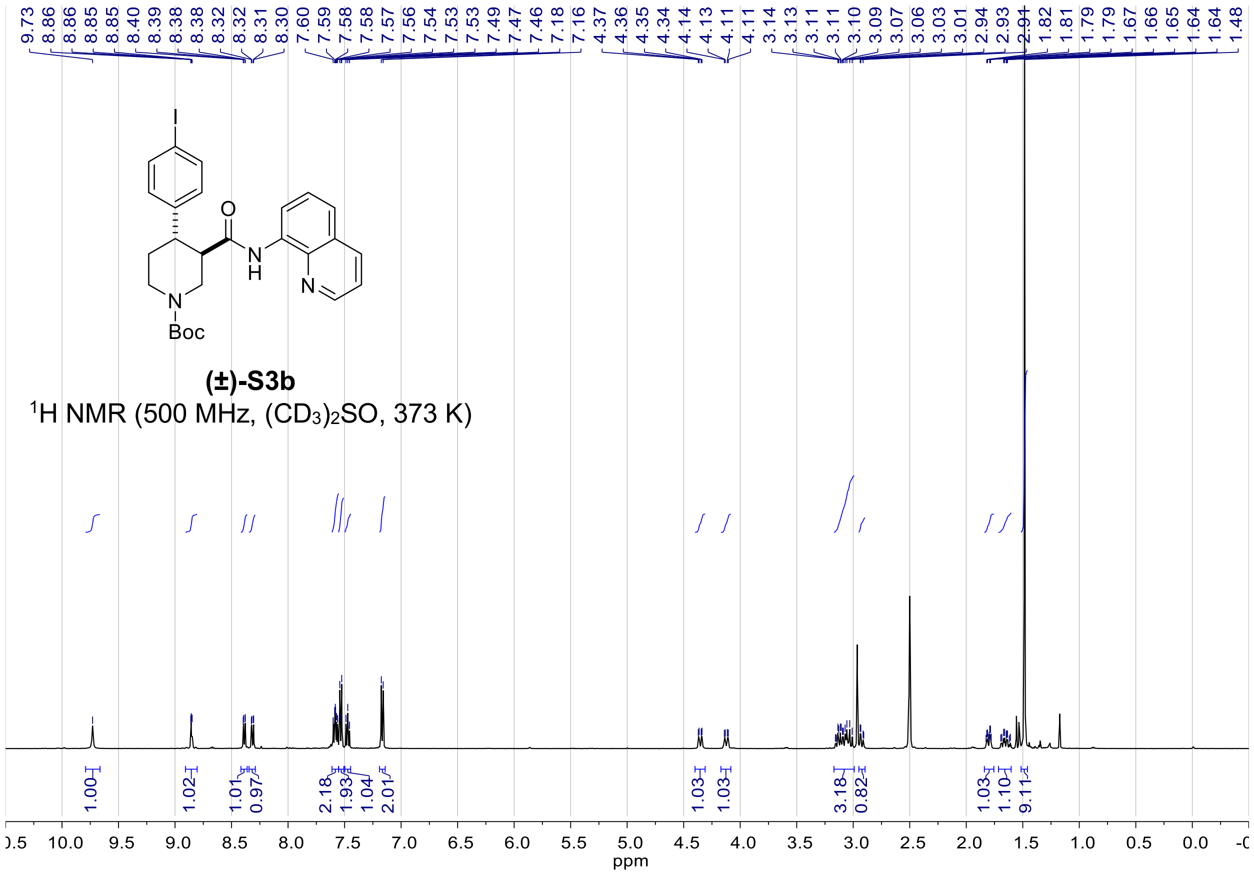
9.73 8.86 8.86 8.85 8.85 8.40 8.39 8.38 8.38 8.32 8.31 8.30 7.60 7.59 7.58 7.58 7.57 7.56 7.54 7.53 7.53 7.49 7.47 7.46 7.18 7.16 4.37 4.36 4.35 4.34 4.14 4.13 4.11 4.11 3.14 3.13 3.11 3.11 3.10 3.09 3.07 3.06 3.03 3.01 2.94 2.93 2.91 1.82 1.81 1.79 1.79 1.67 1.66 1.65 1.64 1.64 1.48
I
O
N
H
N
N
Boc
(±)-S3b
1H NMR (500 MHz, (CD3)2SO, 373 K)
1.00 1.02 1.01 0.97 2.18 1.93 1.04 2.01 1.03 1.03 3.18 0.82 1.03 1.10 9.11
10.0 9.5 9.0 8.5 8.0 7.5 7.0 6.5 6.0 5.5 5.0 4.5 4.0 3.5 3.0 2.5 2.0 1.5 1.0 0.5 0.0
ppm

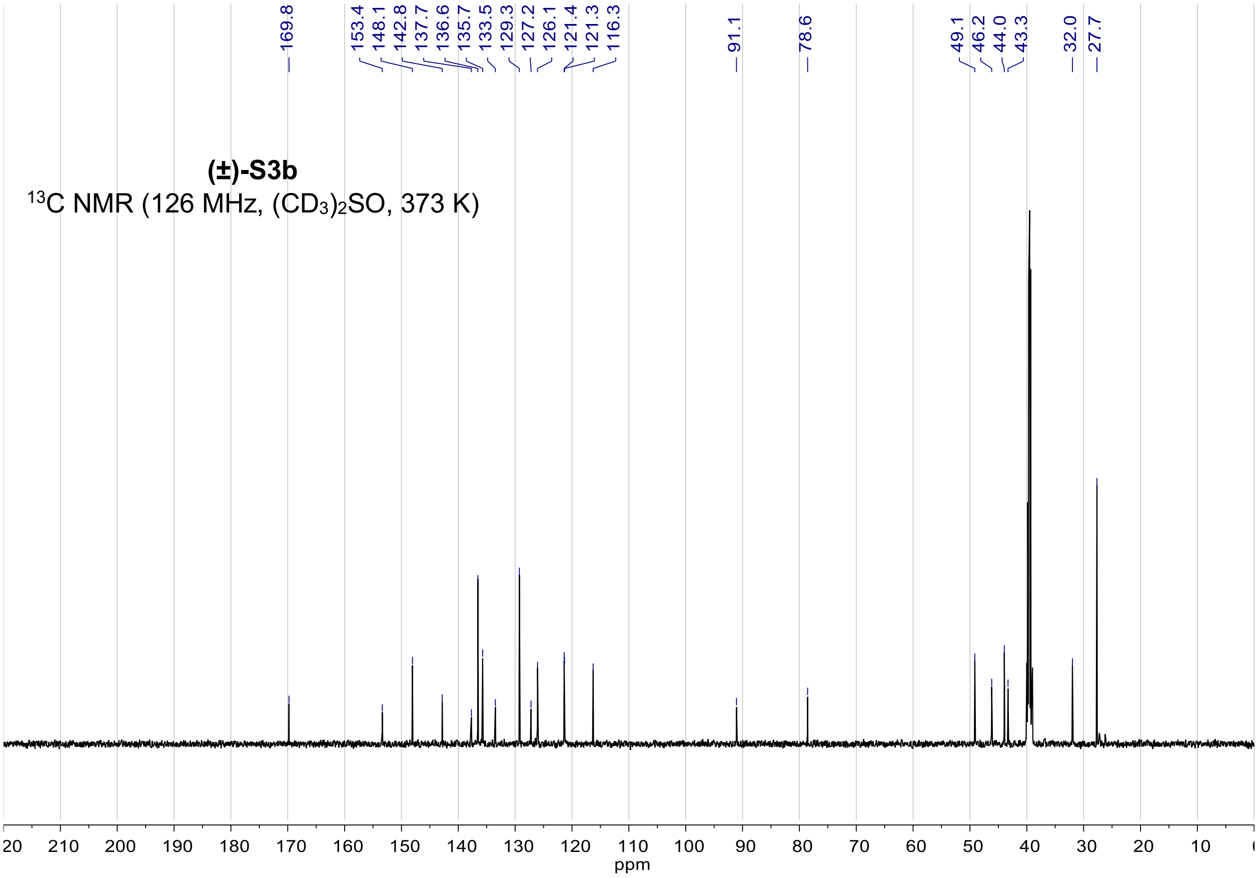
169.8 153.4 148.1 142.8 137.7 136.6 135.7 133.5 129.3 127.2 126.1 121.4 121.3 116.3 91.1 78.6 49.1 46.2 44.0 43.3 32.0 27.7
(±)-S3b
13C NMR (126 MHz, (CD3)2SO, 373 K)
210 200 190 180 170 160 150 140 130 120 110 100 90 80 70 60 50 40 30 20 10
ppm

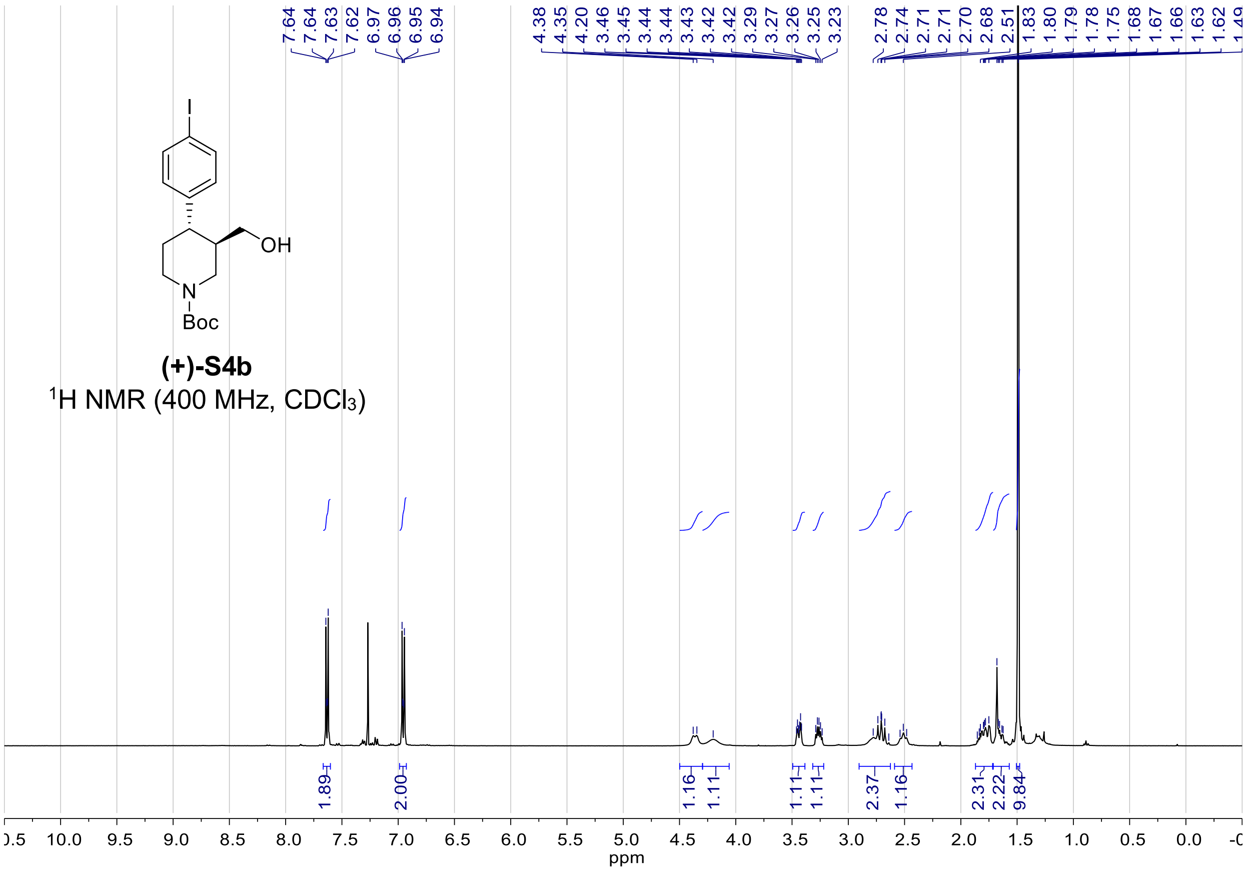

(+)-S4b
$^1$H NMR (400 MHz, CDCl$_3$)
ppm


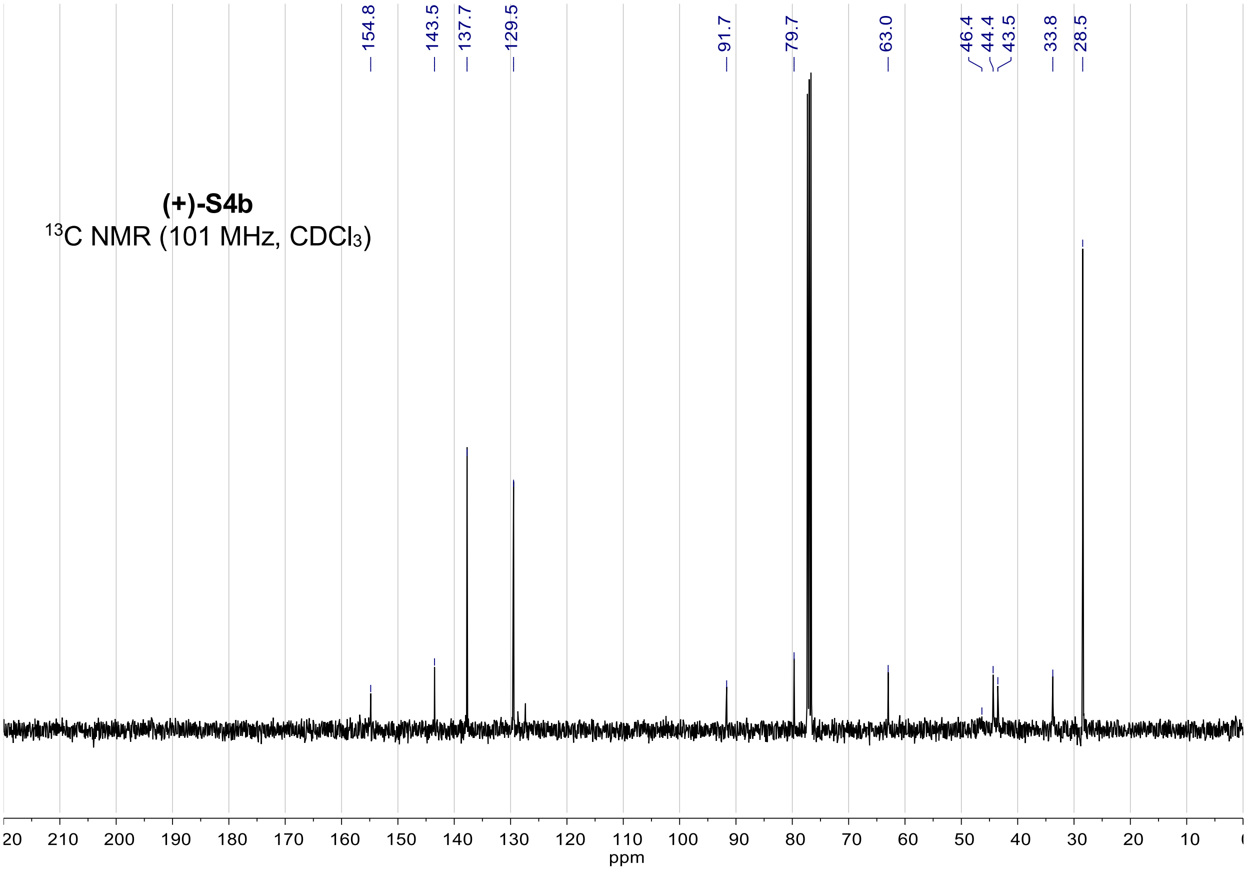

(+)-S4b
$^{13}$C NMR (101 MHz, CDCl$_3$)
154.8
143.5
137.7
129.5
91.7
79.7
63.0
46.4
44.4
43.5
33.8
28.5
ppm

**(–)-9b**

$^{1}$H NMR (400 MHz, $CDCl_3$)

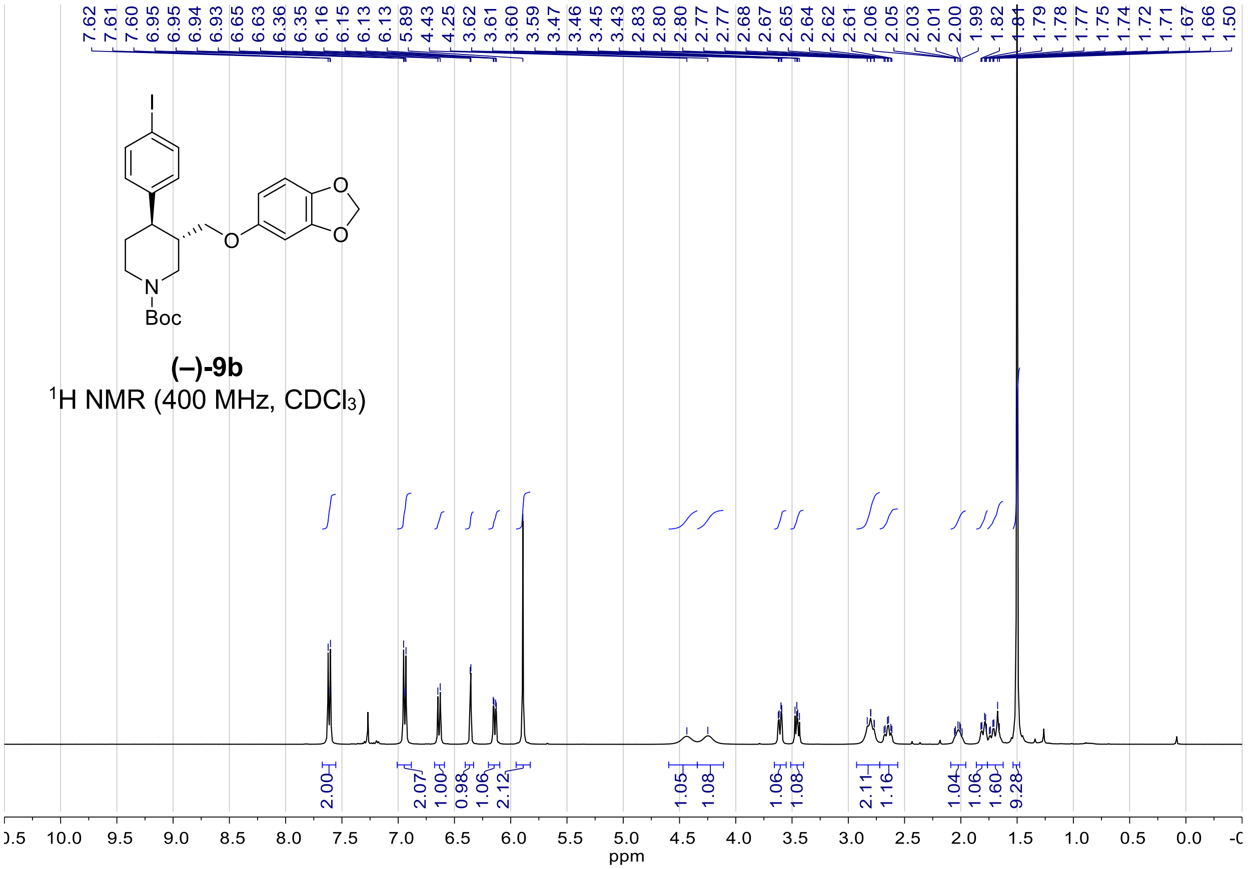


**(–)-9b**

$^{13}$C NMR (101 MHz, $CDCl_3$)

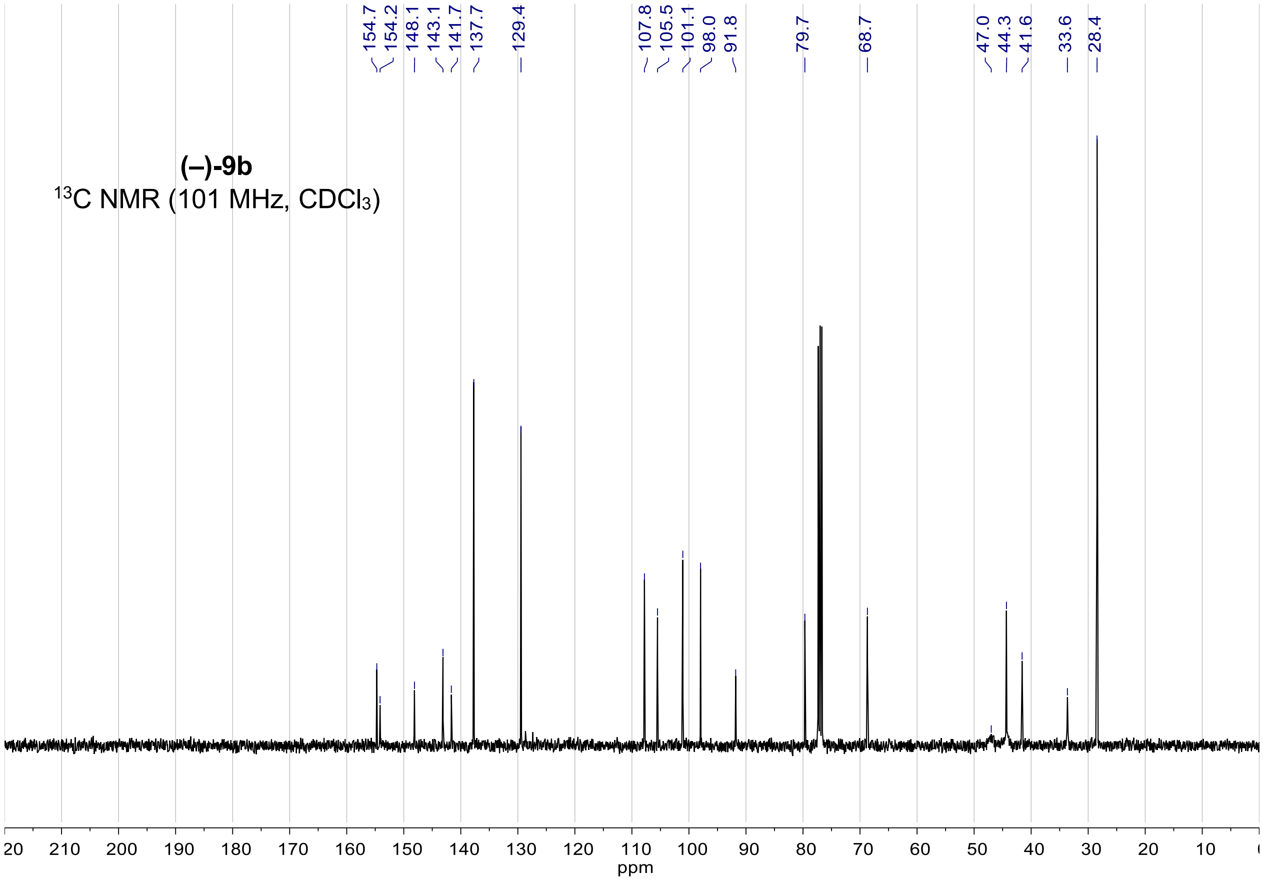

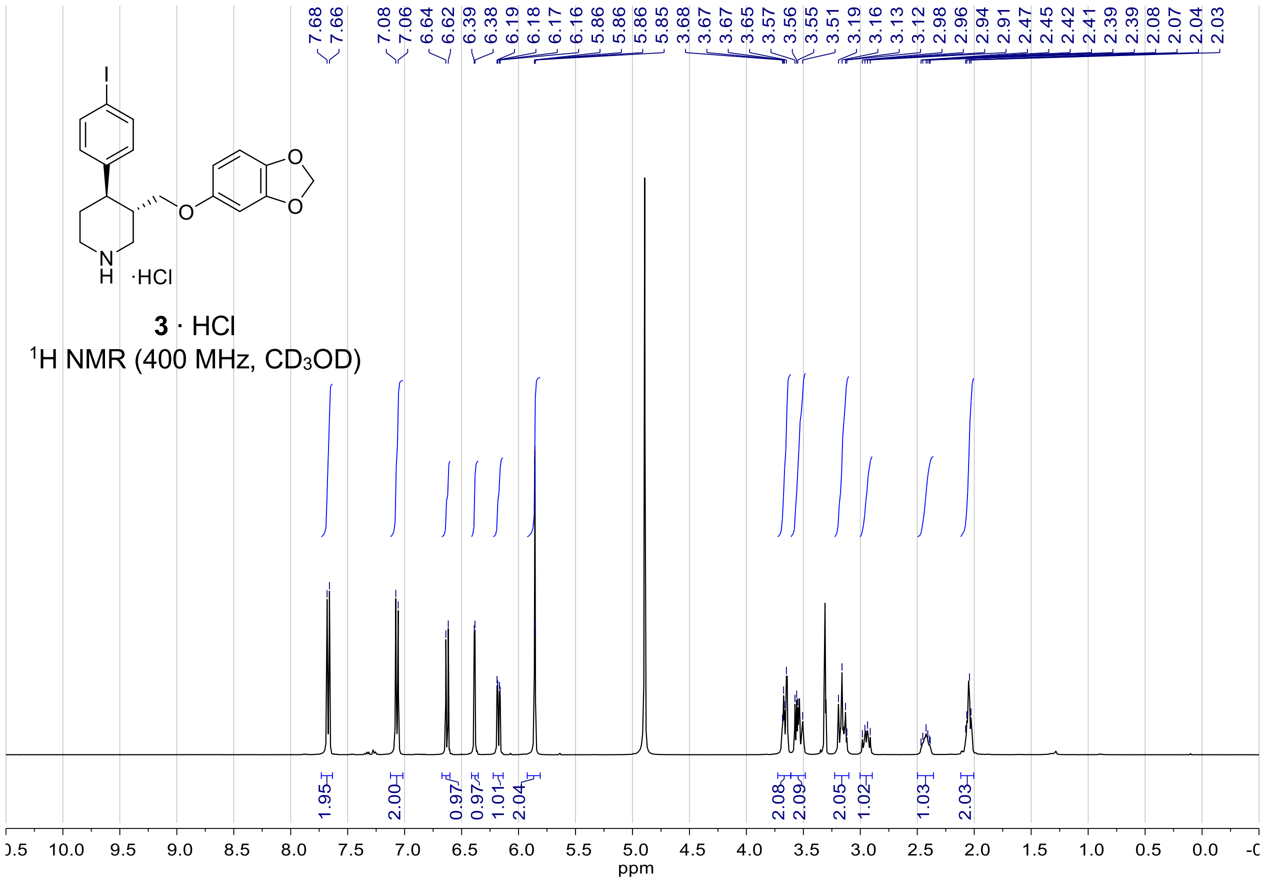
3 · HCl
1H NMR (400 MHz, CD3OD)
7.68
7.66
7.08
7.06
6.64
6.62
6.39
6.38
6.19
6.18
6.17
6.16
5.86
5.86
5.85
3.68
3.67
3.65
3.57
3.56
3.55
3.51
3.19
3.16
3.13
3.12
2.98
2.96
2.94
2.91
2.47
2.45
2.42
2.41
2.39
2.08
2.07
2.04
2.03
1.95
2.00
0.97
0.97
1.01
2.04
2.08
2.09
2.05
1.02
1.03
2.03
ppm

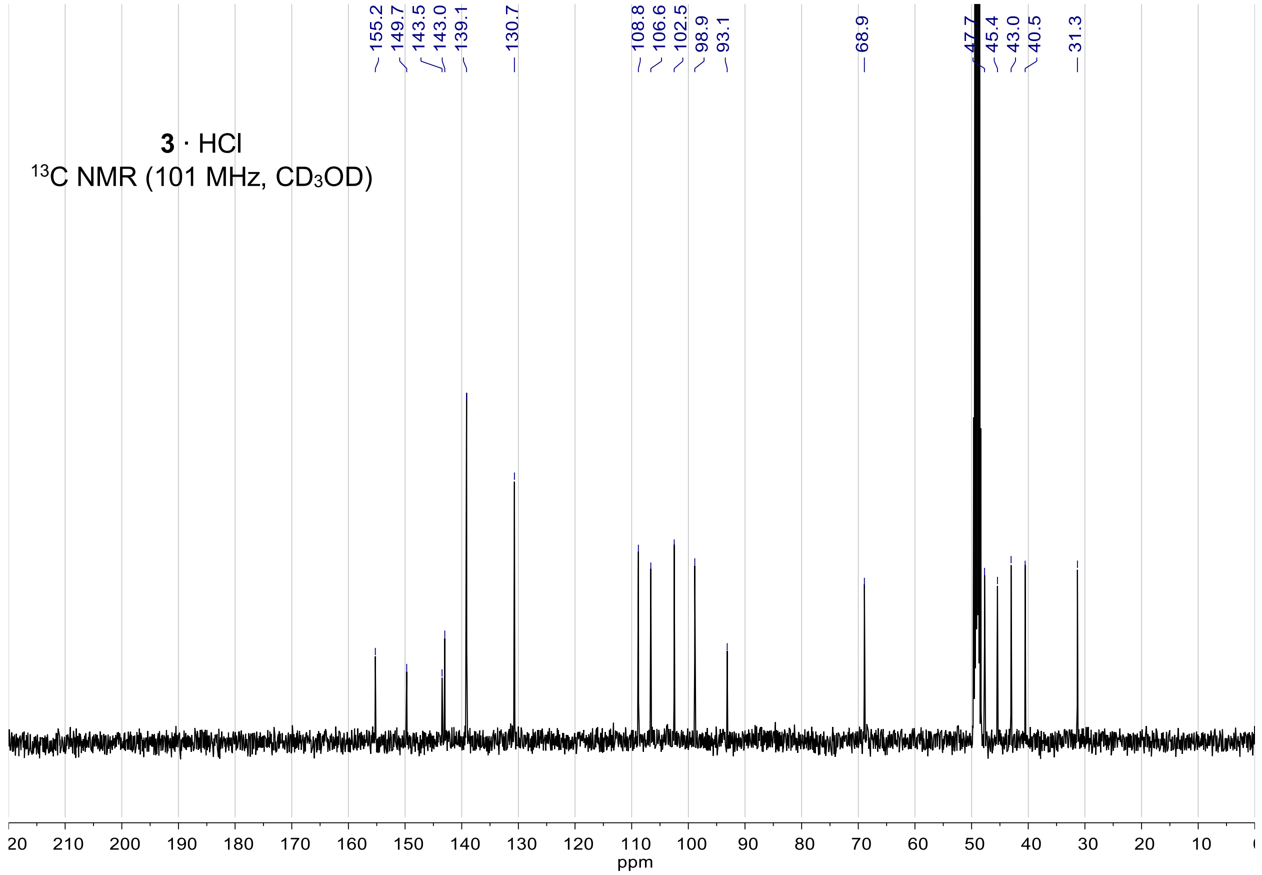
3 · HCl
13C NMR (101 MHz, CD3OD)
155.2
149.7
143.5
143.0
139.1
130.7
108.8
106.6
102.5
98.9
93.1
68.9
47.7
45.4
43.0
40.5
31.3
ppm
